# Genomic landscape of tumor-host interactions with differential prognostic and predictive connotations

**DOI:** 10.1101/546069

**Authors:** Jessica Roelands, Wouter Hendrickx, Peter J.K. Kuppen, Raghvendra Mall, Gabriele Zoppoli, Mohamad Saad, Kyle Halliwill, Giuseppe Curigliano, Darawan Rinchai, Julie Decock, Lucia G Delogu, Tolga Turan, Josue Samayoa, Lotfi Chouchane, Ena Wang, Pascal Finetti, Francois Bertucci, Lance D Miller, Jerome Galon, Francesco M Marincola, Michele Ceccarelli, Davide Bedognetti

**Author notes:** Corresponding authors, Davide Bedognetti, Wouter Hendrickx, Michele Ceccarelli. Equal contribution (Jessica Roelands, Wouter Hendrickx).

## Abstract

An immune active cancer phenotype typified by a T helper 1 (Th-1) immune response has been associated with increased responsiveness to immunotherapy and favorable prognosis in some but not all cancer types. The reason of this differential prognostic connotation remains unknown. Through a multi-modal Pan-cancer analysis among 31 different histologies (9,282 patients), we demonstrated that the favorable prognostic connotation conferred by the presence of a Th-1 immune response was abolished in tumors displaying specific tumor-cell intrinsic attributes such as high TGF-ß signaling and low proliferation capacity. This observation was validated in the context of immune-checkpoint inhibition. WNT-ß catenin, barrier molecules, Notch, hedgehog, mismatch repair, telomerase activity, and AMPK signaling were the pathways most coherently associated with an immune silent phenotype together with mutations of driver genes including *IDH1/2, FOXA2, HDAC3, PSIP1, MAP3K1, KRAS, NRAS, EGFR, FGFR3, WNT5A*, and *IRF7*. Our findings could be used to prioritize hierarchically relevant targets for combination therapies and to refine stratification algorithms.

## Introduction

Evidence of the effects of anti-tumoral immunity on cancer progression has accumulated over the last decades. The identification of tumor immune escape mechanisms, most importantly the characterization of immune checkpoints, led to major advances in immunotherapy. Immune checkpoint inhibitors have dramatically improved clinical outcome for a subset of patients across multiple cancer types. Despite this progress, the majority of patients (60-80%) still fail to respond (Emens et al., 2017; Gong et al., 2018). Understanding the relationship between tumor cell and the immune system is critical to develop more effective therapeutic strategies.

A pre-existing intratumoral anti-tumor immune response has been associated with favorable outcome and responsiveness to immunotherapy (Galon et al., 2013). However, multiple studies have reported differences in the association between measures of intratumoral immune activity and survival across different cancer types (Charoentong et al., 2017; Danaher et al., 2018; Tamborero et al., 2018; Thorsson et al., 2018; Varn et al., 2017). In breast cancer, a positive association between survival and density of tumor infiltrating lymphocytes, as estimated by transcriptomic data, was restricted to tumors displaying a high mutational load or an aggressive/high proliferative phenotype (Miller et al., 2016; Nagalla et al., 2013; Thomas et al., 2018). Proposed transcriptome-based immunological classifications range from a measure of cytolytic activity by mean expression of *GZMA* and *PRF1* genes (Rooney et al., 2015), to reflections of immune cell infiltration by cell-specific transcriptional profiles (Bindea et al., 2013; Nagalla et al., 2013), or gene signatures reflecting molecular components of an active antitumor immune response, including Major Histocompatibility Complex (MHC), co-stimulatory or immunomodulatory molecules (Ayers et al., 2017; Charoentong et al., 2017; Wang et al., 2008). Reported prognostic and predictive signatures typically show overlapping genes or genes involved in conserved immunologic processes (Bedognetti et al., 2016, 2013; Galon et al., 2013; Wang et al., 2013b, 2013a). We termed these mechanisms as the Immunologic Constant of Rejection (ICR) (Galon et al., 2013; Wang et al., 2008). The ICR signature incorporates IFN-stimulated genes driven by transcription factors *IRF1* and *STAT1, CCR5* and *CXCR3* ligands, immune effector molecules, and counter-activated immune regulatory genes (Hendrickx et al., 2017; Turan et al., 2018; Wang et al., 2013b, 2008). Overall, the high expression of ICR typifies “hot”/immune active tumors characterized by the presence of a T helper 1 (Th-1)/cytotoxic immune response, as described in detail elsewhere (Bertucci et al., 2018; Galon et al., 2013; Hendrickx et al., 2017; Turan et al., 2018).

Previously, we observed a significantly prolonged survival of patients with tumors displaying a coordinated expression of ICR genes in breast cancer (Bertucci et al., 2018; Hendrickx et al., 2017). Moreover, we identified genetic determinants of different immune phenotypes (Hendrickx et al., 2017). In particular, we reported that transcriptional dysregulation of the MAPK pathways sustained by genetic alterations (i.e., MAP3K1 and MAP2K4 mutations) are enriched in immune silent tumors (Hendrickx et al., 2017). We also observed that the ICR signature refines and improves the prognostic value of conventional prognostic signatures adopted in breast cancer (Bertucci et al., 2018). Here, we propose a systematic analysis of the entire TCGA cohort encompassing 31 different histologies. Using a pan-cancer approach, we identified novel relationships between tumor genetic programs and immune orientation. After having demonstrated differential associations between ICR classification and overall survival across cancer types, we systemically analyzed in which (molecular) contexts ICR has prognostic value and in which ones it does not. Combination of immune orientation with tumor intrinsic attributes that interact with its prognostic significance could refine tumor immunologic classifications. This approach was validated in the context of immune-checkpoint inhibition allowing better predictive precision.

## Results

### Prognostic impact of ICR classification is different between cancer types

RNA-seq data of samples from a total of 9,282 patients across 31 distinct solid cancer types were obtained from TCGA. To classify cancer samples based on their immune orientation, we performed unsupervised consensus clustering for each cancer type separately based on the expression of the ICR immune gene signature. This signature consists of 20 genes that reflect activation of Th1-signaling (*IFNG, TXB21, CD8B, CD8A, IL12B, STAT1*, and *IRF1*), CXCR3/CCR5 chemokine ligands (*CXCL9, CXCL10*, and *CCL5*), cytotoxic effector molecules (*GNLY, PRF1, GZMA, GZMB*, and *GZMH*) and compensatory immune regulators (*CD274/PD-L1, PDCD1, CTLA4, FOXP3*, and *IDO1*) (**Figure 1A**) (Bedognetti et al., 2016; Galon et al., 2013; Hendrickx et al., 2017; Turan et al., 2018). Expression of these genes showed a positive correlation with each other across all cancer types (**Supplementary Figure 1**). The ICR signature highly correlates with other immune signatures that aim to reflect a highly active immune tumor microenvironment, including the Tumor Inflammation Signatures (TIS) (r = 0.97)(Danaher et al., 2018) (**Supplementary Figure 2**). As a representative example, consensus clustering and cluster assignment of skin cutaneous melanoma (SKCM) is shown in **Figure 1A**. Analogous figures for each of the 31 cancer types are available as cancer datasheets at figshare.com.

**Figure 1:**
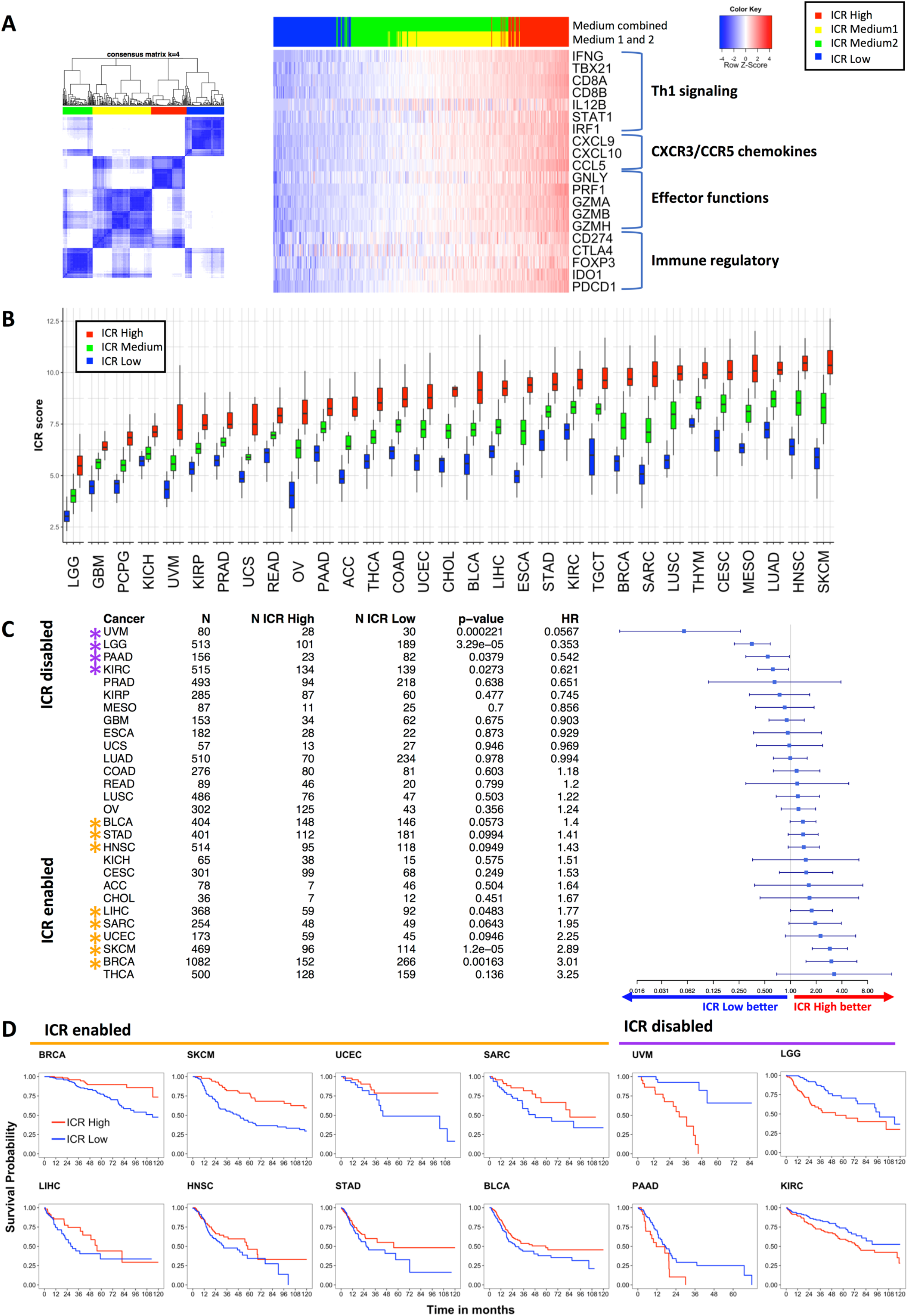
Immunologic classification of 31 cancer types based on expression of ICR gene signature. **A.** Consensus cluster matrix of SKCM samples based on RNA-seq expression values of ICR genes (left panel). RNA-seq expression heatmap of ICR genes annotated with ICR consensus clusters (n = 469). Clusters with intermediate ICR gene expression levels (ICR Medium1 and ICR Medium2) were combined to obtain ICR High, Medium and Low groups (HML classification). ICR genes reflect 4 components of immune mediated tissue rejection: Th1 signaling, CXCR3/CCR5 chemokines, immune effectors and immune regulatory functions (right panel). **B.** Boxplot of ICR scores across ICR clusters in 31 cancer types. Cancer types are ordered by mean ICR score per cancer. **C.** Forest plot showing HRs (overall survival) of ICR Low versus High, p-value and number of patients (N) for each of the cancer types. ICR-enabled cancer types (HR > 1; p < 0.1) are indicated with orange asterisks and ICR-disabled cancer types (HR < 1; p > 0.1) are indicated with purple asterisks. Cancer types PCPG, THYM and TGCT are excluded from the plot, because confidence intervals ranged from 0 to infinite due to low number of deaths in these cancer types. **D.** Kaplan Meier curves showing OS across two three different ICR groups in ICR-enabled and ICR-disabled cancer types. (Figures of panel A and Kaplan Meier curves for each individual cancer type are available in the <u>cancer datasheets</u>).

As shown in **Figure 1B**, the mean expression of ICR genes, or ICR score, varies between cancer types, reflecting general differences in tumor immunogenicity between cancers. While brain tumors (brain lower grade glioma’s (LGG) and glioblastoma multiforme (GBM)) typically display low immunological signals (McGranahan et al., 2017), skin cutaneous melanoma (SKCM) and head and neck squamous cell carcinoma (HNSC) display high levels of immune activation (Economopoulou et al., 2016; Passarelli et al., 2017; Thorsson et al., 2018). In addition, the distribution of ICR scores among patients and the difference between the highest and lowest ICR scores varies between cancers. Accordingly, the proportions of patients assigned to specific ICR clusters are dependent on the cancer type. Even more clinically relevant, the relation of the different immune phenotypes to survival is dissimilar among cancer types (**Figure 1C-D**). While the ICR High phenotype (hot) shows a significant survival benefit compared with the ICR Low phenotype (cold) for various cancer types (BRCA, SKCM, UCEC, SARC), the ICR High cluster is associated with significantly reduced overall survival in other cancer types (UVM, LGG, PAAD, KIRC) (**Figure 1C**). Similar results were obtained when Cox regression analysis was performed on ICR score as a continuous variable (**Supplementary Table 1**). To explore biological differences between cancer types in which a highly active immune phenotype is mostly associated with favorable survival and cancer types in which this phenotype is mostly associated with decreased survival, we categorized cancer types in *ICR-enabled* (BRCA, SKCM, UCEC, SARC, LIHC, HNSC, STAD, BLCA) and ICR-*disabled* (UVM, LGG, PAAD, KIRC) groups, respectively (**Figure 1C**). All other cancer types in which ICR did not show an association or trend were categorized as *ICR-neutral*. Of important note, this classification was used for explorative purposes, a role of the immune mediated tumor rejection cannot be precluded in ICR-neutral cancer types.

First, we explored whether the ICR scores and their distributions were different among these defined groups of cancer types. Mean ICR score is low for most ICR-disabled (ranging from 3.97 to 8.34) compared to ICR-enabled cancer types (ranging from 7.26 to 8.36) (**Supplementary Figure 3A**). This observation is most noticeable for ICR-disabled cancer types LGG and UVM. Moreover, the difference (delta) between ICR scores in ICR High compared to ICR Low groups is higher in ICR-enabled cancer types (range: 2.98-4.97) compared to ICR-neutral (range: 1.48-4.49) and ICR-disabled cancer types (range: 2.29-3.35) (**Supplementary Figure 3B**). These factors could underlie, at least partially, the observed divergent associations with survival.

To define whether tumor pathologic stage might interact with the association between ICR and overall survival (OS), we fitted a Cox proportional hazards model for each group of ICR-enabled, ICR-neutral and ICR-disabled cancer types (**Table 1**). Overall, including ICR High and ICR Low samples from all cancer types, ICR has significant prognostic value independent of AJCC pathologic stage. For ICR-enabled cancer types, the ICR High group also remains significantly associated with improved survival after adjusting for tumor pathologic stage. For ICR-disabled cancer types, ICR High was associated with worse survival in univariate analysis (HR <1). However, in multivariate models this negative prognostic value of ICR was lost (HR=1.054; 95% CI= 0.7702-1.443). Kaplan-Meier plots stratified by pathologic stage showed that within individual pathologic stages, ICR was not associated with OS for ICR-disabled cancers (**Supplementary Figure 4.1**). In fact, in the ICR-disabled tumors (but not in the ICR-enabled ones), ICR was significantly higher (p = 10 e-7) in advanced vs early stages (**Supplementary Figure 4.2**). Similarly, a progressive enrichment of ICR high samples was observed with more advanced stages in the ICR-disables tumors UVM and KIRC, and, in, LGG with more advanced grades.

**Table 1.** Cox proportional hazards regression for association with overall survival in ICR-enabled and ICR-disabled tumors: ICR High and ICR Low samples included (ICR Medium samples excluded).

| Variables | Univariable |  | Multivariable |  |
| --- | --- | --- | --- | --- |
|  | HR (95% CI) | p | HR (95% CI) | p |
| *ICR overall (n = 4735)* |  |  |  |  |
| ~ ICR cluster (ICR Low vs. High) | 1.203 (1.081-1.339) | 0.00073 *** | 1.343 (1.180- 1.528) | 7.85e-06 *** |
| ~ Pathologic stage (Stage I, II, III, IV) | 1.72 (1.615-1.832) | <2e-16 *** | 1.716 (1.611- 1.827) | <2e-16 *** |
| *Samples from ICR-enabled cancer types (n = 1742)* |  |  |  |  |
| ~ ICR cluster (ICR Low vs. High) | 1.631(1.374-1.937) | 2.26e-8 *** | 1.488 (1.233- 1.795) | 3.35e-05 *** |
| ~ Pathologic stage (Stage I, II, III, IV) | 1.817 (1.644-2.008) | <2e-16 *** | 1.798 (1.628- 1.987) | <2e-16 *** |
| *Samples from ICR-disabled cancer types (n = 721)* |  |  |  |  |
| ~ ICR cluster (ICR Low vs. High) | 0.6194 (0.4801-0.7992) | 0.000229 *** | 1.054 (0.7702- 1.443) | 0.742 |
| ~ Pathologic stage (Stage I, II, III, IV) | 1.55 (1.351-1.778) | 4.22e-10 *** | 1.560 (1.3520- 1.801) | 1.19e-9 *** |
| *Samples from ICR neutral cancer types (n = 2272)* |  |  |  |  |
| ~ ICR cluster (ICR Low vs. High) | 1.160 (0.983-1.369) | 0.0789 | 1.336 (1.065- 1.676) | 0.0122 * |
| ~ Pathologic stage (Stage I, II, III, IV) | 1.665 (1.5-1.848) | <2e-16 *** | 1.640 (1.477- 1.821) | <2e-16 *** |

Signif. codes: \*\*\* <0.001; \*\* <0.01; \* <0.05
ICR cluster entered as categorical (factor) variable (factor levels: "ICR High", "ICR Low") Pathologic stage as semi-continuous variable (Stage I = 1; Stage II = 2; Stage III = 3; Stage IV = 4) LGG and GBM were not included, as tumor stage is not available (not applicable) for these cancer types.

For ICR-neutral cancer types, while ICR was not associated with survival in univariate analysis, multivariate analysis indeed identified a positive prognostic value of the ICR classification, though less robust than observed for ICR-enabled cancer types.

### ICR reflects anti-tumor immune activity and is inversely correlated with tumor-related pathways associated with immune escape

To further explore differences between cancer types, we aimed to compare the density of leukocyte subpopulations between ICR High and Low samples across cancers. Gene expression signatures specific to 24 cell types (Bindea et al., 2013) were used to deconvolute the abundance of immune cells in tumor samples by performing single sample gene set enrichment analysis (ssGSEA) (Barbie et al., 2009). Cell-specific enrichment scores (ES) for each patient demonstrated a clear enrichment of transcripts specific to T- and B cells in ICR High patients (**Figure 2A**). More specifically, ICR High samples showed increased expression of transcripts associated with cytotoxic T cells, T-regulatory (T-reg) cells, Th1 cells, NK CD56dim cells, activated dendritic cells (aDC) and macrophages, compared with ICR Medium and ICR Low samples. This observation is consistent across cancer types, in both ICR-enabled and ICR-disabled cancers. So, in addition to the immune functional molecular orientation, the ICR gene signature is a good reflection of anti-tumor immune cell infiltration (Lu et al., 2017). To quantitatively compare immune cell enrichment between individual cancer types, the mean ES was calculated for each cancer type (**Supplementary Figure 5**). Overall, no single consistent difference in terms of immune cell enrichment can be observed that can discriminate ICR-enabled from ICR-disabled cancer types. LGG and UVM show an overall low immune infiltration, consistent with our reported low ICR scores.

**Figure 2:**
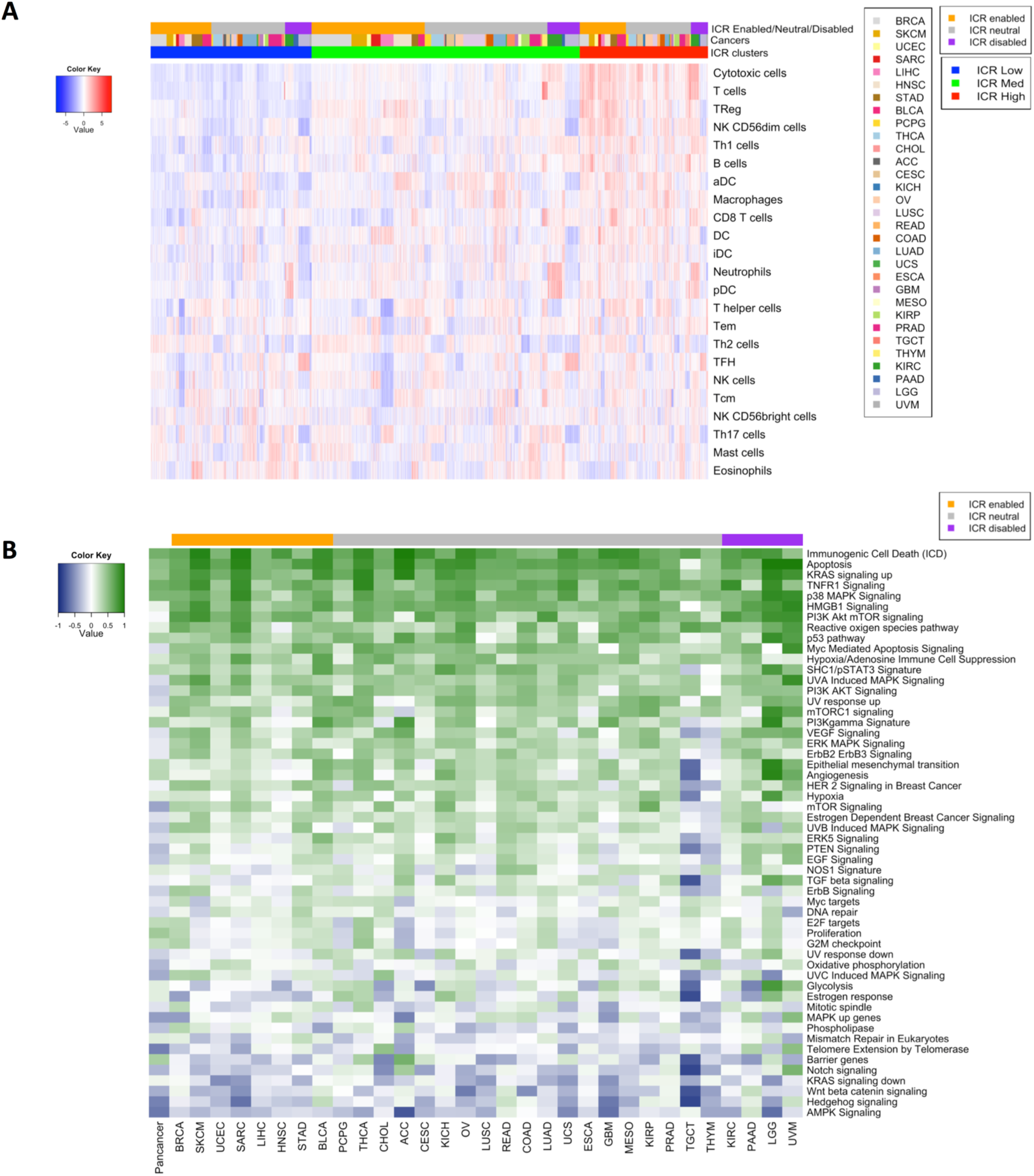
Deconvolution of immune cell populations and enrichment of oncogenic pathways through single sample GSEA. **A.** Heatmap of enrichment values for cell-specific immune-signatures as described by Bindea *et al*. Samples are ordered by ICR cluster and ordered by cancer type within ICR clusters. **B.** Pearson coefficient of correlation between ICR score and enrichment scores of oncogenic pathways per cancer. Pathways that have a positive correlation with ICR are green and those with an inverse correlation are blue.

We then proceeded to examine which tumor intrinsic attributes correlate with immune phenotype as reflected by ICR gene expression. We performed ssGSEA to identify enrichment of transcripts of common tumor-related pathways (Hendrickx et al., 2017; Lu et al., 2017; Salerno et al., 2016). Not surprisingly, immune-related pathways including TNFR1 Signaling and immunogenic cell death showed a strong positive correlation with expression of ICR genes (**Figure 2B**). This implies that our immune signature captures the anti-tumoral immunological processes well across a wide range of cancer types. Interestingly, few pathways were identified that inversely correlated with ICR gene expression, potentially representing mechanisms by which immune silent tumors develop their phenotype. These pathways include WNT-β catenin (Corrales et al., 2017; Spranger and Gajewski, 2015), barrier genes (Salerno et al., 2016), AMPK signaling (Dandapani and Hardie, 2013), mismatch repair, telomerase extension by telomerase, Notch, and Hedgehog, signaling pathways. Of special note, genes that we previously found to be upregulated in MAP3K1/MAP2K4-mutated vs wild-type (wt) breast cancer which perfectly segregated ICR High versus Low samples in the BRCA TCGA cohort (MAPK-up genes) (Hendrickx et al., 2017), were also inversely correlated with ICR in a significant proportion of cancers (i.e, ACC, THYM, GBM, LGG and TGCT).

### Characterization of tumor mutational load and aneuploidy in relation to ICR immune phenotypes

Next, we aimed to identify genomic attributes related to the ICR immune phenotypes. As previously observed (Thorsson et al., 2018), mean neoantigen count of each cancer type strongly correlated with mean mutation rate (**Supplementary Figure 6A-B**). While mean non-silent mutation rate was significantly higher in ICR High tumors for some cancer types, no clear association was observed in most of them. Results for predicted neoantigen load were similar (**Figure 3A-B** and **Supplementary Figure 6C-D**). Overall, mean non-silent mutation rate and mean neoantigen load were higher in ICR-enabled cancers compared with ICR-disabled cancers. However, these differences cannot fully explain the divergent association of ICR with survival, as values for ICR-enabled cancers SARC and BRCA are in the same range as ICR-disabled cancers LGG, PAAD and KIRC.

**Figure 3:**
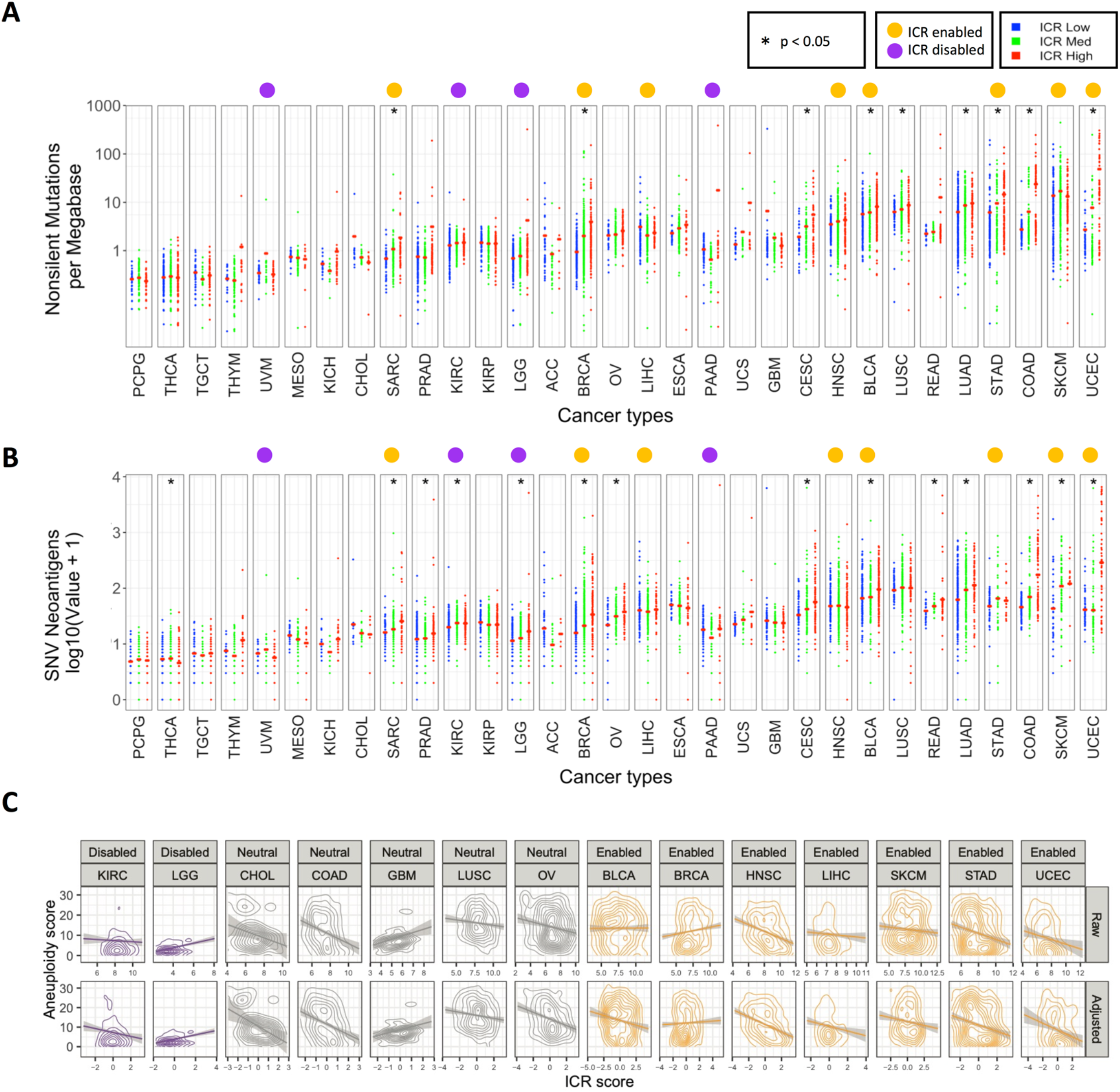
Association of ICR with nonsilent mutation rate, predicted neoantigen load, and tumor aneuploidy. **A.** Scatter plot of log transformed nonsilent mutation count per ICR cluster for each cancer type. **B.** Log transformed predicated neoantigen load per ICR cluster for each cancer type. **A.B.** Red crossbar represents the mean value per ICR cluster. Cancer types are ordered by mean nonsilent mutation count per cancer. Nonsilent mutation rate and predicted neoantigen load were obtained from Thorsson et al (Thorsson et al., 2018). **C.** Correlation between aneuploidy score and raw/purity adjusted ICR score for all cohorts with significant relationships between ICR and aneuploidy.

Similarly, we studied the association between genomic instabilities, or aneuploidy, and ICR. Specifically, we compared the individual tumor aneuploidy scores and the ICR score across cohorts. Aneuploidy score was calculated as in Taylor *et al* (Taylor et al., 2018). As has been reported previously, we found a broad negative association between aneuploidy and raw or tumor purity adjusted ICR score (Davoli et al., 2017) (**Figure 3C**). Interestingly, this negative association was most strongly supported in ICR-enabled cancers, with 6 cancers out of 8 showing a significant negative association between aneuploidy score and purity adjusted ICR (P < 0.01). In ICR-neutral cancers, a small fraction of cancer types showed a negative association (4 of 18, with an additional 4 showing a non-significant but suggestive negative association). Three cohorts (GBM, KICH and PRAD) showed a suggestive positive association. Similarly, in the ICR-disabled cohorts only KIRC showed a significant negative association, while LGG showed a strongly significant positive association (p-value < 10^-8^).

### Specific mutations associate with ICR immune phenotypes

To define the association of specific oncogenic mutations with ICR immune phenotypes, we first selected a set of 470 frequently mutated genes in cancer (Iorio et al., 2016), then trained an elastic net (Zou and Hastie, 2005) model to predict the ICR score as function of mutations in each sample and using the tumor-type as covariate. The positive non-zero coefficients of the trained model were used to identify genes whose mutation are associated with an increase of the ICR and negative non-zero coefficients identify the genes whose mutations are associated to a decrease of the ICR score (**Figure 4A**). The use of tumor-type as covariate tends to limit the effect of the enrichment of mutations in specific cancer-types and their correlation with ICR score. The coefficients of the tumor-type were all different from zero, with the exception of BLCA, BRCA, CHOL, COAD, READ and SARC and retained in the final model. We evaluated the accuracy of the model in a ten-fold cross-validation computing the correlation between the model prediction and the true ICR scores and obtaining a Spearman correlation of 0.669 ± *0.012* (p-value < 10^-400^). Genes associated with a decrease of ICR score include: *FOXA2, NSD1, PSIP1, HDAC3, ZNF814, FRG1, SOX17, CARM1, GATA3, FKBP5, FGFR3, MAT2A, PPP2R5A, MECOM, SMAD2, MED17, WNT5A, KRAS, ADAM10, PRKAR1A, DIS3, PRRX1*, *MFNG, TNPO1, SPOP, KDM6A, EGFR, IRF7, NRAS, SUZ12, RPSAP58*, and *SF3B1*.

**Figure 4:**
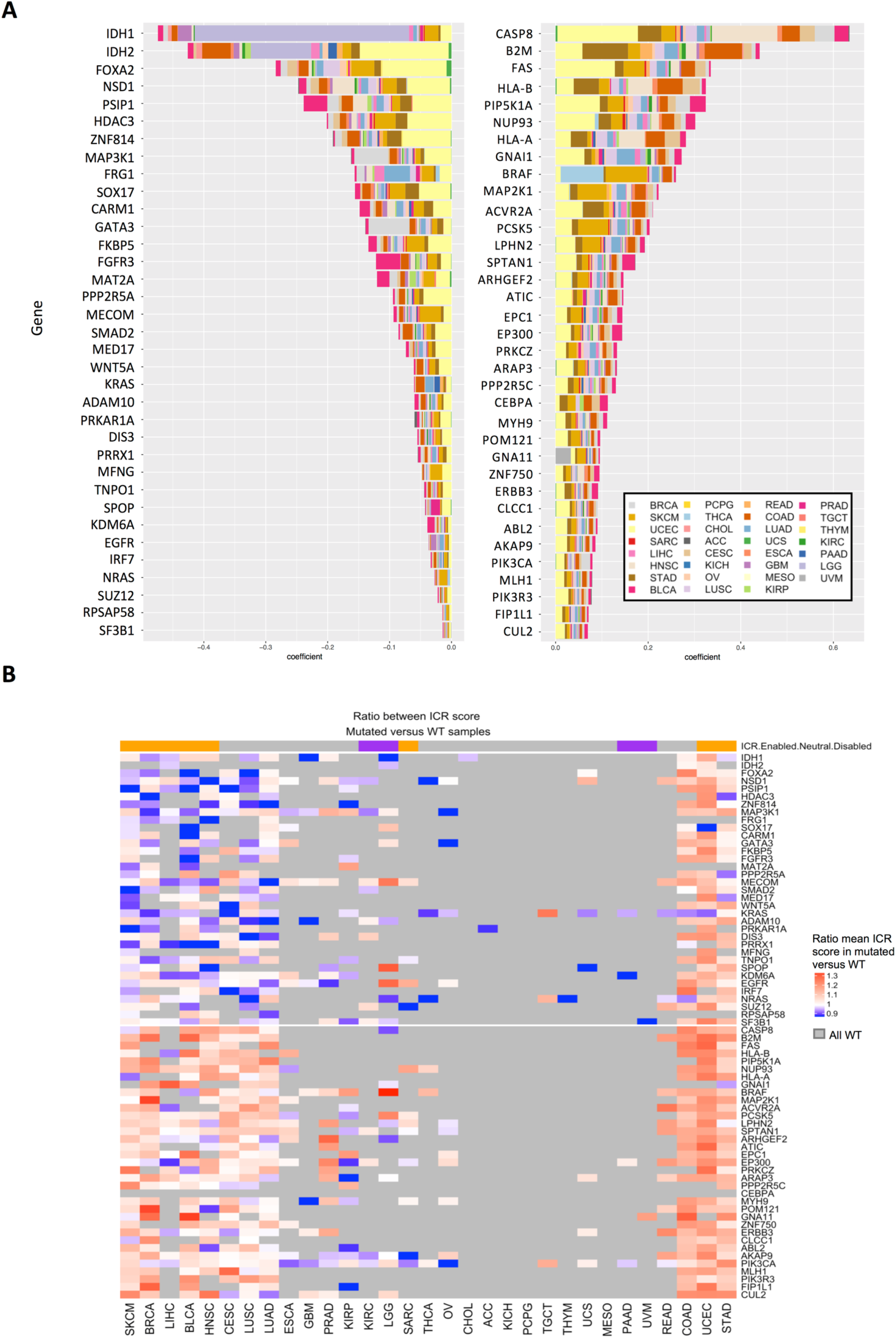
Relationship between ICR score and mutations in individual genes. **A.** Top 35 of mutated genes with negative non-zero coefficients of a trained elastic net model identified genes whose mutation is associated with a decrease of the ICR (left panel). Top 35 mutated genes with a positive association with ICR score in pan-cancer trained model (right panel). Contributions of each individual cancer type to the coefficient in trained elastic net model are proportionally indicated by size of the bars. **B.** Ratio of mean ICR score in mutated samples and ICR score in WT samples. Cancer types are ordered manually based on patterns of calculated ratios.

Interestingly MAP3K1 mutations, whose effect on ICR Low has been described in breast cancer (Hendrickx et al., 2017), were also associated to ICR Low tumors pan-cancer. The top genes of which mutations positively correlate with ICR reflect immune-evasion mechanisms that follow immunologic pressure such as mutations of antigen-presenting machinery transcripts previously described (i.e., *B2M, HLA-A, HLA-B*, and CASP8)(Rooney et al., 2015).

To better compare the association between specific mutations and ICR groups within individual cancer types, we calculated, for each of the identified genes, the mean ICR score in the mutated group divided by the mean ICR score in the wild type (WT) within each individual cancer type. For most cancer types, the genes with a positive coefficient consistently showed a higher ICR score in mutated samples, supporting their association with an ICR High phenotype (**Figure 4B**). On the other hand, genes with a negative coefficient (genes associated with an ICR Low phenotype) as identified at the pan-cancer level, do show some clear deviations between cancer types. While for most cancer types, ICR score is indeed lower in the mutated group, results for cancer types COAD, UCEC and STAD show the reverse (**Figure 4B**). Interestingly, a common characteristic of these three cancer types is frequent hypermutation as a consequence of microsatellite instability (MSI) (Cortes-Ciriano et al., 2017). This hypermutator phenotype could be responsible for the observed increased ICR score in the mutated group, as the genes with negative coefficient could be mutated in the context of hypermutation. We indeed observed an increased ICR score in the MSI-high group compared to MSI-low and microsatellite stable (MSS) groups in COAD and STAD datasets for which sufficient data on MSI status were available (Cortes-Ciriano et al., 2017) (**Supplementary Figure 7A-B**).

Mutated genes were frequently part of multiple pathways, suggesting impact on various tumor biological systems (**Supplementary Figure 8**).

### Prognostic impact of ICR classification is dependent on the expression of cancer-related pathways

Although we observed interesting differences between ICR High and ICR Low immune phenotypes across different cancer types, these do not explain the divergent association between immune phenotype and survival as we observed in ICR-enabled versus ICR-disabled cancer types (**Figure 1C-D**). As previously stated, an active immune phenotype has different impacts on survival depending on molecular subtype (for e.g. breast cancer (Miller et al., 2016)). To examine tumor intrinsic differences between ICR-enabled and ICR-disabled cancer types, we compared the enrichment of tumor intrinsic pathways between these two groups. Differentially enriched pathways (t-test; FDR <0.05; **Supplementary Table 2**) between ICR-enabled and disabled cancer types were selected and used for pan-cancer hierarchical clustering. Interestingly, a wide variety of pathways were differentially enriched between both groups. Whereas enrichment for pathways involved in proliferation were mostly upregulated in ICR-enabled cancer types (proliferation metagene (Miller et al., 2016), E2F targets, G2M checkpoints and mismatch repair), a large number of tumor intrinsic pathways (n=43) were enriched in ICR-disabled cancer types. Visualization of ES for these pathways across different cancer types in a heatmap confirms these findings. Hierarchical clustering based on ES of tumor intrinsic pathways differentially dysregulated by ICR-enabled and ICR-disabled cancer types segregates specimens into two main clusters (**Figure 5A**). As anticipated, pan-cancer survival analysis of all samples that formed a cluster along with samples of the ICR-disabled cancer types, named the *ICR non-beneficial cluster*, revealed no survival benefit of a high ICR expression. On the other hand, survival analysis of all samples in the other cluster, named the *ICR beneficial cluster*, showed a clear survival benefit for ICR High samples (**Figure 5B**). Of note, the prognostic significance of ICR was higher in this ICR beneficial cluster (HR = 1.82; p-value = 4.13^-9^; 95% CI = 1.49-2.23) compared to the prognostic significance of all samples of ICR-enabled cancer types combined (HR = 1.63, p = 2.26^-8^; 95% CI = 0.88-1.14), suggesting that tumor intrinsic attributes beyond the tumor site of origin are important to determine the relevance of cancer immune phenotypes. Interestingly, samples from ICR-neutral cancers, in which no clear trend was observed between ICR and survival (**Figure 1C**), and which were not used in calculation of differentially enriched pathways, were divided across the *ICR beneficial* and *ICR non-beneficial clusters*. To evaluate whether the prognostic impact of the ICR was relevant to a subset of samples from ICR-neutral cancer types, subgroup analysis was performed for samples of ICR-neutral cancer types. Indeed, for all samples from ICR-neutral cancer types that clustered to the *ICR non-beneficial cluster*, ICR was not associated with survival. On the other hand, for samples of ICR-neutral cancer types which clustered to the *ICR beneficial cluster*, ICR showed a significant positive association with survival (**Figure 5C**), indicating that the ICR has prognostic relevance in this subgroup of cancer patients as well.

**Figure 5:**
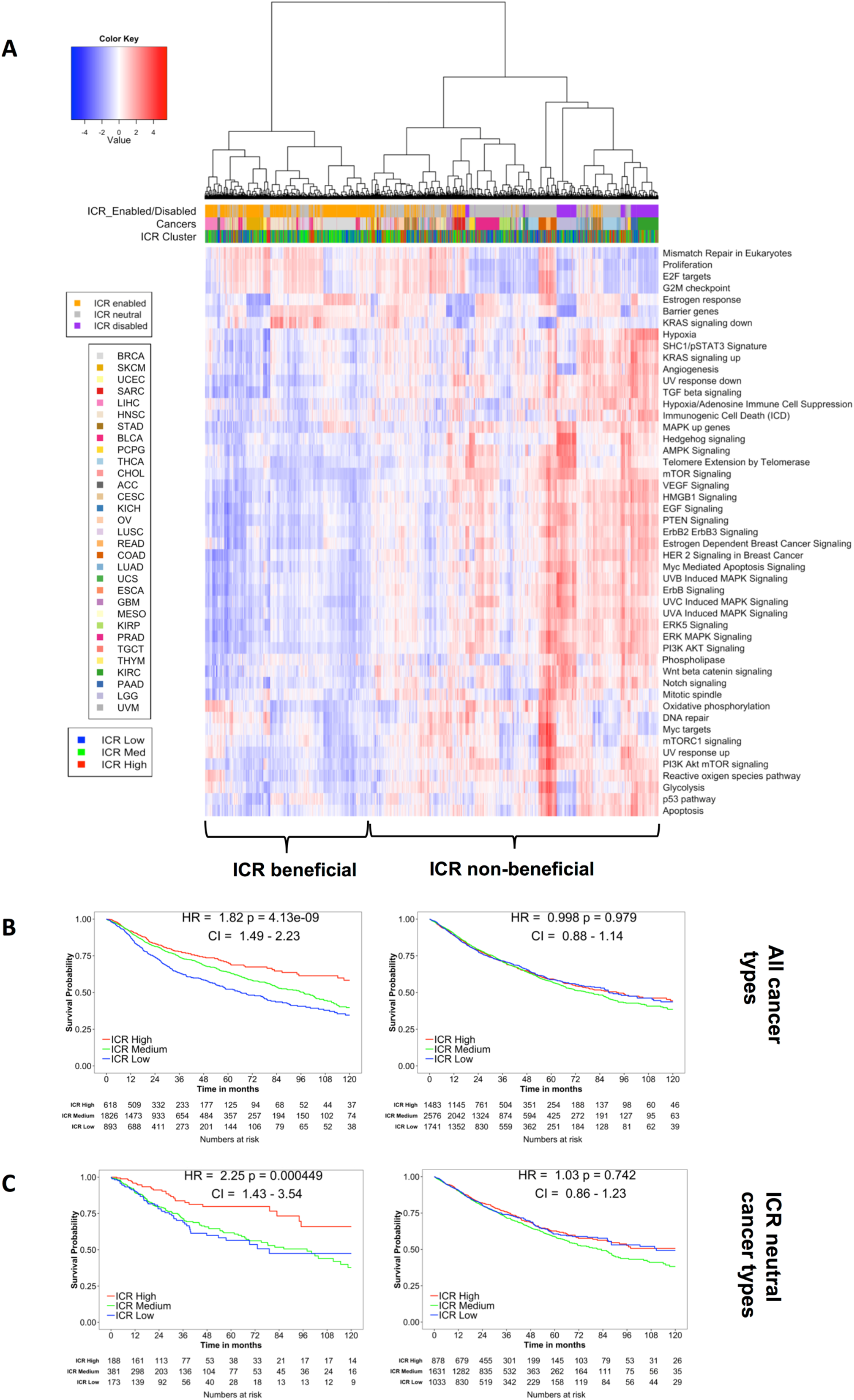
Pan-cancer clustering based on oncogenic pathway enrichment segregates ICR-enabled and ICR-disabled cancer types. **A.** Heatmap of enrichment scores of selected oncogenic pathways, samples are hierarchically clustered in two main clusters: one cluster consists mostly of ICR-enabled cancer types (ICR beneficial cluster), while the second cluster contains all samples from ICR-disabled cancer types (ICR non-beneficial cluster). **B.** Kaplan-Meier OS curves for ICR High, Medium, and Low clusters for samples in the ICR beneficial and ICR non-beneficial cluster separately. **C.** Subgroup survival analysis of all samples of ICR-neutral cancer types clustered in the ICR beneficial cluster and ICR non-beneficial cluster.

To better clarify this concept, we selected two of the differentially expressed pathways that were of special interest. Firstly, the “Proliferation” signature was used to classify all samples independent of tumor origin in “Proliferation High” and “Proliferation Low” categories, defined as an ES value >median or <median of all samples, respectively. This 52-gene cluster described by Nagalla *et al* (Nagalla et al., 2013) has previously been associated with the prognostic value of immune gene signatures in breast cancer (Miller et al., 2016). As represented by a histogram, the proportion of samples with high proliferation signature enrichment was larger in ICR-enabled cancer types compared with ICR-disabled cancers (**Figure 6A**). This very basic binary classification was already capable of segregating samples in a group in which ICR has a positive prognostic value from a group in which ICR is not associated with survival (**Figure 6B**). As a second illustration, “TGF-ß signaling” was used to classify samples based on this pathway using the same approach. For this oncogenic pathway, ICR-enabled cancer types typically had a lower enrichment of this pathway compared to ICR-disabled cancer types (**Figure 6C**). This classification could also divide samples in a group in which ICR has a positive association with survival and a group in which this association is absent (**Figure 6D**).

**Figure 6:**
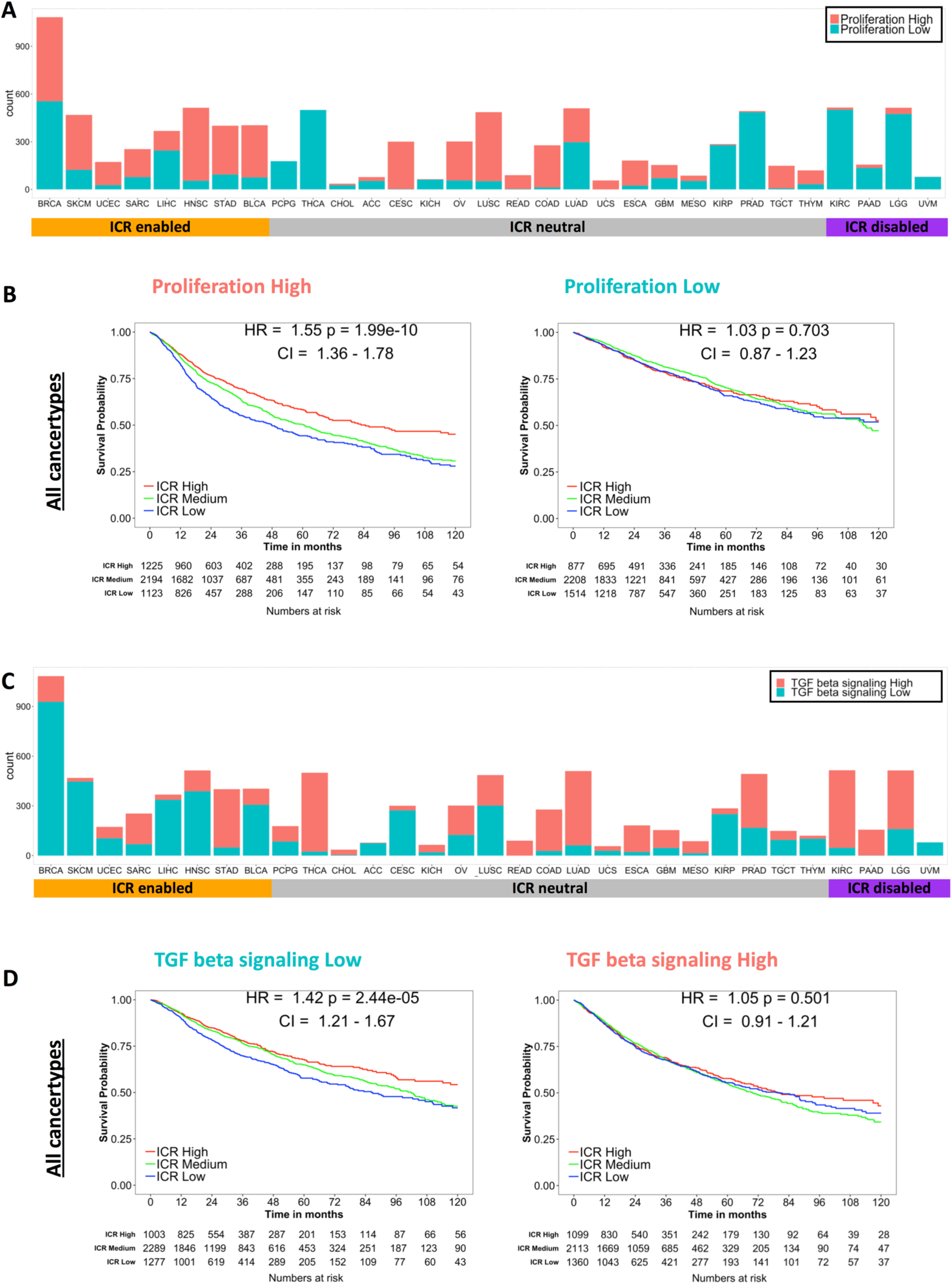
Examples of pan-cancer binary classifications based on enrichment of individual tumor intrinsic gene signatures and corresponding stratified pan-cancer survival analysis. **A.** Histogram showing pan-cancer classification based on median pan-cancer enrichment value of the proliferation signature as described by Miller *et* al(Miller et al., 2016) (Proliferation Low: ES is lower than median ES observed pan-cancer; Proliferation High: ES is higher or equal to median ES observed pan-cancer). **B.** Pan-cancer Kaplan Meier curves of ICR groups stratified by Proliferation High (left panel) and Proliferation Low (right panel) groups corresponding to classification as shown in panel A. **C.** Histogram showing pan-cancer classification based on pan-cancer enrichment values of the Hallmark pathway TGF-ß signaling. **D.** Pan-cancer Kaplan Meier curves stratified by TGF-ß signaling Low (left panel) and TGF-ß signaling High (right panel) groups corresponding to classification as shown in panel C.

As proliferation positively correlates with tumor mutational load (Pearson’s correlation coefficient = 0.49) (**Supplementary Figure 9**), we investigated whether tumor proliferation independently contributes to the prognostic value of ICR. Therefore, we segregated pan-cancer samples in four categories based on both mutation rate and proliferation (**Supplementary Figure 10**). Interestingly, in the proliferation high group, ICR High was associated with significantly improved survival independent of mutation rate. A similar observation is made for the mutation rate high group, ICR High is associated with better survival independent of proliferation. These finding suggest that mutation rate and enrichment of proliferation-related transcripts provide additive information to define the prognostic value of ICR. Furthermore, in a multivariate Cox proportional hazards model including ICR classification, proliferation enrichment, TGF-ß signaling enrichment, and tumor mutation rate, all parameters remain significant (**Supplementary Figure 11**). This implies that ICR, proliferation rate, TGF-ß signaling and tumor mutation rate all have independent prognostic value.

We then continued by verifying whether these tumor intrinsic attributes that interact with the prognostic impact of ICR when evaluated pan-cancer, could also translate to individual cancer types. For each individual cancer type, samples were divided by median ES for each of the selected pathways. ICR HRs (ICR Low vs. ICR High) were compared between each pathway-High and pathway-Low group for each cancer type (**Supplementary Figure S12A-B**). Overall, we indeed observed an increased HR for samples with a high enrichment of ICR enabling pathways for most cancer types. For samples with a high enrichment of ICR disabling pathways, the HR was indeed lower (**Supplementary Figure S12C**).

These data confirm an association between the prognostic impact of ICR classification and enrichment of oncogenic pathways in individual cancer types as well as pan-cancer. Of note, these interactions between the prognostic significance of ICR and tumor intrinsic pathways were mostly present in enabled and neutral cancer types. Within disabled cancer types, with the exception of KIRC, similar associations were not found.

### Predictive value of ICR score in immune checkpoint therapy is dependent on proliferation and TGF-ß signaling

To define the clinical relevance of classification of ICR immune phenotypes, in the setting of immune checkpoint treatment, we first evaluated the predictive value of ICR score across multiple public datasets of anti-CTLA4 and anti-PD1 treatment. A significantly increased expression of ICR in responders compared to non-responders was observed across most of the datasets (**Figure 7A**) (Chen et al., 2016; Hugo et al., 2016; Prat et al., 2017; Riaz et al., 2017; Van Allen et al., 2015). The conditional activation of the prognostic impact of the ICR was tested in the Van Allen dataset, which was the only one for which survival information was available. Strikingly, in the proliferation high subgroup, ICR score was significantly higher in pre-treatment samples of patients with long-survival or response (p=0.021), whereas this difference was not significant in proliferation low samples (**Figure 7B**). Cohort dichotomization based on TGF-ß signaling, again demonstrated the reverse trend: a significant difference in ICR score was only observed in the TGF-ß signaling low group (p=0.0044), not in the TGF-ß high group. Stratified survival analysis in these categories confirmed that the prognostic impact of ICR depends on proliferation and TGF-ß signaling (**Figure 7C**). These findings confirm a conditional prognostic and predictive impact of ICR based immune infiltration estimates in the setting of immune checkpoint treatment and demonstrate that these findings might have important clinical implications.

**Figure 7:**
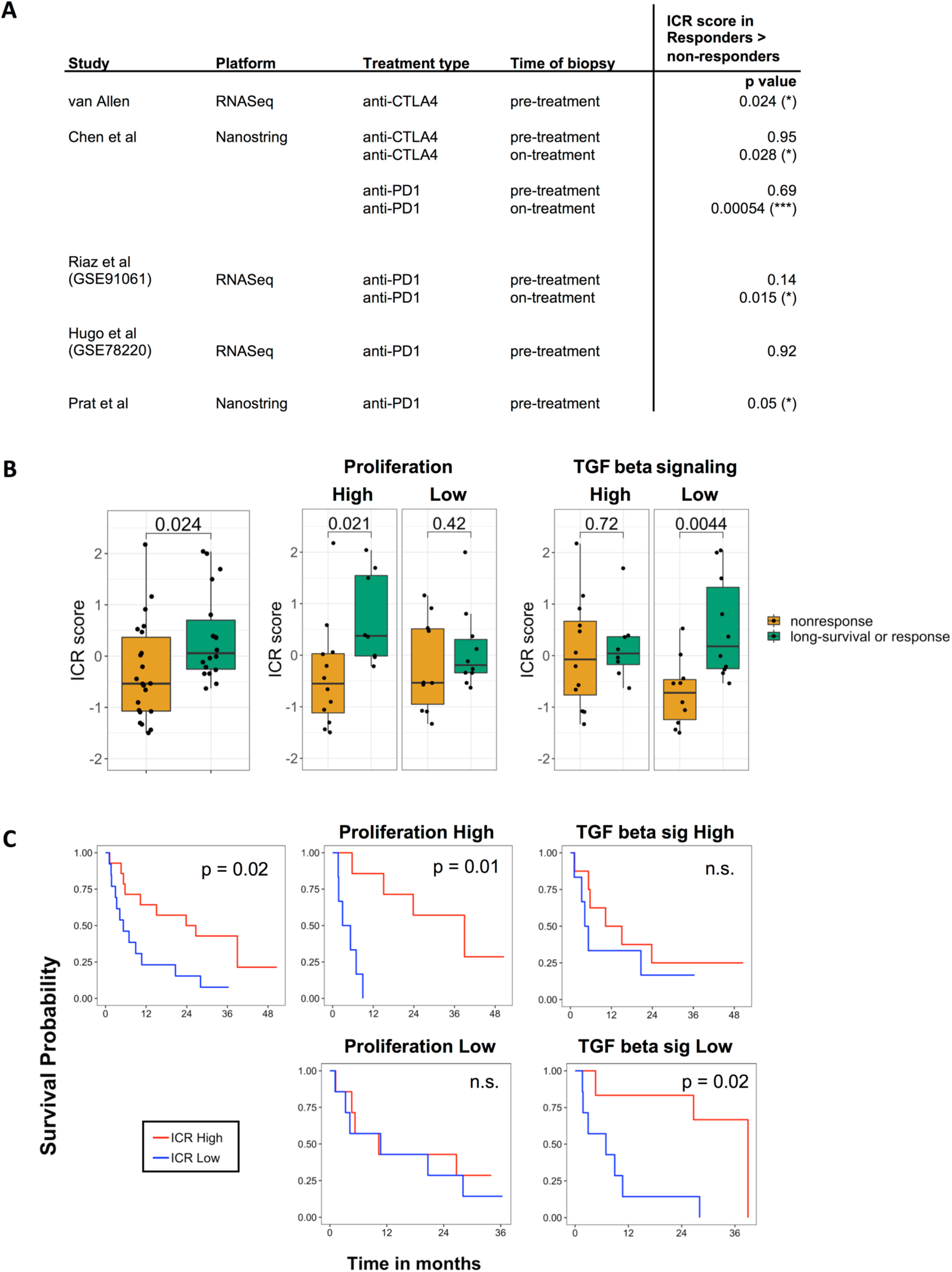
Conditional predictive value of ICR for response to immune checkpoint treatment. **A.** Predictive value of ICR across public datasets with response to immune checkpoint treatment indicated by p-value of two-sided t-test comparing ICR score in samples of responding versus non-responding patients. ICR score was highest in response group for all significant comparisons. Response was defined as long-survival or response in the van Allen dataset, stable disease, partial response (PR) and complete response (CR) in the Chen dataset, and as PRCR in Riaz, Hugo and Prat datasets. **B.** Boxplot of ICR score in “nonresponse” compared with “long-survival or response” to anti-CTLA4 treatment in van Allen dataset (left). Boxplots of subgroup analysis of proliferation groups (middle) and TGF-ß signaling groups (right). P-value of t-test comparing means are indicated in the plot. **C.** Kaplan Meier curves showing OS across ICR tertiles in all samples (left), across proliferation (middle), and TGF-ß signaling subgroups (left).

## Materials and Methods

### Data acquisition and normalization

RNA-seq data from The Cancer Genome Atlas (TCGA) were downloaded and processed using TCGA Assembler (v2.0.3). Gene symbols were converted to official HGNC gene symbols and genes without symbol or gene information were excluded. RNA-seq data from as wide as possible sample set of the total of 33 available cancer types of tissue types Primary Solid Tumor (TP), Recurrent Solid Tumor (TR), Additional-New Primary (TAP), Metastatic (TM), Additional Metastatic (TAM) and Solid Tissue Normal (NT) were used to generate a *pan-cancer* normalized dataset. Normalization was performed within lanes, to correct for gene-specific effects (including GC-content and gene length) and between lanes, to correct for sample-related differences (including sequencing depth) using R package EDASeq (v2.12.0) and quantile normalized using preprocessCore (v1.36.0). After normalization, samples were extracted to obtain a single primary tumor tissue (TP) sample per patient. For SKCM patients without available TP sample, a metastatic sample (TM) was included. Finally, the pan-cancer normalized dataset was filtered to remove duplicate patients and samples that did not pass assay-specific QCs (Thorsson et al., 2018) data was log2 transformed. Clinical data were sourced from the TCGA Pan-Cancer Clinical Data Resource (Liu et al., 2018). Mutation rate and predicted neoantigen load were obtained from the recent immunogenomic analysis by Thorsson *et al* (Thorsson et al., 2018). The dataset published by Ellrott *et al* was used for mutation data analysis(Ellrott et al., 2018). Hematological cancer types LAML and DLBC were excluded from analysis.

Raw fastq files of datasets GSE78220 (Hugo et al., 2016) and GSE78220 (Riaz et al., 2017) were downloaded from NCBI SRA servers, quality control and adapter trimming was performed using Trim_Galore (https://github.com/FelixKrueger/TrimGalore). Reads were aligned to hg19 using STAR (Dobin et al., 2013). GenomicFeatures and GenomicAlignments Bioconductor packages were used to generate row counts. The raw counts were normalized with EDASeq (Risso et al., 2011) and log2 transformed. The dataset phs000452.v2.p1 (Van Allen et al., 2015) was downloaded, already normalized, from http://tide.dfci.harvard.edu/.

### ICR classification

Consensus clustering based on the 20 ICR genes (**Figure 1A**) was performed for each cancer type separately using the ConsensusClusterPlus (v1.42.0) R package with the following parameters: 5,000 repeats, a maximum of six clusters, and agglomerative hierarchical clustering with ward criterion (Ward.D2) inner and complete outer linkage. The optimal number of clusters (≥ 3) for best segregation of samples based on the ICR signature was determined heuristically using the Calinski-Harabasz criterion(Caliński and Harabasz, 1974) (<u>source function</u> available on GitHub repository, see cancer datasheets for plots with local maximum). As we were interested to compare cancer samples with a highly active immune phenotype with those that have not, the cluster with the highest expression of ICR genes was designated as “ICR High”, while the cluster with the lowest ICR gene expression was designated “ICR Low”. All samples in intermediate cluster(s) were defined as “ICR Medium”. Samples were annotated with an ICR score, defined as the mean of the normalized, log2 transformed expression values of the ICR genes. For generation of the ICR Heatmaps (**Figure 1C** and the cancer datasheets), a modified version of heatmap.3 function was used (<u>source function</u>).

### Survival analysis

Overall survival (OS) from the TCGA Pan-Cancer Clinical Data Resource (Liu et al., 2018) was used to generate Kaplan-Meier curves using a modified version of the ggkm function (Abhijit, n.d.). Patients with less than one day of follow-up were excluded and survival data were censored after a follow-up duration of 10 years. Hazard ratios (HR) between ICR Low and ICR High groups, including corresponding p-values based on chi-squared test, and confidence interval were calculated using the R package survival (v2.41-3). The forest plot (**Figure 1C**) was generated using the R package forestplot (v1.7.2). Cancer types PCPG, THYM, TGCT, and KICH were excluded before generation of the plot, as the number of deaths in the comparison groups was too small for calculations. Cancer types with a HR > 1 with a p-value < 0.1 were termed ICR-enabled and cancer types with a HR < 1 with a p-value < 0.1 were termed ICR-disabled. The forest plot was annotated manually with indicators for ICR-enabled and ICR-disabled cancer types. Cox proportional hazards regression analysis was performed with the R package survival with the AJCC pathologic tumor stage as described in the TCGA Pan-Cancer Clinical Data Resource (Liu et al., 2018). For simplification, stage categories were reduced to “Stage I”, “Stage II”, “Stage III” and “Stage IV” for subcategories (e.g. Stage IIA, Stage IIB, Stage IIC etc). In multivariate analysis, pathologic stage was entered as a semi-continuous (ordinal) variable. Cancer types LGG and GBM were not included in the multivariate analysis as tumor stage is unavailable (not applicable) for these histologies.

### Gene Set Enrichment Analysis

To define the enrichment of specific gene sets, either reflecting immune cell types (**Figure 2A**) or specific oncogenic pathways (**Figure 2B**), single sample GSEA (Barbie et al., 2009) was performed on the log2 transformed, normalized expression data. Immune cell-specific signatures as described in Bindea *et al* (Bindea et al., 2013) were used as gene sets using this method to deconvolute immune cell abundance. Gene sets to define enrichment of specific tumor-related pathways were obtained from the multiple sources. We started with a selection of 24 Hallmark pathways (Liberzon et al., 2015) which are regularly expressed in cancer. Subsequently, we added 21 non-redundant Ingenuity Pathway Analysis (IPA) pathways (http://www.ingenuity.com, Ingenuity System Inc., Redwood City, CA, USA). Finally, several pathways were added that have previously been hypothesized to associate with cancer immune phenotypes, including Hypoxia/Adenosine Immune Cell Suppression, Immunogenic Cell Death (ICD), NOS1 Signature, PI3Kgamma signature, and SHC1/pSTAT3 signatures as described by Lu *et al* (Lu et al., 2017), barrier genes as described by Salerno *et al* (Salerno et al., 2016), the proliferation metagene as described by Miller *et al* (Miller et al., 2016) and genes upregulated in MAPK mutated breast cancer (Bedognetti et al., 2017).

### Correlation matrix

The correlation matrices of ICR genes (**Supplementary Figure 1**) and correlation between ICR score and ES of selected pathways (**Figure 2B**) were calculated using Pearson test and plotted using “corrplot” version 0.84.

### Mutational Analysis

Mutation rate and predicted neoantigen count data (Thorsson et al., 2018) were log10-transformed and distribution across ICR clusters was plotted using R package “ggplot2”. Differences between ICR High, Medium and Low clusters were calculated through t-test, using a cut-off p-value of < 0.05. For specific mutation analysis, a set of 470 frequently mutated genes in cancer (Iorio et al., 2016), was selected. An elastic net regularized (Zou and Hastie, 2005) model was built to predict the ICR score as function of mutations in each sample and using the tumor-type as a covariate. The accuracy of the model was evaluated in a ten-fold cross-validation setting computing the correlation between the model prediction and the true ICR scores, finally obtaining a Spearman correlation of 0.669 ± 0.012 (p-value < 10^-400^).

The R package “ComplexHeatmap” was used to plot ICR score ratios between mutated versus wild-type groups. For cancer type/ gene combinations with a number of samples of <3 in the mutated group, ratios were not calculated (NA; grey color in plot). A ratio >1 implies that the ICR score is higher in the mutated group compared with WT, while a ratio <1 implies that the ICR score is higher in subset of tumors without mutation.

### Aneuploidy

Aneuploidy scores for each individual cancer were taken from Taylor *et al* (Taylor et al., 2018). Briefly, each tumor was scored for the presence of aneuploid chromosome arms after accounting for tumor ploidy. Tumor aneuploidy scores for each cohort were then compared to ICR scores via linear model with and without purity adjustment. Purity adjustment entailed correlating ICR score and tumor purity (as estimated via ABSOLUTE) and using the residuals to evaluate the post-adjustment relationship between ICR score and tumor aneuploidy. In particular we made use of the precomputed aneuploidy scores and ABSOLUTE tumor purity values. Raw ICR and aneuploidy score associations were evaluated by linear model in R via the *lm()* function for each cohort independently. Adjusted ICR and aneuploidy score associations were evaluated by first modeling ICR score by tumor purity, then taking the ICR score residuals and assessing the association with aneuploidy score via linear model. Cohorts with model p-values below 0.01 for adjusted or unadjusted ICR score and aneuploidy, regardless of the directionality of the association, were included in **Figure 3C**.

### Differential GSEA and stratified survival analysis

Differential ES analysis between samples of ICR-enabled and those of ICR-disabled cancer types was performed using t-tests, with a cut-off of FDR-adjusted p-value (i.e., q-value) < 0.05 (**Supplementary Table 2**). Tumor intrinsic pathways that were differentially enriched between ICR-enabled and disabled cancer types were selected. The heatmap used for visualization of these differences was generated using the adapted heatmap.3 function (<u>source function</u>). For each of these selected pathways, samples were categorized pan-cancer as pathway-High (ES > median ES) or pathway-Low (ES < median ES). Associations between ICR and survival were defined for each pathway “High” and pathway “Low” group separately using the survival analysis methodology as described above. Pathways for which a significant association between ICR and survival was present in one group, but not in the other one, were selected (**Supplementary Table 3**). Similarly, these pathways were used to categorize samples per individual cancer type in pathway-High (ES > cancer specific median ES) and pathway-Low (ES < cancer specific median ES). Differences between HRs of groups in individual cancer types were calculated and plotted using “ComplexHeatmap” (v1.17.1).

### Predictive value ICR score in immune checkpoint datasets

ICR scores, or the mean expression of ICR genes, were compared between responders and non-responders to immune checkpoint therapy. For the Chen et al dataset, performed on Nanostring platform, scores were calculated using the 17 ICR genes available in the nanostring panel. Difference in mean ICR score between groups was tested using two-side t-test (cutoff <0.95) (**Fig 7A**). For datasets, GSE78220 (Riaz et al., 2017), GSE78220 (Hugo et al., 2016) and Prat *et al* (Prat et al., 2017), the response category includes both partial and complete clinical responders according to respective publications. For Chen et al, clinical responders also included stable disease, as described by Chen *et al* (Chen et al., 2016). Dataset van Allen *et al*, response was defined as patients with clinical response or long-term survival after treatment (Van Allen et al., 2015). Samples of van Allen dataset were dichotomized based on median ssGSEA of 1) genes of the proliferation metagene and 2) TGF-ß signaling signature. Stratified analysis was performed in each of the categories. ICR High, Medium and Low groups were defined according to ICR score tertiles, to obtain groups of sufficient size. Stratified survival analysis was performed using the same approach as applied to the TCGA data.

## Discussion

Transcriptional signatures used to define the continuum of cancer immune surveillance and the functional orientation of a protective anti-tumor immunity typically reflect common immune processes and include largely overlapping genes (Ayers et al., 2017; Hendrickx et al., 2017; Wang et al., 2008). We termed this signature as the ICR (Galon et al., 2013; Wang et al., 2008).

In our systematic analysis we showed that, across and within different tumors, the coordinated overexpression of ICR identifies a microenvironment polarized toward a Th-1/cytotoxic response, which was then used to define the hot/immune active tumors.

In tumor types with medium/high mutational burden, the mutational or neoantigenic load tended to be higher in hot (ICR high) vs cold (ICR low) tumors while this association was not observed within cancer types with overall low mutational burden. By adding granularity to previous observations that described an overall weak correlation between immunologic correlates of anti-tumor immune response and mutational load (Danaher et al., 2018; Ock et al., 2017; Rooney et al., 2015; Spranger et al., 2016; Thorsson et al., 2018), we demonstrated here that the differences in term of mutational load was especially evident in tumors types known to be constituted by a significant proportion of microsatellite instable cases, such as COAD, STAD and UCEC. It is likely that, in hypermutated tumors, the excess of neoantigens plays a major role in the immune recognition, while, in the other cases, additional mechanisms, such as cell-intrinsic features, play a major role in shaping the anti-tumor immune response (Hendrickx et al., 2017). Overall, a high mutational/neoantigen load was neither sufficient nor necessary for the displaying of an active immune microenvironment.

When the ICR score was intersected with the enrichment of oncogenic signals as predicted by the transcriptional data, interesting associations emerged. Although some differences in terms of the degree of the correlation were observed across cancers, few tumor-cell intrinsic pathways displayed a coherent progressive enrichment in the immune-silent tumors. The top pathways associated with the absence of the Th1/hot immune phenotype included, barriers genes, WNT-ß catenin, mismatch repair, telomerase extension by telomerase, Notch, Hedgehog, and AMPK signaling pathways. Barrier genes encode for molecules with mechanical barrier function in the skin and other tissues and include filaggrin (FLG), tumor-associated calcium signal transducer 2 (TACSTD2), desmosomal proteins (DST, DSC3, DSP, PPL, PKP3, and JUP) (Salerno et al., 2016). Their expression was associated with a T-cell excluded phenotype in melanoma and ovarian cancer, and here we extended our previous observation across multiple tumors (Salerno et al., 2016). The cell-intrinsic WNT-ß catenin activation impairs CCL4-mediated recruitment of Batf3 dendritic cells, followed by absence of CXCL10 mediated T-cell recruitment, and was described initially associated with T-cell exclusion in melanoma, and recently, in other tumor types (Luke et al., 2019; Spranger et al., 2015). The efficiency of our approach in capturing previously described oncogenic pathways indicates the robustness of the analysis. At the same time, our integrative pipeline unveiled additional relevant pathways: telomere extension by telomerase and mismatch repair, Notch, Hedgehog and AMPK signaling. Our findings suggest that the lack of expression of transcripts involved with mismatch repair (in addition to their genetic integrity (Barnetson et al., 2006)) might influence immunogenicity. Telomere dysfunctions result in various disease, including cancer and inflammatory disease (Calado and Young, 2012). To our knowledge, this is the first time that telomerase activity has been linked to differential intratumor immune response. The Notch pathway can regulate several target genes controlled by the NFκB, TGF-β, mTORC2, PI3K, and HIF1α pathways (Janghorban et al., 2018) and is involved in the induction of cancer stem cells, but has not been described to be associated with differential intratumoral immune response so far. As for the Hedgehog pathway, in breast cancer models, inhibition of this signaling induces a marked reduction in immune-suppressive innate and adaptive cells paralleled with an enrichment of cytotoxic immune cells (Hanna et al., 2019). Intriguingly, the AMPK pathway was the most coherently dysregulated pathway in relationship to the ICR score. In lung cancer mouse models, the deletion of LKB1 (an upstream modulator of AMPK pathway) was associated with decrease T cell tumor infiltration, and impaired production of pro-inflammatory cytokines, which was mediated by induction of STAT3 and IL-6 secretion (Koyama et al., 2016; Spranger and Gajewski, 2018). The strength of the inverse association between the AMPK pathway and ICR score strongly calls for in-depth investigation of the immune-modulatory role of this pathway. Overall, we identified novel putative hierarchically relevant cancer-cell intrinsic pathways associated with immune evasion mechanisms in humans that might warrant further mechanistic investigations and that might be explored as targets for reprogramming the tumor microenvironment. The biological relevance here is substantiated by the consistency of the associations across tumor types, in which each cohort can be seen as an independent validation. The coherence of the associations rules out the possibility of a spurious correlation.

As for somatic mutations, the top ten genes associated with the immune silent phenotype include *IDH1, IDH2, FOXA2, NSD1, PSIP1, HDAC3, ZNF814, MAP3K1, FRG1 and SOX17*. Findings of *IDH1* and *NSD1* are consistent with the report of Thorsson et al (Thorsson et al., 2018), in which these have been associated with decreased leukocyte infiltration, and are complemented here by additional identification of *IDH2*. Interestingly, MAP3K1 mutations were previously associated with low ICR in breast cancer in our previous work (Bedognetti et al., 2017; Hendrickx et al., 2017). Remarkably, mutations of other genes of the RAS/MAPK pathways such as *FGFR3* (previously associated with T-cell exclusion in bladder cancer (Sweis et al., 2016)), *EGFR, NRAS*, and *KRAS* were associated with a low ICR score, substantiating their potential role in mediating immune exclusion. FOXA2 is involved in both neoplastic transformation and epithelial-mesenchymal transition (Wang et al., 2018, p. 2) and T helper differentiation (Chen et al., 2010) but its role in modulating anti-tumor immune response is unknown. Similarly, no data exists on the effect of *HDAC3, PSIP1* and *ZNF814* on tumor immunogenicity. Considering the strength of the association, further investigations should mechanistically address the role of these signaling pathways in mediating immune evasion mechanisms. Other mutations associated with the immune silent phenotype include *WNT5A* (corroborating the immune-suppressive role of the WNT ß catenin pathway (Luke et al., 2019)), and *GATA3*, which was also previously associated with low leukocyte infiltration (Thorsson et al., 2018). Mutations of *FKBP5, MAT2A, PPP2R5A, MECOM, SMAD2, MED17, ADAM10, PRKAR1A, DIS3, PRRX1, MFNG, TNPO1, KDM6A, IRF7, SUZ12, RPSAP58*, and *SF3B1* represent additional novel findings. Similar to previous observations, we found *HLA-A, HLA-B, B2M, CASP8* and *FAS* to be associated with an ICR High immune phenotype (Ock et al., 2017; Rooney et al., 2015; Shukla et al., 2015; Siemers et al., 2017; Thorsson et al., 2018). These mutations are probably the result of immune escape mechanisms triggered by immunologic pressure.

As for genomic instability, tumors with high aneuploidy are associated with decreased ICR score in a major subset of cancer types (Davoli et al., 2017). This observation is also in agreement with negative association of a chromosome-instable type with an immune signature that predicts response to immunotherapy with MAGE-A3 antigen as well as response to anti-CTLA-4 treatment in melanoma (Ock et al., 2017). The only exceptions we found were brain tumors LGG and GBM in which a positive association between aneuploidy and ICR score was detected. In LGG tumors, however, ICR scores positively correlate with tumor grade (**Supplementary Figure 4**), and it is possible that the observed positive correlation between aneuploidy and ICR is actually driven by the higher genomic instability characterizing the more advanced tumors.

To compare cancer types based on the prognostic value of ICR, we categorized them into two groups: one for which ICR High was associated with increased OS and one for which ICR was associated with worse OS. For the first group, multivariate analysis confirmed a positive prognostic value of ICR independent of pathologic tumor stage. SKCM, BRCA, UCEC, LIHC, SARC, HNSC, STAD, and BLCA are consequently referred to as **ICR-enabled** cancer types. For the second group, including UVM, LGG, PAAD, and KIRC (**ICR-disabled** tumors), survival analysis showed a detrimental (univariate analysis) or neutral (multivariate analysis with stage) role of ICR.

These discrepancies in term of prognostic implication of intratumoral immune response have been observed in independent investigations based on transcriptomic analysis (Chifman et al., 2016; Thorsson et al., 2018) or immunohistochemistry (Fridman et al., 2012) but never explained.

The first notable difference we observed between ICR-enabled and -disabled cancer types was the overall lower ICR value in the disabled cancer cohorts. In particular for UVM and LGG, this low ICR could be a partial explanation for the lack of positive prognostic value of the ICR. On the other hand, mean ICR score of PAAD and KIRC was not different compared with the other cancer types. Therefore, other factors must have an effect on the prognostic value of the ICR. When we compared enrichment of tumor-cell intrinsic pathways in ICR-enabled and -disabled cancer types, as much as 43 of 54 analyzed pathways showed differential enrichment between the two groups. While ICR-enabled cancer types are typically more enriched in proliferation-related signatures, ICR-disabled cancer types have high enrichment of pathways generally attributed to tumor signaling including known pathways associated with immune suppression such as TGF-ß (Chakravarthy et al., 2018). In fact, when samples of the entire cohort were segregated according to representative enabling and disabling pathways (i.e., proliferation and TGF-ß signaling, respectively), the prognostic role of ICR was restricted to proliferation high/TGF-ß signaling low tumors (**Figure 6**). Hierarchical clustering based on the enrichment of transcripts of these differentially enriched pathways segregated most samples of ICR-enabled cancer types from samples of ICR-disabled cancer types. Interestingly, this clustering was even relevant to samples of ICR neutral cancer types. The pan-cancer survival analysis of samples of ICR neutral cancer types showed that for samples that co-clustered with samples of ICR-enabled cancer types (the ICR beneficial cluster), ICR High was associated with significant prolonged survival. Conversely, in samples of ICR neutral cancer types clustered to the ICR non-beneficial cluster, ICR lost its prognostic value. Adding the mutational load component further refined this stratification. In fact, the positive prognostic role of ICR was present also in a subset of samples with low proliferation and high mutational load but absent only in tumors with both low proliferation and low mutational load. We hypothesize that, in tumor with high mutational burden and/or high proliferative capacity, the high level of ICR captures a true protective anti-tumoral immune response, while in the other cases, such as in tumors dominated by TGF-ß signaling and low proliferation, the high ICR captures a bystander, or heavily suppressed, lymphocyte infiltration with no protective effect. Therefore, it is possible to speculate that a proportion of phenotypically immune active tumors are functionally immune silent. Single cell RNA sequencing, T-cell receptor sequencing, and spatial transcriptional analysis might be employed to characterize with higher fidelity the true functional orientation of human tumors.

The clinical relevance of the observed conditional impact of ICR was confirmed in the setting of anti-CTLA4 treatment, in which the predictive value of ICR was demonstrated to be dependent on tumor intrinsic pathways, such as TGF-ß and proliferation. To the best of our knowledge, we are the first to report an interaction between tumor intrinsic pathways and the prognostic value of immune phenotypes in a pan-cancer analysis. An association between proliferation and the prognostic value of immune phenotypes has previously been identified in breast cancer (Miller et al., 2016). In non-small cell lung cancer, proliferation was shown to improve prediction of immune checkpoint inhibitors response in PD-L1 positive samples (data recently presented at SITC annual meeting 2018 (“SITC 2018 Annual Meeting Schedule,” n.d.)). Our study clearly demonstrates that such interactions between tumor intrinsic attributes and prognostic and potentially predictive value of immune phenotypes are also relevant in a pan-cancer context. Moreover, we defined additional tumor intrinsic attributes beyond tumor proliferation to correlate with the prognostic significance of immune signatures reflecting a Th1 immune response. Prognostication algorithms should be refined by inclusion of tumor intrinsic attributes in order to define the prognostic impact of the immune signatures.

In conclusion, we observed a clear relationship between enrichment of tumor intrinsic pathways and the prognostic and predictive significance of the immune signatures and identified novel cell-intrinsic features associated with immune exclusion. This information can be used to prioritize candidates for immunogenic conversion and to refine stratification algorithms.

## Author contributions

J.R. contributed to the conception and design of the work, data acquisition and data interpretation, performed data analysis, and drafted the manuscript;

W.H. contributed to the conception and design of the work, data acquisition and data interpretation, performed data analysis, and drafted the manuscript;

P.K. contributed to the design of the work, interpretation of data, and substantively revised the manuscript;

R.M. contributed to analysis and interpretation of data;

G.Z. contributed to the analysis and substantively revised the manuscript;

M.S. contributed to interpretation of data and revision of manuscript;

K.H. contributed to the analysis and revision of manuscript;

G.C. contributed to interpretation of data;

D.R. contributed to the acquisition and analysis, interpretation of data, and revision of manuscript;

J.D. contributed to data interpretation and writing of the manuscript;

L.D. contributed to data interpretation and writing of the manuscript;

T.T. contributed to the analysis and revision of manuscript;

J.S. contributed to the analysis and revision of manuscript;

L.C. contributed to data interpretation and writing of the manuscript;

E.W. contributed to data interpretation, and substantively revised the manuscript;

P.F. substantively contributed to the manuscript;

F.B. contributed to data interpretation and writing of the manuscript;

L.M. contributed to the conception and design of the work, interpretation of data, and substantively revised the manuscript;

J.G. contributed to data interpretation and revision of manuscript;

F.M. contributed to data interpretation, and substantively revised the manuscript;

M.C. contributed to the conception and design of the work, data acquisition, data analysis and data interpretation, and substantively revised the manuscript;

D.B. conceived and designed the study, contributed to the acquisition, data analysis and data interpretation, supervised the analysis and drafted the manuscript.

## AbbVie disclosures

Tolga Turan, Josue Samayoa, and Michele Ceccarelli are all employees of AbbVie.

## Materials and Correspondence

Michele Ceccerelli

Wouter Hendrickx

Davide Bedognetti

## Ethics approval and consent to participate

Not applicable.

## Availability of data and material

Scripts for analysis are shared on https://github.com/Sidra-TBI-FCO/ISPC.git. Data generated or analyzed during this study are included in this article. Data sheets for results of individual cancer types are available as cancer data sheets at figshare (https://figshare.com/articles/Cancer_Datasheets/7937246).

## Supplementary Figures

**Supplementary Figure 1:** Pearson correlation between RNA-seq expression values of ICR genes for each of the 31 cancer types.

**Supplementary Figure 2:** Scatterplot showing correlation between ICR score and TIS score(Danaher et al., 2018) (**A**), ICR score and leukocyte fraction (**B**), and ICR score and TIL percentage (**C**). Leukocyte fraction and TIL percentage values were obtained from Thorsson *et al* (Thorsson et al., 2018). Each dot represents a single sample.

**Supplementary Figure 3: A.** Boxplot showing mean ICR score for each cancer type per group of cancer types: ICR-enabled, ICR-neutral and ICR-disabled. A single dot represents a single cancer type. **B.** Boxplot showing delta between mean ICR score in ICR High cluster compared with mean ICR score in ICR Low cluster. A single dot represents a single cancer type.

**Supplementary Figure 4:** Pan-cancer Kaplan-Meier curves in ICR-disabled (top left panel) and ICR-enabled (top right panel) groups and stratified analysis by AJCC pathologic stage I & II (middle panels) and stage III & IV (bottom panels).

**Supplementary Figure 5:** Dotted heatmap showing mean ES for each immune cell population per cancer type, mean ES scores were z-scored per row.

**Supplementary Figure 6: A.** Scatterplot of mean mutation rate versus mean neoantigen load per cancer type. **B.** Ratio of nonsilent mutation rate between ICR High and ICR Low groups versus the ratio of predicted neoantigen load between in ICR High compared to ICR Low groups. **C.** Ratio of nonsilent mutation rate between ICR High and ICR Low groups versus mean nonsilent mutation rate. **D.** Ratio of predicted neoantigen load between ICR High compared to ICR Low groups versus mean predicated neoantigen load.

**Supplementary Figure 7: A.** Boxplot of ICR score by MSI status in COAD (left panel) and STAD (right panel). P-values of t-test to compare mean ICR score per MSI group are indicated in the plot. **B.** Boxplot of number of mutated genes with a negative coefficient in ICR trained elastic net model by MSI status in COAD (left panel) and STAD (right panel). P-values of t-test to compare mean number of mutations per MSI group are indicated in the plot.

**Supplementary Figure 8:** Table to check overlap between tumor intrinsic pathways genes and frequently mutated genes. When a gene (columns) is part of a gene signature (rows), this is indicated by “YES”, if not, it is indicated by “NO”. Genes that have a negative coefficient in trained model are shown in blue, pathways that inversely correlate with ICR (**Figure 2B**) are indicated in blue. Genes that have a positive coefficient in trained model are shown in red, pathways that positively correlate with ICR (**Figure 2B**) are indicated in red.

**Supplementary Figure 9:** Scatterplots of each of the combinations of: 1) ICR scores, 2) proliferation ES, 3) TGF-ß signaling ES, and 4) mutation rate (n = 4452). Pearson’s correlation coefficient and regression line (red) are indicated in the plots.

**Supplementary Figure 10:** Pan-cancer Kaplan Meier curves of ICR groups stratified by both Proliferation High (left panels) and Proliferation Low (right panels) groups (corresponding to classification of shown in **Figure 6A**) and by Mutation rate High (top panels) and Mutation rate Low (bottom panels) based on pan-cancer median mutation rate.

**Supplementary Figure 11:** Multivariate Cox proportional hazards model including ICR classification, proliferation enrichment, TGF-ß signaling enrichment, and tumor mutation rate.

**Supplementary Figure 12:** Survival analysis of ICR Low versus ICR High in pathway enrichment categories across 40 pathway signatures (rows) for each cancer type (columns). HRs (hazard ratios) for death in high enrichment categories (**A**) are compared with HRs in low enrichment categories (**B**). **C.** Differences in prognostic impact of ICR classification between pathway signature enrichment categories for each cancer type. HR of ICR Low vs. ICR High was calculated per category from binary classification of enrichment of oncogenic pathway signatures (rows) within individual cancer types (columns). The delta between HR in the highly enriched group and the HR in the group with low enrichment was calculated for each signature/cancer type combination.

## Supplementary Tables

**Supplementary Table 1:** Association of ICR with OS across 31 cancer types with ICR as a categorical variable ICR Low versus ICR High (first and second column; yellow), and ICR as continuous variable (third and fourth column; blue). HR, hazard ratio for death.

**Supplementary Table 2:** Comparison of mean ES of samples from ICR-disabled cancer types with mean ES of samples from ICR-enabled cancer types for 54 oncogenic pathway gene signatures.

**Supplementary Table 3:** Pan-cancer survival analysis stratified by binary classification based on enrichment of selected oncogenic pathway signatures. HR, hazard ratio for death.

