## Supplementary figures for "Genomic landscape of tumor-host interactions with differential prognostic and predictive connotations" (from another version of this preprint: version 1, 2019-02-12)

FIG S1.1

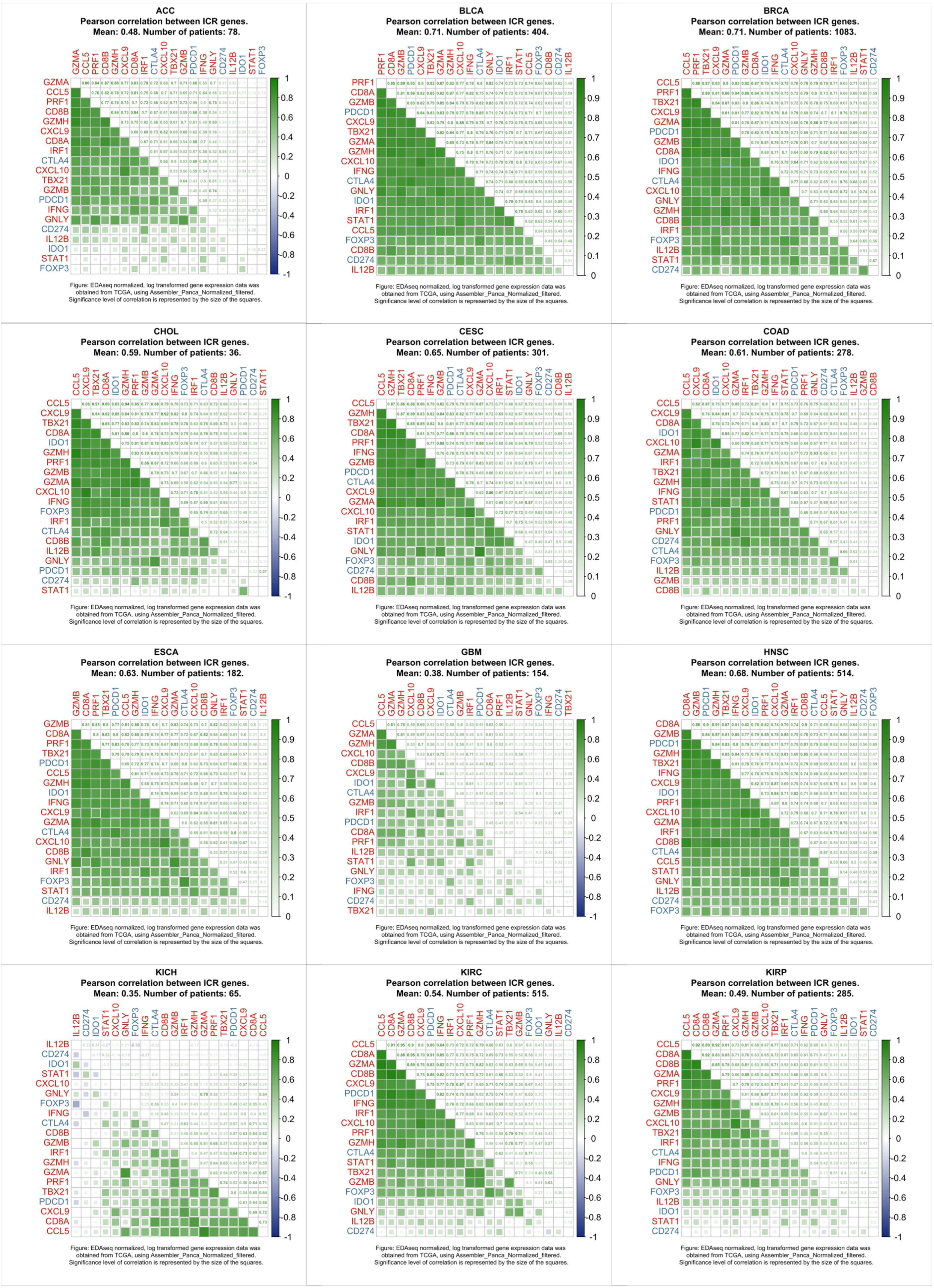
ACC
Pearson correlation between ICR genes.
Mean: 0.48. Number of patients: 78.
Figure: EDAseq normalized, log transformed gene expression data was obtained from TCGA, using Assembler_Panca_Normalized_filtered. Significance level of correlation is represented by the size of the squares.
BLCA
Pearson correlation between ICR genes.
Mean: 0.71. Number of patients: 404.
Figure: EDAseq normalized, log transformed gene expression data was obtained from TCGA, using Assembler_Panca_Normalized_filtered. Significance level of correlation is represented by the size of the squares.
BRCA
Pearson correlation between ICR genes.
Mean: 0.71. Number of patients: 1083.
Figure: EDAseq normalized, log transformed gene expression data was obtained from TCGA, using Assembler_Panca_Normalized_filtered. Significance level of correlation is represented by the size of the squares.
CHOL
Pearson correlation between ICR genes.
Mean: 0.59. Number of patients: 36.
Figure: EDAseq normalized, log transformed gene expression data was obtained from TCGA, using Assembler_Panca_Normalized_filtered. Significance level of correlation is represented by the size of the squares.
CESC
Pearson correlation between ICR genes.
Mean: 0.65. Number of patients: 301.
Figure: EDAseq normalized, log transformed gene expression data was obtained from TCGA, using Assembler_Panca_Normalized_filtered. Significance level of correlation is represented by the size of the squares.
COAD
Pearson correlation between ICR genes.
Mean: 0.61. Number of patients: 278.
Figure: EDAseq normalized, log transformed gene expression data was obtained from TCGA, using Assembler_Panca_Normalized_filtered. Significance level of correlation is represented by the size of the squares.
ESCA
Pearson correlation between ICR genes.
Mean: 0.63. Number of patients: 182.
Figure: EDAseq normalized, log transformed gene expression data was obtained from TCGA, using Assembler_Panca_Normalized_filtered. Significance level of correlation is represented by the size of the squares.
GBM
Pearson correlation between ICR genes.
Mean: 0.38. Number of patients: 154.
Figure: EDAseq normalized, log transformed gene expression data was obtained from TCGA, using Assembler_Panca_Normalized_filtered. Significance level of correlation is represented by the size of the squares.
HNSC
Pearson correlation between ICR genes.
Mean: 0.68. Number of patients: 514.
Figure: EDAseq normalized, log transformed gene expression data was obtained from TCGA, using Assembler_Panca_Normalized_filtered. Significance level of correlation is represented by the size of the squares.
KICH
Pearson correlation between ICR genes.
Mean: 0.35. Number of patients: 65.
Figure: EDAseq normalized, log transformed gene expression data was obtained from TCGA, using Assembler_Panca_Normalized_filtered. Significance level of correlation is represented by the size of the squares.
KIRC
Pearson correlation between ICR genes.
Mean: 0.54. Number of patients: 515.
Figure: EDAseq normalized, log transformed gene expression data was obtained from TCGA, using Assembler_Panca_Normalized_filtered. Significance level of correlation is represented by the size of the squares.
KIRP
Pearson correlation between ICR genes.
Mean: 0.49. Number of patients: 285.
Figure: EDAseq normalized, log transformed gene expression data was obtained from TCGA, using Assembler_Panca_Normalized_filtered. Significance level of correlation is represented by the size of the squares.

FIG S1.2

**LGG**
**Pearson correlation between ICR genes.**
**Mean: 0.4. Number of patients: 514.**

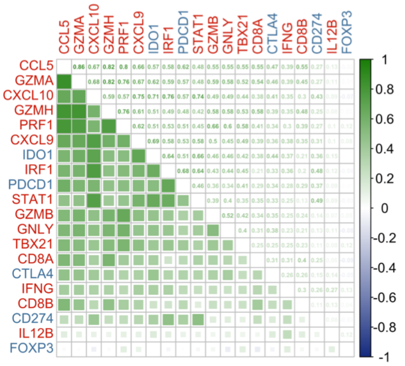

Figure: EDAseq normalized, log transformed gene expression data was obtained from TCGA, using Assembler_Panca_Normalized_filtered. Significance level of correlation is represented by the size of the squares.

**LIHC**
**Pearson correlation between ICR genes.**
**Mean: 0.51. Number of patients: 368.**

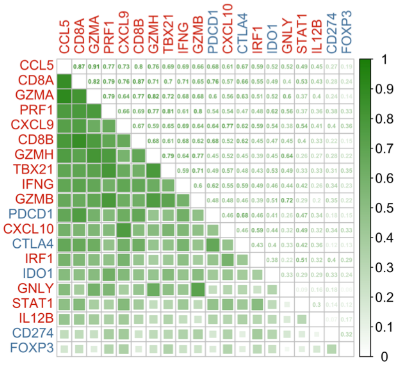

Figure: EDAseq normalized, log transformed gene expression data was obtained from TCGA, using Assembler_Panca_Normalized_filtered. Significance level of correlation is represented by the size of the squares.

**LUAD**
**Pearson correlation between ICR genes.**
**Mean: 0.61. Number of patients: 510.**

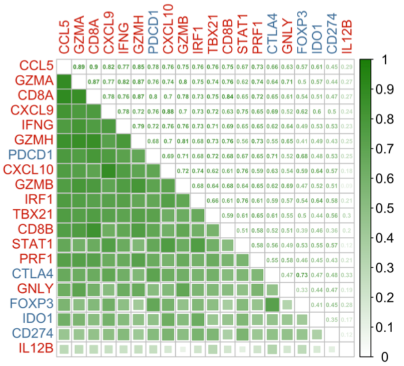

Figure: EDAseq normalized, log transformed gene expression data was obtained from TCGA, using Assembler_Panca_Normalized_filtered. Significance level of correlation is represented by the size of the squares.

**MESO**
**Pearson correlation between ICR genes.**
**Mean: 0.63. Number of patients: 87.**

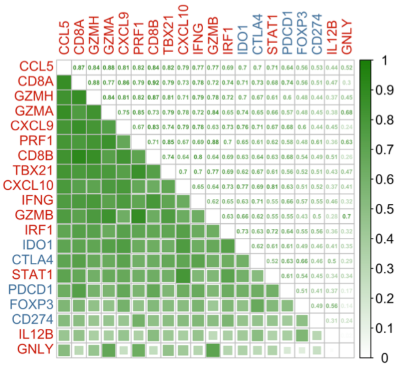

Figure: EDAseq normalized, log transformed gene expression data was obtained from TCGA, using Assembler_Panca_Normalized_filtered. Significance level of correlation is represented by the size of the squares.

**LUSC**
**Pearson correlation between ICR genes.**
**Mean: 0.65. Number of patients: 486.**

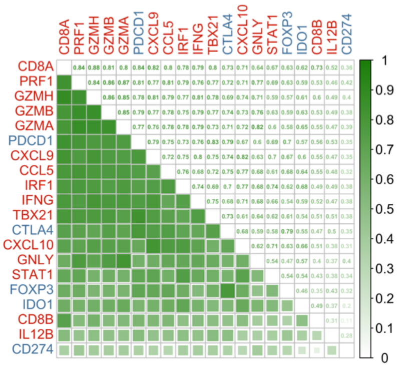

Figure: EDAseq normalized, log transformed gene expression data was obtained from TCGA, using Assembler_Panca_Normalized_filtered. Significance level of correlation is represented by the size of the squares.

**OV**
**Pearson correlation between ICR genes.**
**Mean: 0.65. Number of patients: 302.**

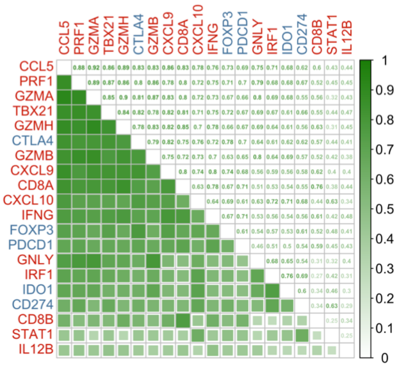

Figure: EDAseq normalized, log transformed gene expression data was obtained from TCGA, using Assembler_Panca_Normalized_filtered. Significance level of correlation is represented by the size of the squares.

**PAAD**
**Pearson correlation between ICR genes.**
**Mean: 0.51. Number of patients: 156.**

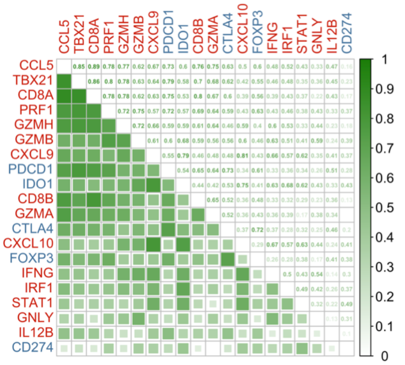

Figure: EDAseq normalized, log transformed gene expression data was obtained from TCGA, using Assembler_Panca_Normalized_filtered. Significance level of correlation is represented by the size of the squares.

**PCPG**
**Pearson correlation between ICR genes.**
**Mean: 0.38. Number of patients: 178.**

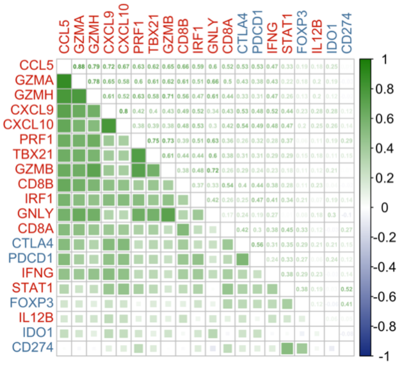

Figure: EDAseq normalized, log transformed gene expression data was obtained from TCGA, using Assembler_Panca_Normalized_filtered. Significance level of correlation is represented by the size of the squares.

**READ**
**Pearson correlation between ICR genes.**
**Mean: 0.5. Number of patients: 90.**

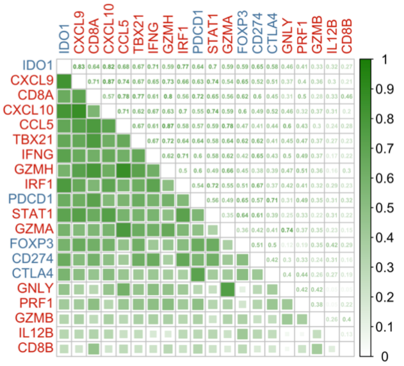

Figure: EDAseq normalized, log transformed gene expression data was obtained from TCGA, using Assembler_Panca_Normalized_filtered. Significance level of correlation is represented by the size of the squares.

**PRAD**
**Pearson correlation between ICR genes.**
**Mean: 0.51. Number of patients: 493.**

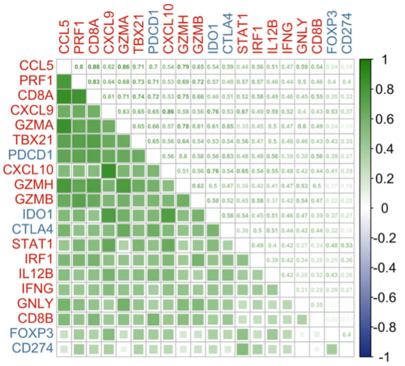

Figure: EDAseq normalized, log transformed gene expression data was obtained from TCGA, using Assembler_Panca_Normalized_filtered. Significance level of correlation is represented by the size of the squares.

**SARC**
**Pearson correlation between ICR genes.**
**Mean: 0.66. Number of patients: 254.**

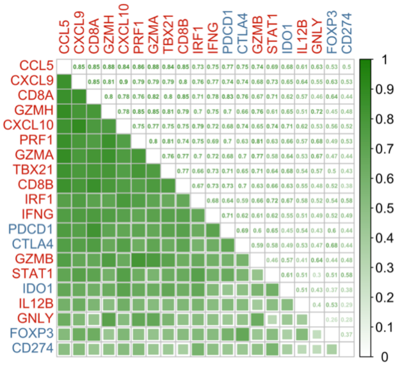

Figure: EDAseq normalized, log transformed gene expression data was obtained from TCGA, using Assembler_Panca_Normalized_filtered. Significance level of correlation is represented by the size of the squares.

**SKCM**
**Pearson correlation between ICR genes.**
**Mean: 0.74. Number of patients: 469.**

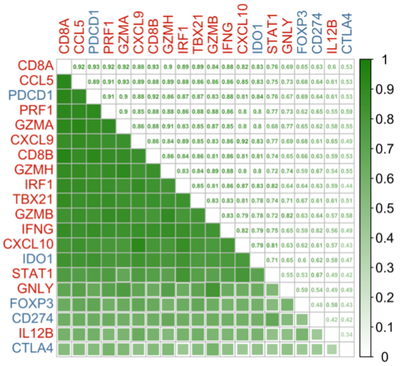

Figure: EDAseq normalized, log transformed gene expression data was obtained from TCGA, using Assembler_Panca_Normalized_filtered. Significance level of correlation is represented by the size of the squares.

### FIG S1.3

**STAD**
**Pearson correlation between ICR genes.**
**Mean: 0.66. Number of patients: 401.**

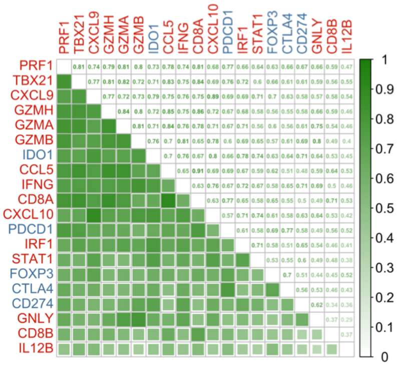


Figure: EDAseq normalized, log transformed gene expression data was obtained from TCGA, using Assembler_Panca_Normalized_filtered. Significance level of correlation is represented by the size of the squares.

**TGCT**
**Pearson correlation between ICR genes.**
**Mean: 0.78. Number of patients: 149.**

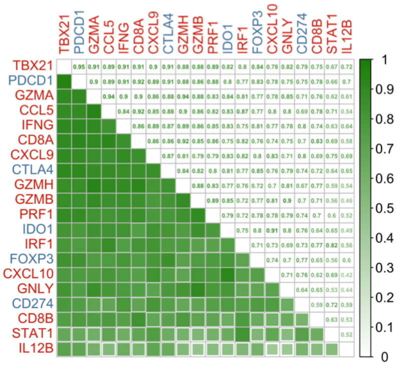


Figure: EDAseq normalized, log transformed gene expression data was obtained from TCGA, using Assembler_Panca_Normalized_filtered. Significance level of correlation is represented by the size of the squares.

**THCA**
**Pearson correlation between ICR genes.**
**Mean: 0.61. Number of patients: 500.**

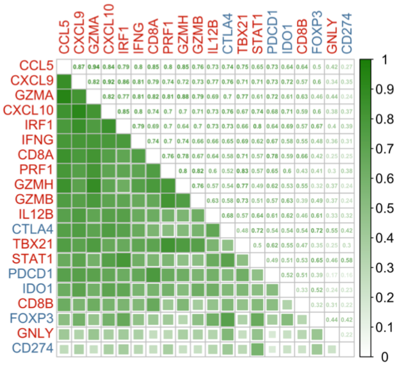


Figure: EDAseq normalized, log transformed gene expression data was obtained from TCGA, using Assembler_Panca_Normalized_filtered. Significance level of correlation is represented by the size of the squares.

**THYM**
**Pearson correlation between ICR genes.**
**Mean: 0.39. Number of patients: 120.**

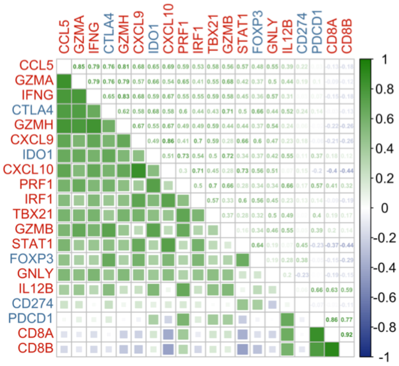


Figure: EDAseq normalized, log transformed gene expression data was obtained from TCGA, using Assembler_Panca_Normalized_filtered. Significance level of correlation is represented by the size of the squares.

**UCEC**
**Pearson correlation between ICR genes.**
**Mean: 0.58. Number of patients: 173.**

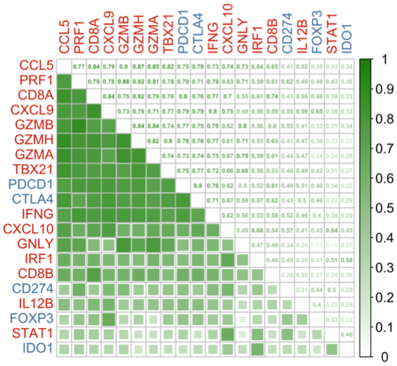


Figure: EDAseq normalized, log transformed gene expression data was obtained from TCGA, using Assembler_Panca_Normalized_filtered. Significance level of correlation is represented by the size of the squares.

**UCS**
**Pearson correlation between ICR genes.**
**Mean: 0.53. Number of patients: 57.**

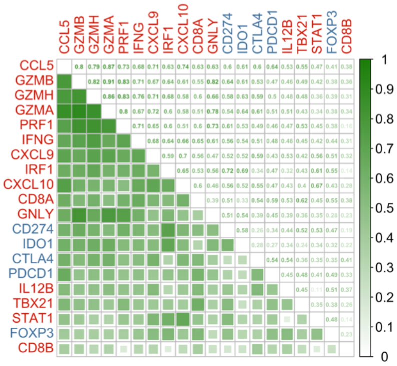


Figure: EDAseq normalized, log transformed gene expression data was obtained from TCGA, using Assembler_Panca_Normalized_filtered. Significance level of correlation is represented by the size of the squares.

**UVM**
**Pearson correlation between ICR genes.**
**Mean: 0.62. Number of patients: 80.**

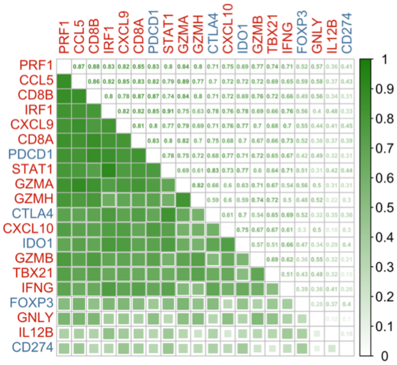


Figure: EDAseq normalized, log transformed gene expression data was obtained from TCGA, using Assembler_Panca_Normalized_filtered. Significance level of correlation is represented by the size of the squares.

FIG S2

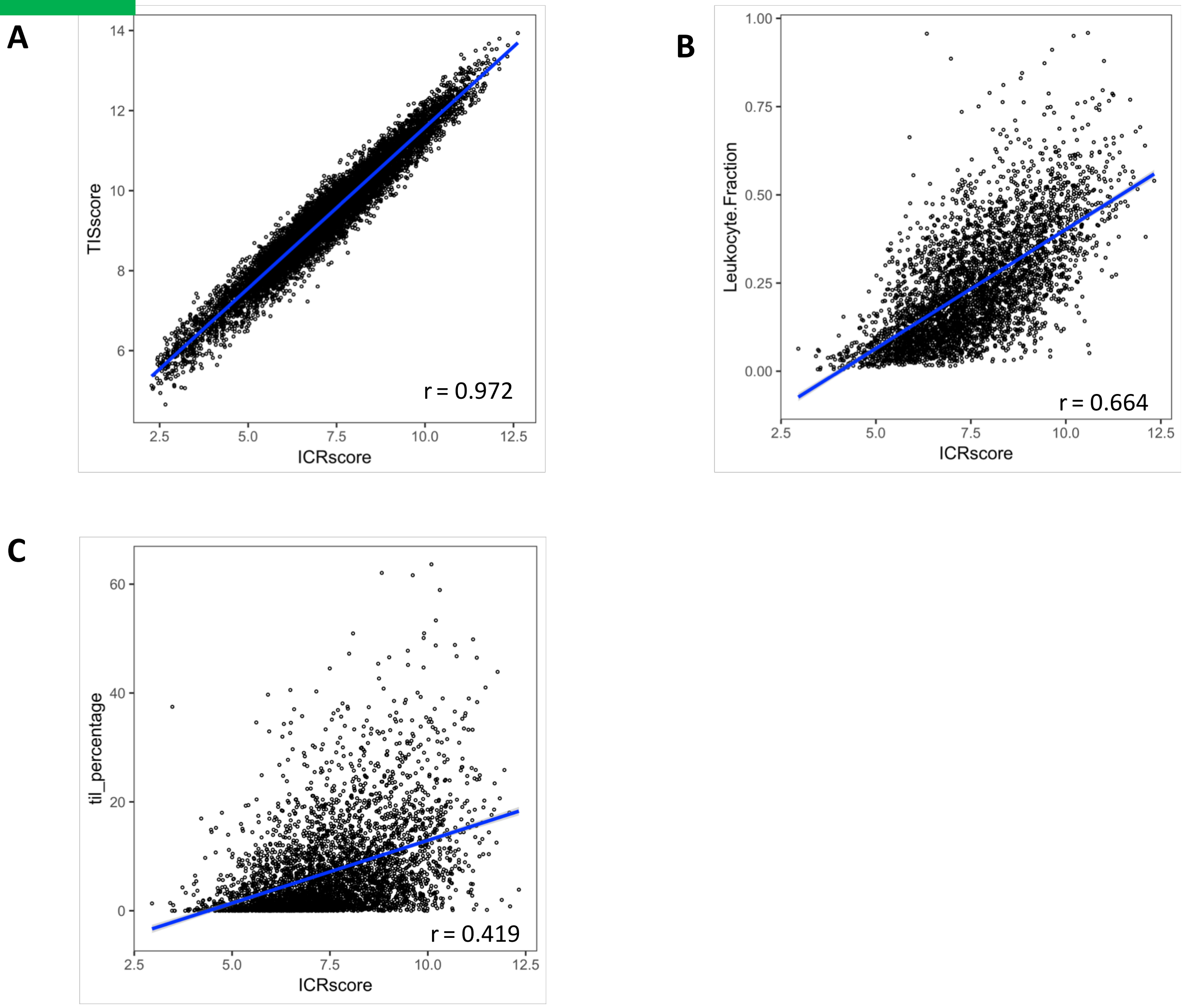

A
TISscore
14
12
10
8
6
2.5
5.0
7.5
10.0
12.5
ICRscore
r = 0.972
B
Leukocyte.Fraction
1.00
0.75
0.50
0.25
0.00
2.5
5.0
7.5
10.0
12.5
ICRscore
r = 0.664
C
til_percentage
60
40
20
0
2.5
5.0
7.5
10.0
12.5
ICRscore
r = 0.419

FIG S3

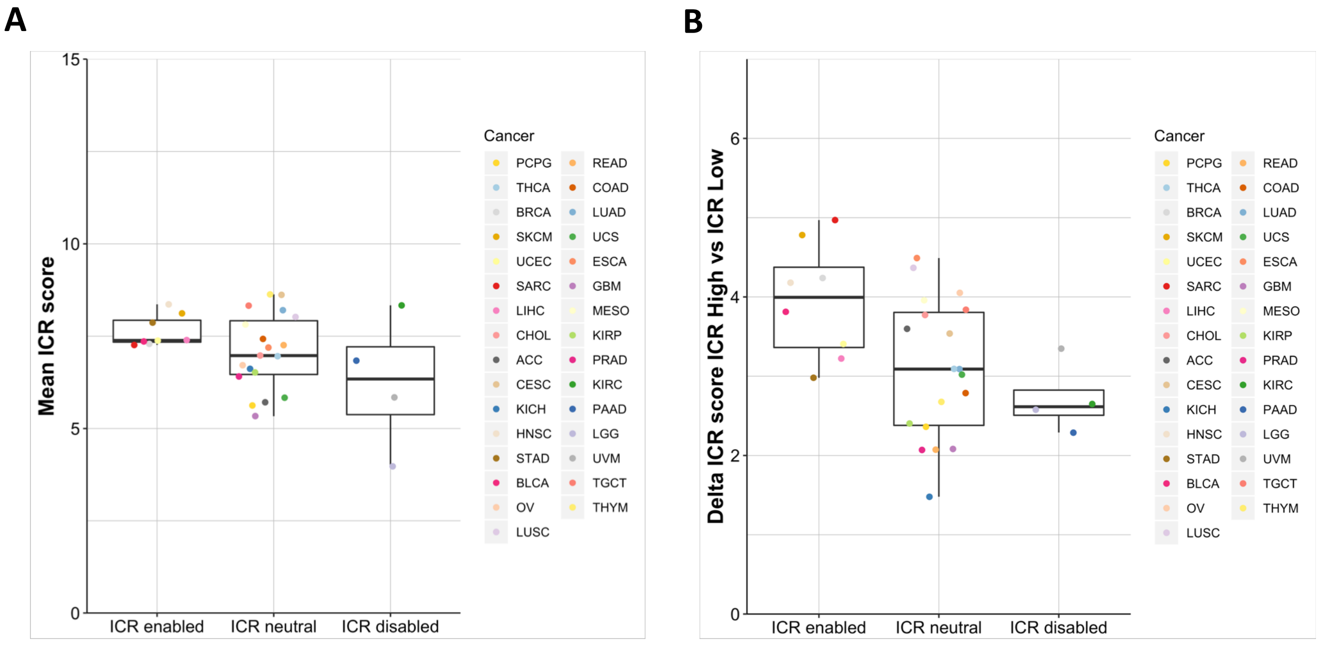
A
Mean ICR score
15
10
5
0
ICR enabled
ICR neutral
ICR disabled
Cancer
PCPG
THCA
BRCA
SKCM
UCEC
SARC
LIHC
CHOL
ACC
CESC
KICH
HNSC
STAD
BLCA
OV
LUSC
READ
COAD
LUAD
UCS
ESCA
GBM
MESO
KIRP
PRAD
KIRC
PAAD
LGG
UVM
TGCT
THYM
B
Delta ICR score ICR High vs ICR Low
6
4
2
0
ICR enabled
ICR neutral
ICR disabled
Cancer
PCPG
THCA
BRCA
SKCM
UCEC
SARC
LIHC
CHOL
ACC
CESC
KICH
HNSC
STAD
BLCA
OV
LUSC
READ
COAD
LUAD
UCS
ESCA
GBM
MESO
KIRP
PRAD
KIRC
PAAD
LGG
UVM
TGCT
THYM

FIG S4

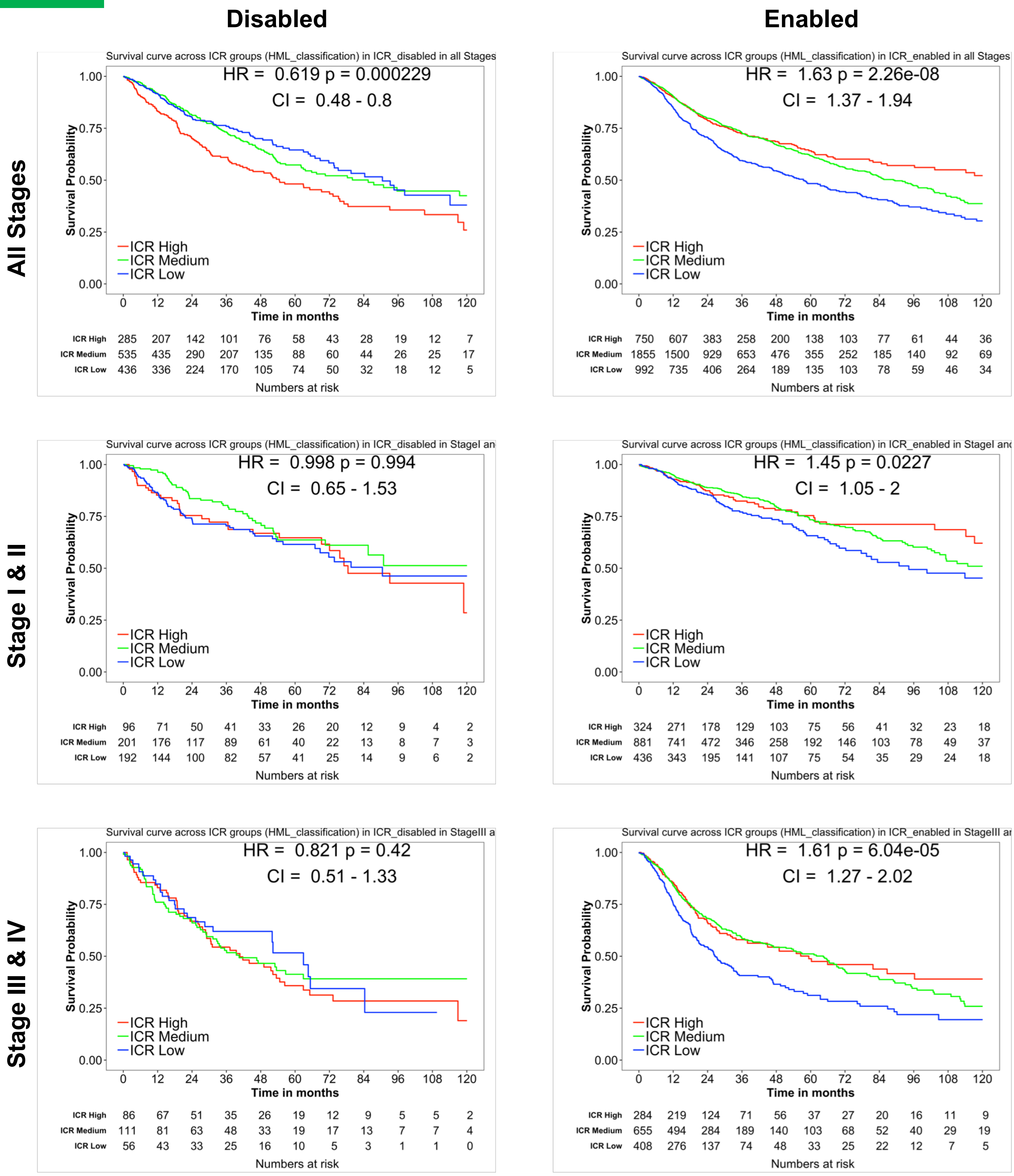
Disabled
Enabled
All Stages
Stage I & II
Stage III & IV
Survival curve across ICR groups (HML_classification) in ICR_disabled in all Stages
HR = 0.619 p = 0.000229
CI = 0.48 - 0.8
Survival curve across ICR groups (HML_classification) in ICR_enabled in all Stages
HR = 1.63 p = 2.26e-08
CI = 1.37 - 1.94
Survival curve across ICR groups (HML_classification) in ICR_disabled in StageI an
HR = 0.998 p = 0.994
CI = 0.65 - 1.53
Survival curve across ICR groups (HML_classification) in ICR_enabled in StageI and
HR = 1.45 p = 0.0227
CI = 1.05 - 2
Survival curve across ICR groups (HML_classification) in ICR_disabled in StageIII a
HR = 0.821 p = 0.42
CI = 0.51 - 1.33
Survival curve across ICR groups (HML_classification) in ICR_enabled in StageIII ar
HR = 1.61 p = 6.04e-05
CI = 1.27 - 2.02
Survival Probability
Time in months
ICR High
ICR Medium
ICR Low
Numbers at risk
ICR High 285 207 142 101 76 58 43 28 19 12 7
ICR Medium 535 435 290 207 135 88 60 44 26 25 17
ICR Low 436 336 224 170 105 74 50 32 18 12 5
ICR High 750 607 383 258 200 138 103 77 61 44 36
ICR Medium 1855 1500 929 653 476 355 252 185 140 92 69
ICR Low 992 735 406 264 189 135 103 78 59 46 34
ICR High 96 71 50 41 33 26 20 12 9 4 2
ICR Medium 201 176 117 89 61 40 22 13 8 7 3
ICR Low 192 144 100 82 57 41 25 14 9 6 2
ICR High 324 271 178 129 103 75 56 41 32 23 18
ICR Medium 881 741 472 346 258 192 146 103 78 49 37
ICR Low 436 343 195 141 107 75 54 35 29 24 18
ICR High 86 67 51 35 26 19 12 9 5 5 2
ICR Medium 111 81 63 48 33 19 17 13 7 7 4
ICR Low 56 43 33 25 16 10 5 3 1 1 0
ICR High 284 219 124 71 56 37 27 20 16 11 9
ICR Medium 655 494 284 189 140 103 68 52 40 29 19
ICR Low 408 276 137 74 48 33 25 22 12 7 5

### FIG S5

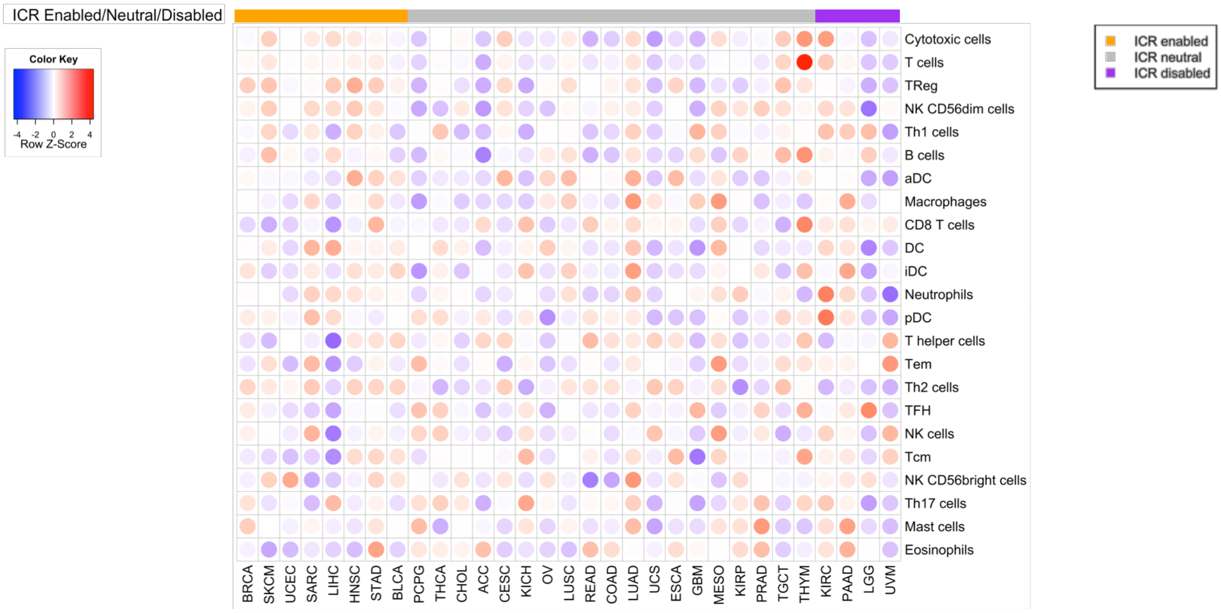
ICR Enabled/Neutral/Disabled
Color Key
-4 -2 0 2 4
Row Z-Score
ICR enabled
ICR neutral
ICR disabled
Cytotoxic cells
T cells
TReg
NK CD56dim cells
Th1 cells
B cells
aDC
Macrophages
CD8 T cells
DC
iDC
Neutrophils
pDC
T helper cells
Tem
Th2 cells
TFH
NK cells
Tcm
NK CD56bright cells
Th17 cells
Mast cells
Eosinophils
BRCA
SKCM
UCEC
SARC
LIHC
HNSC
STAD
BLCA
PCPG
THCA
CHOL
ACC
CESC
KICH
OV
LUSC
READ
COAD
LUAD
UCS
ESCA
GBM
MESO
KIRP
PRAD
TGCT
THYM
KIRC
PAAD
LGG
UVM

FIG S6

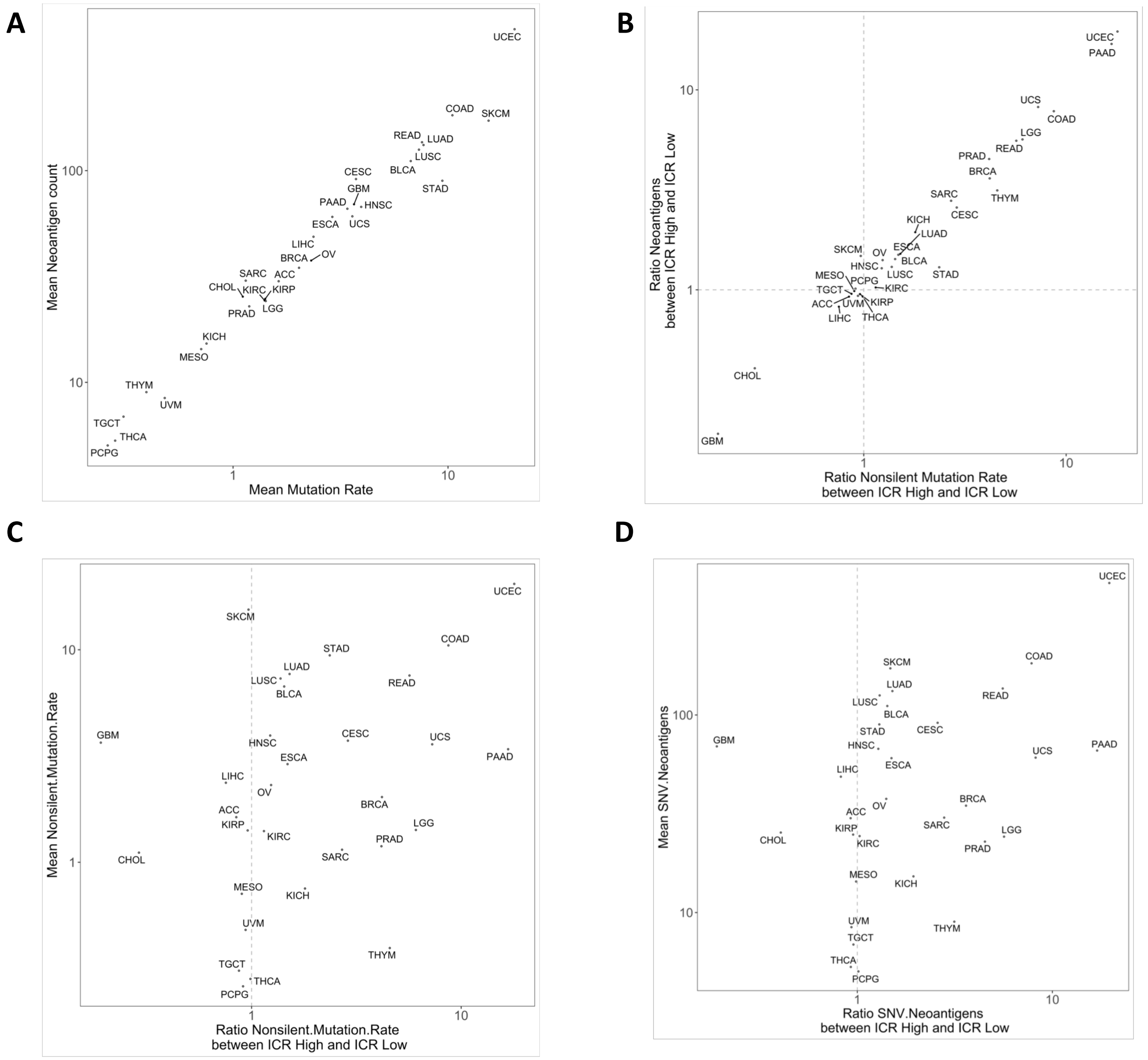
A
UCEC
COAD
SKCM
READ
LUAD
LUSC
BLCA
CESC
GBM
STAD
PAAD
HNSC
ESCA
UCS
LIHC
BRCA
OV
SARC
ACC
CHOL
KIRC
KIRP
PRAD
LGG
KICH
MESO
THYM
UVM
TGCT
THCA
PCPG
100
10
1
10
Mean Neoantigen count
Mean Mutation Rate
B
UCEC
PAAD
UCS
COAD
LGG
PRAD
READ
BRCA
SARC
THYM
CESC
KICH
LUAD
SKCM
OV
ESCA
HNSC
BLCA
LUSC
STAD
MESO
PCPG
TGCT
KIRC
ACC
UVM
KIRP
LIHC
THCA
CHOL
GBM
10
1
1
10
Ratio Neoantigens
between ICR High and ICR Low
Ratio Nonsilent Mutation Rate
between ICR High and ICR Low
C
UCEC
SKCM
COAD
STAD
LUAD
LUSC
READ
BLCA
GBM
CESC
UCS
HNSC
PAAD
ESCA
LIHC
OV
BRCA
ACC
LGG
KIRP
KIRC
CHOL
PRAD
SARC
MESO
KICH
UVM
THYM
TGCT
THCA
PCPG
10
1
1
10
Mean Nonsilent.Mutation.Rate
Ratio Nonsilent.Mutation.Rate
between ICR High and ICR Low
D
UCEC
SKCM
COAD
LUAD
LUSC
READ
BLCA
GBM
STAD
CESC
HNSC
PAAD
UCS
ESCA
LIHC
BRCA
OV
ACC
SARC
KIRP
LGG
CHOL
KIRC
PRAD
MESO
KICH
UVM
THYM
TGCT
THCA
PCPG
100
10
1
10
Mean SNV.Neoantigens
Ratio SNV.Neoantigens
between ICR High and ICR Low

FIG S7

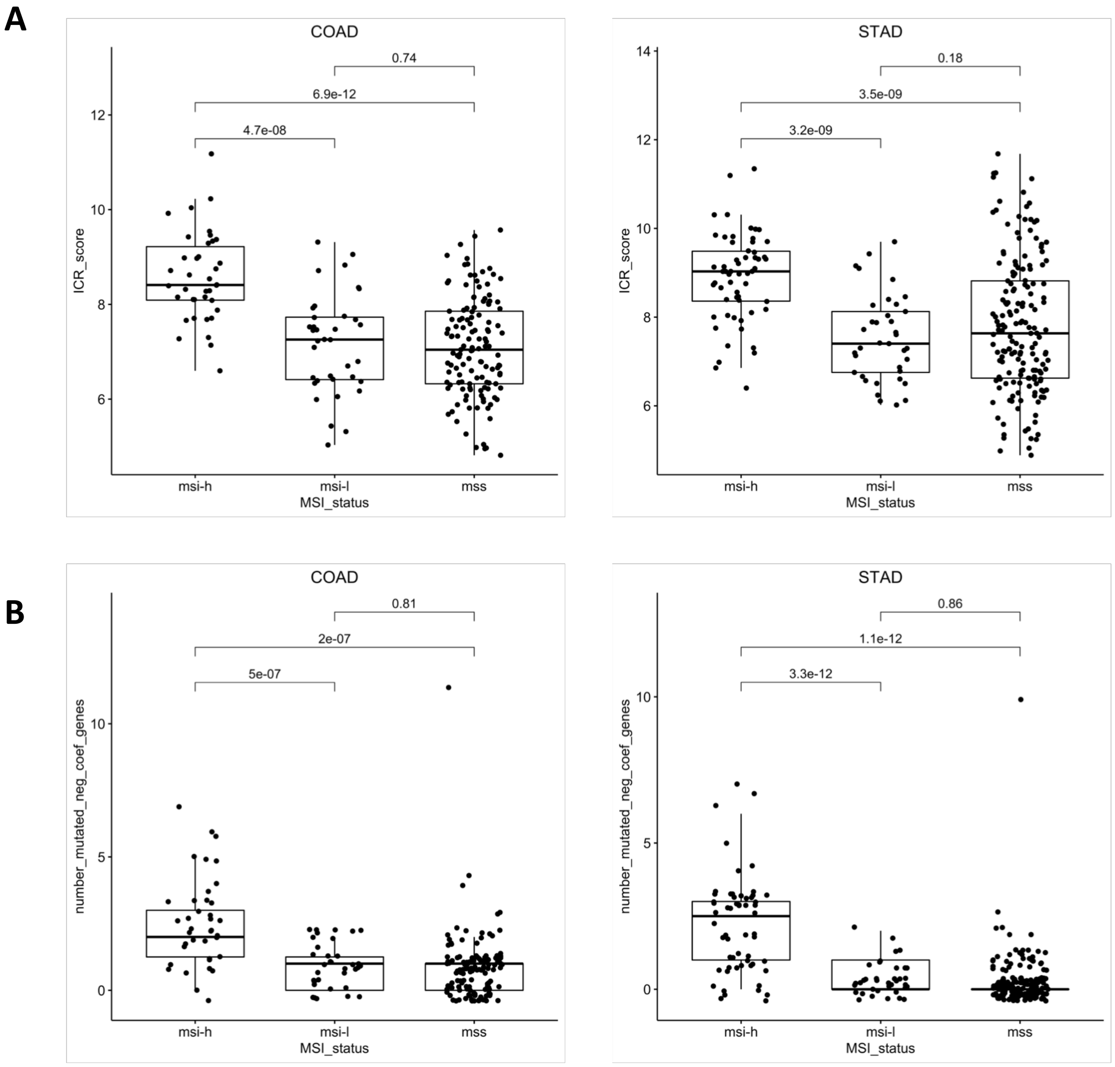

A
COAD
0.74
6.9e-12
4.7e-08
ICR_score
msi-h
msi-l
mss
MSI_status
STAD
0.18
3.5e-09
3.2e-09
B
COAD
0.81
2e-07
5e-07
number_mutated_neg_coef_genes
STAD
0.86
1.1e-12
3.3e-12

### FIG S8

Pathways that inversely correlate with ICR

Pathways that positively correlate with ICR

#### Genes with negative coefficient

| | IDH1 | IDH2 | FOXA2 | NSD1 | PSIP1 | HDAC3 | ZNF814 | MAP3K1 | FRG1 | SOX17 | CARM1 | GATA3 | FKBP5 | FGFR3 | MAT2A | PPP2R5 | MECOM | SMAD2 | MED17 | WNT5A | KRAS | ADAM10 | PRKAR1 | DIS3 | PRRX1 | MFNG | TNPO1 | SPOP | KDM6A | EGFR | IRF7 | NRAS | SUZ12 | RPSAP5 | SF3B1 |
|---|---|---|---|---|---|---|---|---|---|---|---|---|---|---|---|---|---|---|---|---|---|---|---|---|---|---|---|---|---|---|---|---|---|---|---|
| [IPA] AMPK Signaling | NO | NO | NO | NO | NO | NO | NO | YES | NO | NO | NO | NO | NO | NO | NO | NO | NO | NO | NO | NO | NO | NO | NO | NO | NO | NO | NO | NO | NO | YES | NO | NO | NO | NO | NO |
| [IPA] Telomere Extension by Telomer | NO | NO | NO | NO | NO | NO | NO | NO | NO | NO | NO | NO | NO | NO | NO | NO | NO | NO | NO | NO | NO | NO | NO | NO | NO | NO | NO | NO | NO | NO | NO | NO | NO | NO | NO |
| [HM] Hedgehog signaling | NO | NO | NO | NO | NO | NO | NO | NO | NO | NO | NO | NO | NO | NO | NO | NO | NO | NO | NO | NO | NO | NO | NO | NO | NO | NO | NO | NO | NO | NO | NO | NO | NO | NO | NO |
| [TBI] MAPK up genes | NO | NO | NO | NO | NO | NO | NO | NO | NO | NO | NO | NO | NO | NO | NO | NO | NO | NO | NO | NO | NO | NO | NO | NO | NO | NO | NO | NO | NO | NO | NO | NO | NO | NO | NO |
| [IPA] mTOR Signaling | NO | NO | NO | NO | NO | NO | NO | NO | NO | NO | NO | NO | NO | NO | NO | NO | NO | NO | NO | NO | NO | NO | NO | NO | NO | NO | NO | NO | NO | NO | NO | NO | NO | NO | NO |
| [IPA] UVC Induced MAPK Signaling | NO | NO | NO | NO | NO | NO | NO | NO | NO | NO | NO | NO | NO | NO | NO | NO | NO | NO | NO | NO | NO | NO | NO | NO | NO | NO | NO | NO | NO | NO | NO | NO | NO | NO | NO |
| [HM] Wnt beta catenin signaling | NO | NO | NO | NO | NO | NO | NO | NO | NO | NO | NO | NO | NO | NO | NO | NO | NO | NO | NO | NO | NO | NO | NO | NO | NO | NO | NO | NO | NO | NO | NO | NO | NO | NO | NO |
| [HM] Notch signaling | NO | NO | NO | NO | NO | NO | NO | NO | NO | NO | NO | NO | NO | NO | NO | NO | NO | NO | NO | YES | NO | NO | NO | NO | NO | NO | NO | NO | NO | NO | NO | NO | NO | NO | NO |
| [HM] Oxidative phosphorylation | NO | NO | NO | NO | NO | NO | NO | NO | NO | NO | NO | NO | NO | NO | NO | NO | NO | NO | NO | NO | NO | NO | NO | NO | NO | NO | NO | NO | NO | NO | NO | NO | NO | NO | NO |
| [IPA] PTEN Signaling | NO | NO | NO | NO | NO | NO | NO | NO | NO | NO | NO | NO | NO | NO | NO | NO | NO | YES | NO | NO | NO | NO | NO | NO | NO | NO | NO | NO | NO | YES | NO | NO | NO | NO | NO |
| [TBI] Phospholipase | YES | NO | NO | NO | NO | NO | NO | NO | NO | NO | NO | NO | NO | NO | NO | NO | NO | NO | NO | NO | NO | NO | NO | NO | NO | NO | NO | NO | NO | NO | NO | NO | NO | NO | NO |
| [IPA] EGF Signaling | NO | NO | NO | NO | YES | NO | NO | NO | NO | NO | NO | NO | NO | NO | NO | NO | NO | NO | NO | NO | NO | NO | NO | NO | NO | NO | NO | NO | NO | NO | NO | NO | NO | NO | NO |
| [IPA] UVB Induced MAPK Signaling | NO | NO | NO | NO | NO | NO | NO | NO | NO | NO | NO | NO | NO | NO | NO | NO | NO | NO | NO | YES | NO | NO | NO | NO | YES | NO | NO | NO | NO | NO | NO | NO | NO | NO | NO |
| [IPA] PI3K AKT Signaling | YES | YES | NO | NO | NO | NO | NO | NO | NO | NO | NO | NO | NO | NO | NO | NO | NO | NO | NO | NO | NO | NO | NO | NO | NO | NO | NO | NO | NO | NO | NO | NO | NO | NO | NO |
| [IPA] ErbB Signaling | YES | NO | NO | NO | NO | NO | NO | NO | NO | NO | NO | NO | NO | NO | NO | NO | NO | NO | NO | NO | NO | NO | NO | NO | NO | NO | NO | NO | NO | YES | NO | NO | NO | NO | NO |
| [HM] UV response down | NO | NO | NO | NO | NO | NO | NO | NO | NO | NO | NO | NO | NO | NO | NO | NO | NO | NO | NO | NO | NO | NO | NO | NO | NO | NO | NO | NO | NO | NO | NO | NO | NO | NO | NO |
| [HM] UV response up | NO | NO | NO | NO | NO | YES | NO | NO | NO | NO | NO | NO | NO | NO | NO | NO | NO | NO | NO | NO | NO | NO | NO | NO | NO | NO | NO | NO | NO | NO | NO | NO | NO | NO | NO |
| [HM] DNA repair | NO | NO | NO | NO | NO | NO | NO | NO | NO | NO | NO | NO | NO | NO | NO | NO | NO | NO | NO | NO | NO | NO | NO | NO | NO | NO | NO | NO | NO | NO | NO | NO | NO | NO | NO |
| [IPA] ERK5 Signaling | NO | NO | NO | NO | NO | NO | NO | NO | NO | NO | NO | NO | NO | NO | NO | NO | NO | NO | NO | NO | NO | NO | NO | NO | NO | NO | NO | YES | NO | NO | NO | NO | NO | NO | NO |
| [IPA] UVA Induced MAPK Signaling | NO | NO | NO | NO | NO | NO | NO | NO | NO | NO | NO | NO | NO | NO | NO | NO | NO | NO | NO | NO | NO | NO | NO | NO | NO | NO | NO | NO | NO | NO | NO | NO | NO | NO | NO |
| [IPA] Estrogen Dependent Breast Can | NO | NO | NO | NO | NO | NO | NO | YES | NO | NO | NO | NO | NO | NO | NO | NO | NO | NO | NO | NO | NO | NO | NO | NO | YES | NO | NO | NO | NO | NO | NO | NO | NO | NO | NO |
| [IPA] Myc Mediated Apoptosis Signali | NO | NO | NO | NO | NO | NO | NO | NO | NO | NO | NO | NO | NO | YES | NO | NO | NO | NO | NO | NO | NO | NO | NO | NO | NO | NO | NO | NO | NO | NO | NO | NO | NO | NO | NO |
| [HM] Mitotic spindle | NO | NO | NO | NO | NO | NO | NO | NO | NO | NO | NO | NO | NO | NO | NO | NO | NO | NO | NO | NO | NO | NO | NO | NO | NO | NO | NO | NO | NO | NO | NO | NO | NO | NO | NO |
| [IPA] VEGF Signaling | NO | YES | NO | NO | NO | NO | NO | NO | NO | NO | NO | NO | YES | YES | NO | NO | NO | NO | NO | NO | NO | NO | NO | NO | NO | NO | NO | NO | NO | NO | NO | NO | NO | NO | NO |
| [HM] Myc targets | NO | NO | NO | NO | NO | NO | NO | NO | NO | NO | NO | NO | NO | NO | NO | YES | NO | NO | NO | NO | YES | NO | YES | NO | NO | NO | NO | NO | NO | NO | NO | YES | NO | NO | NO |
| [IPA] ERK MAPK Signaling | NO | NO | NO | NO | NO | NO | NO | NO | NO | NO | NO | NO | NO | NO | NO | NO | NO | NO | NO | NO | YES | NO | NO | NO | NO | NO | NO | NO | NO | NO | NO | YES | NO | NO | NO |
| [IPA] ErbB2 ErbB3 Signaling | NO | NO | NO | NO | NO | NO | NO | NO | NO | NO | NO | NO | NO | NO | NO | NO | NO | NO | NO | NO | YES | NO | NO | NO | NO | NO | NO | NO | NO | YES | NO | YES | NO | NO | NO |
| [HM] TGF beta signaling | NO | NO | NO | NO | NO | NO | NO | NO | NO | NO | NO | NO | NO | YES | NO | NO | NO | NO | NO | NO | YES | NO | NO | NO | NO | NO | NO | NO | NO | YES | NO | YES | NO | NO | NO |
| [IPA] HER 2 Signaling in Breast Cancer | NO | NO | NO | NO | NO | NO | NO | NO | NO | NO | NO | NO | NO | NO | NO | YES | NO | NO | NO | NO | YES | NO | NO | NO | NO | NO | NO | NO | NO | NO | NO | YES | NO | NO | NO |
| [HM] PI3K Akt mTOR signaling | NO | NO | NO | NO | NO | NO | NO | NO | NO | NO | NO | NO | NO | NO | NO | NO | NO | NO | NO | NO | YES | NO | NO | NO | NO | NO | NO | NO | NO | YES | NO | YES | NO | NO | NO |
| [HM] KRAS signaling down | NO | NO | NO | NO | NO | NO | NO | NO | NO | NO | NO | NO | NO | NO | NO | YES | NO | NO | NO | NO | YES | NO | NO | NO | NO | NO | NO | NO | NO | NO | NO | YES | NO | NO | NO |
| [IPA] HMGB1 Signaling | NO | NO | NO | NO | NO | NO | NO | NO | NO | NO | NO | NO | NO | NO | NO | YES | NO | NO | NO | NO | NO | NO | YES | NO | NO | NO | NO | NO | NO | NO | NO | NO | NO | NO | NO |
| [HM] mTORC1 signaling | NO | NO | NO | NO | NO | NO | NO | NO | NO | NO | NO | NO | NO | NO | NO | NO | NO | NO | NO | NO | NO | NO | NO | NO | NO | NO | NO | NO | NO | NO | NO | NO | NO | NO | NO |
| [HM] Reactive oxigen species pathwa | NO | NO | NO | NO | NO | NO | NO | NO | NO | NO | NO | NO | NO | NO | NO | NO | NO | NO | NO | NO | YES | NO | NO | NO | NO | NO | NO | NO | NO | NO | NO | YES | NO | NO | NO |
| [HM] Glycolysis | NO | NO | NO | NO | NO | NO | NO | YES | NO | NO | NO | NO | NO | NO | NO | NO | NO | NO | NO | NO | NO | NO | NO | NO | NO | NO | NO | NO | NO | YES | NO | NO | NO | NO | NO |
| [HM] Estrogen response | NO | NO | NO | NO | NO | NO | NO | NO | NO | NO | NO | NO | NO | NO | NO | NO | NO | NO | NO | NO | YES | NO | NO | NO | NO | NO | NO | NO | NO | NO | NO | YES | NO | NO | NO |
| [IPA] Mismatch Repair in Eukaryotes | NO | NO | NO | NO | NO | NO | NO | NO | NO | NO | NO | NO | NO | NO | NO | NO | NO | NO | NO | NO | YES | NO | NO | NO | NO | NO | NO | NO | NO | YES | NO | YES | NO | NO | NO |
| [HM] Hypoxia | NO | NO | NO | NO | NO | NO | NO | YES | NO | NO | NO | NO | NO | NO | NO | NO | NO | NO | NO | NO | NO | NO | NO | NO | NO | NO | NO | NO | NO | NO | NO | NO | NO | NO | NO |
| [TPW] NOS1 Signature | NO | NO | NO | NO | NO | NO | NO | NO | NO | NO | NO | NO | NO | NO | NO | NO | NO | NO | NO | NO | YES | NO | NO | NO | NO | NO | NO | NO | NO | NO | NO | YES | NO | NO | NO |
| [TPW] Hypoxia/Adenosine Immune C | NO | NO | NO | NO | NO | NO | NO | NO | NO | NO | NO | NO | NO | NO | NO | NO | NO | NO | NO | NO | YES | NO | NO | NO | NO | NO | NO | NO | NO | YES | NO | YES | NO | NO | NO |
| [HM] G2M checkpoint | NO | NO | NO | NO | NO | NO | NO | NO | NO | NO | NO | NO | NO | NO | NO | NO | NO | NO | NO | NO | YES | NO | NO | NO | NO | NO | NO | NO | NO | YES | NO | YES | NO | NO | NO |
| [HM] E2F targets | NO | NO | NO | NO | NO | NO | NO | NO | NO | NO | NO | NO | NO | NO | NO | NO | NO | NO | NO | NO | NO | NO | NO | NO | NO | NO | NO | NO | NO | YES | NO | NO | NO | NO | NO |
| [HM] Epithelial mesenchymal transiti | NO | NO | NO | NO | NO | NO | NO | NO | NO | NO | NO | NO | NO | NO | NO | NO | NO | NO | NO | NO | YES | NO | NO | NO | NO | NO | NO | NO | NO | YES | NO | YES | NO | NO | NO |
| [HM] Angiogenesis | NO | NO | NO | NO | NO | NO | NO | NO | NO | NO | NO | NO | NO | NO | NO | NO | NO | NO | NO | NO | NO | NO | NO | NO | NO | NO | NO | NO | NO | NO | NO | NO | NO | NO | NO |
| [TPW] SHC1/pSTAT3 Signature | NO | NO | NO | NO | NO | NO | NO | NO | NO | NO | NO | NO | NO | NO | NO | NO | NO | NO | NO | NO | NO | NO | NO | NO | NO | NO | NO | NO | NO | NO | NO | NO | NO | NO | NO |
| [HM] p53 pathway | NO | NO | NO | NO | NO | NO | NO | NO | NO | NO | NO | NO | NO | NO | NO | NO | NO | NO | NO | NO | NO | NO | NO | NO | NO | NO | NO | NO | NO | NO | NO | NO | NO | NO | NO |
| [TBI] Barrier genes | NO | NO | NO | NO | NO | NO | NO | YES | NO | NO | NO | NO | NO | NO | NO | NO | NO | NO | NO | NO | NO | NO | NO | NO | NO | NO | NO | NO | NO | NO | NO | NO | NO | NO | NO |
| [TPW] PI3Kgamma Signature | NO | NO | NO | NO | NO | NO | NO | NO | NO | NO | NO | NO | NO | NO | NO | NO | NO | NO | NO | NO | NO | NO | NO | NO | NO | NO | NO | NO | NO | NO | NO | NO | NO | NO | NO |
| [LM] Proliferation | NO | NO | NO | NO | NO | NO | NO | NO | NO | NO | NO | NO | NO | NO | NO | NO | NO | NO | NO | NO | NO | NO | NO | NO | NO | NO | NO | NO | NO | NO | NO | NO | NO | NO | NO |
| [HM] KRAS signaling up | NO | NO | NO | NO | NO | NO | NO | NO | NO | NO | NO | NO | NO | NO | NO | NO | NO | NO | NO | NO | NO | NO | NO | NO | NO | NO | NO | NO | NO | NO | NO | NO | NO | NO | NO |
| [HM] Apoptosis | NO | NO | NO | NO | NO | NO | NO | NO | NO | NO | NO | NO | NO | NO | NO | NO | NO | NO | NO | NO | NO | NO | NO | NO | NO | NO | NO | NO | NO | NO | NO | NO | NO | NO | NO |
| [IPA] p38 MAPK Signaling | NO | NO | NO | NO | NO | NO | NO | NO | NO | NO | NO | NO | NO | NO | NO | NO | NO | NO | NO | NO | NO | NO | NO | NO | NO | NO | NO | NO | NO | NO | NO | NO | NO | NO | NO |
| [TPW] Immunogenic Cell Death (ICD) | NO | NO | NO | NO | NO | NO | NO | NO | NO | NO | NO | NO | NO | NO | NO | NO | NO | NO | NO | NO | NO | NO | NO | NO | NO | NO | NO | NO | NO | NO | NO | NO | NO | NO | NO |
| [IPA] TNFR1 Signaling | NO | NO | NO | NO | NO | NO | NO | NO | NO | NO | NO | NO | NO | NO | NO | NO | NO | NO | NO | NO | NO | NO | NO | NO | NO | NO | NO | NO | NO | NO | YES | NO | NO | NO | NO |

#### Genes with positive coefficient

| | CASP8 | B2M | FAS | HLA-B | PIP5K1A | NUP93 | HLA-A | GNAI1 | BRAF | MAP2K1 | ACVR2A | PCSK5 | LPHN2 | SPTAN1 | ARHGEF | ATIC | EPC1 | EP300 | PRKCZ | ARAP3 | PPP2R5 | CEBPA | MYH9 | POM12 | GNA11 | ZNF750 | ERBB3 | CLCC1 | ABL2 | AKAP9 | PIK3CA | MLH1 | PIK3R3 | FIP1L1 | CL |
|---|---|---|---|---|---|---|---|---|---|---|---|---|---|---|---|---|---|---|---|---|---|---|---|---|---|---|---|---|---|---|---|---|---|---|---|
| [IPA] AMPK Signaling | NO | NO | NO | NO | NO | NO | NO | NO | NO | NO | NO | NO | NO | NO | NO | NO | NO | NO | NO | NO | NO | NO | YES | NO | NO | NO | NO | NO | NO | NO | NO | NO | NO | NO | NO |
| [IPA] Telomere Extension by Telomer | NO | NO | NO | NO | NO | NO | NO | NO | NO | NO | NO | NO | NO | YES | YES | NO | NO | NO | NO | YES | NO | NO | YES | NO | NO | NO | NO | NO | NO | NO | NO | NO | NO | NO | NO |
| [HM] Hedgehog signaling | NO | NO | NO | NO | NO | NO | NO | YES | NO | NO | NO | NO | NO | NO | NO | NO | NO | NO | NO | NO | NO | NO | NO | NO | NO | NO | NO | NO | NO | NO | NO | NO | NO | NO | NO |
| [TBI] MAPK up genes | NO | NO | NO | NO | NO | NO | NO | NO | NO | NO | NO | NO | NO | NO | NO | NO | NO | NO | NO | NO | NO | NO | NO | NO | NO | NO | NO | NO | NO | NO | NO | NO | NO | NO | NO |
| [IPA] mTOR Signaling | NO | NO | NO | NO | NO | NO | NO | NO | NO | NO | NO | NO | NO | NO | NO | NO | NO | NO | NO | NO | NO | NO | NO | YES | NO | NO | NO | NO | NO | NO | NO | NO | NO | NO | NO |
| [IPA] UVC Induced MAPK Signaling | NO | NO | NO | NO | NO | NO | NO | NO | NO | NO | NO | NO | NO | NO | NO | NO | NO | NO | NO | NO | NO | NO | NO | NO | NO | NO | NO | NO | NO | NO | NO | NO | NO | NO | NO |
| [HM] Wnt beta catenin signaling | YES | NO | YES | NO | NO | NO | NO | NO | NO | NO | NO | NO | NO | YES | NO | NO | NO | NO | NO | NO | NO | NO | NO | NO | NO | NO | YES | NO | NO | NO | NO | NO | NO | NO | NO |
| [HM] Notch signaling | NO | NO | NO | NO | NO | NO | NO | NO | NO | NO | NO | NO | NO | NO | NO | NO | NO | NO | NO | NO | NO | NO | NO | NO | NO | NO | NO | NO | NO | NO | NO | NO | NO | NO | NO |
| [HM] Oxidative phosphorylation | NO | NO | NO | NO | NO | NO | NO | NO | NO | NO | NO | NO | NO | NO | NO | NO | NO | NO | NO | NO | NO | NO | YES | NO | NO | NO | NO | NO | NO | NO | NO | NO | NO | NO | NO |
| [IPA] PTEN Signaling | NO | NO | NO | NO | NO | NO | NO | NO | NO | NO | NO | NO | NO | NO | NO | NO | NO | NO | NO | NO | NO | NO | NO | NO | NO | NO | NO | NO | NO | NO | NO | NO | YES | NO | NO |
| [TBI] Phospholipase | NO | NO | NO | NO | NO | NO | NO | NO | NO | NO | NO | NO | NO | NO | NO | NO | NO | NO | NO | NO | NO | NO | NO | NO | NO | NO | NO | NO | NO | NO | NO | NO | YES | NO | NO |
| [IPA] EGF Signaling | NO | NO | NO | NO | NO | NO | NO | NO | NO | NO | NO | NO | NO | NO | NO | NO | NO | NO | NO | NO | NO | NO | NO | NO | NO | NO | NO | NO | NO | NO | NO | YES | NO | NO | NO |
| [IPA] UVB Induced MAPK Signaling | NO | NO | YES | NO | NO | NO | NO | NO | NO | NO | NO | NO | NO | NO | NO | NO | NO | NO | NO | NO | NO | NO | NO | NO | NO | NO | NO | NO | NO | NO | NO | NO | NO | NO | NO |
| [IPA] PI3K AKT Signaling | NO | NO | NO | NO | NO | NO | NO | NO | NO | NO | NO | NO | NO | NO | NO | NO | NO | NO | NO | NO | NO | NO | NO | NO | NO | NO | NO | NO | NO | NO | NO | NO | NO | NO | NO |
| [IPA] ErbB Signaling | NO | NO | NO | NO | NO | NO | NO | NO | NO | NO | NO | NO | NO | NO | NO | NO | NO | NO | NO | NO | NO | NO | NO | NO | NO | NO | NO | NO | NO | NO | NO | NO | NO | NO | NO |
| [HM] UV response down | NO | NO | NO | NO | NO | NO | NO | NO | NO | NO | NO | NO | NO | NO | NO | NO | NO | NO | NO | NO | NO | NO | NO | NO | NO | NO | NO | NO | NO | NO | NO | NO | NO | NO | NO |
| [HM] UV response up | NO | NO | YES | NO | NO | NO | NO | NO | NO | NO | NO | NO | NO | NO | NO | NO | NO | NO | NO | NO | NO | YES | NO | YES | NO | NO | NO | NO | NO | NO | NO | NO | NO | NO | NO |
| [HM] DNA repair | NO | NO | NO | NO | NO | NO | NO | NO | NO | NO | NO | NO | NO | NO | NO | NO | NO | NO | NO | NO | NO | NO | NO | NO | NO | NO | NO | NO | NO | NO | NO | NO | NO | NO | NO |
| [IPA] ERK5 Signaling | NO | NO | NO | NO | NO | NO | NO | NO | NO | NO | YES | NO | YES | NO | NO | NO | NO | NO | NO | NO | NO | NO | NO | NO | NO | NO | NO | NO | NO | NO | NO | NO | YES | NO | NO |
| [IPA] UVA Induced MAPK Signaling | NO | NO | NO | NO | NO | NO | NO | NO | NO | NO | NO | NO | NO | NO | NO | NO | NO | NO | NO | NO | NO | NO | NO | NO | NO | NO | NO | NO | NO | NO | NO | NO | NO | NO | NO |
| [IPA] Estrogen Dependent Breast Can | NO | NO | NO | NO | NO | NO | NO | NO | NO | NO | NO | NO | NO | NO | NO | NO | NO | NO | NO | NO | NO | NO | NO | NO | NO | NO | NO | NO | NO | NO | NO | NO | NO | NO | NO |
| [IPA] Myc Mediated Apoptosis Signali | NO | NO | NO | NO | NO | NO | NO | NO | NO | NO | NO | NO | NO | NO | NO | NO | NO | NO | NO | NO | NO | NO | NO | NO | NO | NO | NO | NO | NO | NO | NO | NO | NO | NO | NO |
| [HM] Mitotic spindle | NO | NO | NO | NO | NO | NO | NO | NO | NO | NO | NO | NO | NO | NO | NO | NO | NO | NO | NO | NO | NO | NO | NO | NO | NO | NO | NO | NO | NO | NO | NO | NO | NO | NO | NO |
| [IPA] VEGF Signaling | NO | NO | NO | NO | NO | NO | NO | NO | NO | NO | NO | NO | NO | NO | NO | NO | NO | NO | NO | NO | NO | NO | NO | NO | NO | NO | NO | NO | NO | NO | NO | NO | NO | NO | NO |
| [HM] Myc targets | NO | NO | NO | NO | NO | NO | NO | NO | YES | YES | NO | NO | NO | NO | NO | NO | NO | YES | NO | NO | YES | NO | NO | NO | NO | NO | NO | NO | NO | NO | YES | NO | YES | NO | NO |
| [IPA] ERK MAPK Signaling | NO | NO | NO | NO | NO | NO | NO | NO | NO | YES | NO | NO | NO | NO | NO | NO | NO | NO | NO | NO | NO | NO | NO | NO | NO | NO | NO | NO | NO | NO | YES | NO | YES | NO | NO |
| [IPA] ErbB2 ErbB3 Signaling | NO | NO | NO | NO | NO | NO | NO | NO | NO | YES | NO | NO | NO | NO | NO | NO | NO | NO | YES | NO | NO | NO | NO | NO | NO | NO | YES | NO | NO | NO | YES | NO | YES | NO | NO |
| [HM] TGF beta signaling | NO | NO | NO | NO | NO | NO | NO | NO | NO | YES | NO | NO | NO | NO | NO | NO | NO | NO | YES | NO | NO | NO | NO | NO | NO | NO | NO | NO | NO | NO | YES | NO | YES | NO | NO |
| [IPA] HER 2 Signaling in Breast Cancer | NO | NO | NO | NO | NO | NO | NO | NO | NO | YES | NO | NO | NO | NO | NO | NO | NO | NO | YES | NO | YES | NO | NO | NO | NO | NO | NO | NO | NO | NO | YES | NO | YES | NO | NO |
| [HM] PI3K Akt mTOR signaling | NO | NO | NO | NO | NO | NO | NO | NO | NO | NO | NO | NO | NO | NO | NO | NO | NO | NO | YES | NO | NO | NO | NO | NO | NO | NO | YES | NO | NO | NO | YES | NO | YES | NO | NO |
| [HM] KRAS signaling down | NO | NO | NO | NO | NO | NO | NO | NO | NO | NO | NO | NO | NO | NO | NO | NO | NO | NO | YES | NO | YES | NO | NO | NO | NO | NO | NO | NO | NO | NO | YES | NO | YES | NO | NO |
| [IPA] HMGB1 Signaling | NO | NO | NO | NO | NO | NO | NO | NO | NO | NO | NO | NO | NO | NO | NO | NO | NO | NO | NO | NO | YES | NO | NO | NO | NO | NO | NO | NO | NO | NO | YES | NO | YES | NO | NO |
| [HM] mTORC1 signaling | NO | NO | YES | NO | NO | NO | NO | NO | NO | NO | NO | NO | NO | NO | NO | NO | NO | YES | NO | NO | NO | NO | NO | NO | NO | NO | NO | NO | NO | NO | NO | NO | NO | NO | NO |
| [HM] Reactive oxigen species pathwa | YES | NO | YES | NO | NO | NO | NO | NO | NO | NO | NO | NO | NO | NO | NO | NO | NO | NO | YES | NO | NO | NO | NO | NO | NO | NO | NO | NO | NO | NO | YES | NO | YES | NO | NO |
| [HM] Glycolysis | NO | NO | NO | NO | NO | NO | NO | NO | NO | YES | NO | NO | NO | NO | NO | NO | NO | NO | NO | NO | NO | NO | NO | NO | NO | NO | NO | NO | NO | NO | YES | NO | YES | NO | NO |
| [HM] Estrogen response | NO | NO | NO | NO | NO | NO | NO | NO | NO | YES | NO | NO | NO | NO | NO | NO | NO | NO | NO | NO | NO | NO | NO | NO | NO | NO | NO | NO | NO | NO | YES | NO | YES | NO | NO |
| [IPA] Mismatch Repair in Eukaryotes | NO | NO | NO | NO | NO | NO | NO | NO | NO | NO | NO | NO | NO | NO | NO | NO | NO | YES | NO | NO | NO | NO | NO | NO | NO | NO | NO | NO | NO | NO | YES | NO | YES | NO | NO |
| [HM] Hypoxia | YES | NO | NO | NO | NO | NO | NO | NO | NO | NO | NO | NO | NO | NO | NO | NO | NO | NO | NO | NO | NO | NO | NO | NO | NO | NO | NO | NO | NO | NO | NO | NO | NO | NO | NO |
| [TPW] NOS1 Signature | NO | NO | NO | NO | NO | NO | NO | NO | NO | YES | NO | NO | NO | NO | NO | NO | NO | NO | NO | NO | NO | NO | NO | NO | NO | NO | YES | NO | NO | NO | YES | NO | YES | NO | NO |
| [TPW] Hypoxia/Adenosine Immune C | NO | NO | NO | NO | NO | NO | NO | NO | YES | YES | NO | NO | NO | NO | NO | NO | NO | NO | YES | NO | NO | NO | NO | NO | NO | NO | NO | NO | NO | NO | NO | NO | NO | NO | NO |
| [HM] G2M checkpoint | NO | NO | NO | NO | NO | NO | NO | NO | NO | NO | NO | NO | NO | NO | NO | NO | NO | NO | NO | NO | NO | NO | NO | NO | NO | NO | NO | NO | NO | NO | YES | NO | YES | NO | NO |
| [HM] E2F targets | NO | NO | NO | NO | NO | NO | NO | NO | NO | YES | NO | NO | NO | NO | NO | NO | NO | NO | YES | NO | NO | NO | NO | NO | NO | NO | NO | NO | NO | NO | YES | NO | YES | NO | NO |
| [HM] Epithelial mesenchymal transiti | NO | NO | NO | NO | NO | NO | NO | NO | NO | NO | NO | NO | NO | NO | NO | NO | NO | YES | YES | NO | NO | NO | NO | NO | NO | NO | NO | NO | NO | NO | NO | NO | NO | NO | NO |
| [HM] Angiogenesis | NO | NO | NO | NO | NO | NO | NO | NO | NO | NO | NO | NO | NO | NO | NO | NO | NO | NO | NO | NO | NO | NO | NO | NO | NO | NO | NO | NO | NO | NO | NO | YES | NO | NO | NO |
| [TPW] SHC1/pSTAT3 Signature | NO | NO | NO | NO | NO | NO | NO | NO | NO | NO | NO | NO | NO | NO | NO | NO | NO | NO | NO | NO | NO | NO | NO | NO | NO | NO | NO | NO | NO | NO | NO | NO | NO | NO | NO |
| [HM] p53 pathway | NO | NO | NO | NO | NO | NO | NO | NO | NO | NO | NO | NO | NO | NO | NO | NO | NO | NO | NO | NO | NO | NO | NO | NO | NO | NO | NO | NO | NO | NO | NO | NO | NO | NO | NO |
| [TBI] Barrier genes | NO | NO | NO | NO | NO | NO | NO | NO | NO | NO | NO | NO | NO | NO | NO | NO | NO | NO | NO | NO | NO | NO | NO | NO | NO | NO | NO | NO | NO | NO | NO | NO | NO | NO | NO |
| [TPW] PI3Kgamma Signature | NO | NO | NO | NO | NO | NO | NO | NO | NO | NO | NO | NO | NO | NO | NO | NO | NO | NO | NO | NO | NO | NO | NO | NO | NO | NO | NO | NO | NO | NO | NO | NO | NO | NO | NO |
| [LM] Proliferation | NO | NO | NO | NO | NO | NO | NO | NO | NO | NO | NO | NO | NO | NO | NO | NO | NO | NO | NO | NO | NO | NO | NO | NO | NO | NO | NO | NO | NO | NO | NO | NO | NO | NO | NO |
| [HM] KRAS signaling up | NO | NO | NO | NO | NO | NO | NO | NO | NO | NO | NO | NO | NO | NO | NO | NO | NO | NO | NO | NO | NO | NO | NO | NO | NO | NO | NO | NO | NO | NO | NO | NO | NO | NO | NO |
| [HM] Apoptosis | NO | NO | NO | NO | NO | NO | NO | NO | NO | NO | NO | NO | NO | NO | NO | NO | NO | NO | NO | NO | NO | NO | NO | NO | NO | NO | NO | NO | NO | NO | NO | NO | NO | NO | NO |
| [IPA] p38 MAPK Signaling | NO | NO | NO | NO | NO | NO | NO | NO | NO | NO | NO | NO | NO | NO | NO | NO | NO | NO | NO | NO | NO | NO | NO | NO | NO | NO | NO | NO | NO | NO | NO | NO | NO | NO | NO |
| [TPW] Immunogenic Cell Death (ICD) | NO | NO | NO | NO | NO | NO | NO | NO | NO | NO | NO | NO | NO | NO | NO | NO | NO | NO | NO | NO | NO | NO | NO | NO | NO | NO | NO | NO | NO | NO | NO | NO | NO | NO | NO |
| [IPA] TNFR1 Signaling | NO | NO | NO | NO | NO | NO | NO | NO | NO | NO | NO | NO | NO | NO | NO | NO | NO | NO | NO | NO | NO | NO | NO | NO | NO | NO | NO | NO | NO | NO | NO | NO | NO | NO | NO |

FIG S9

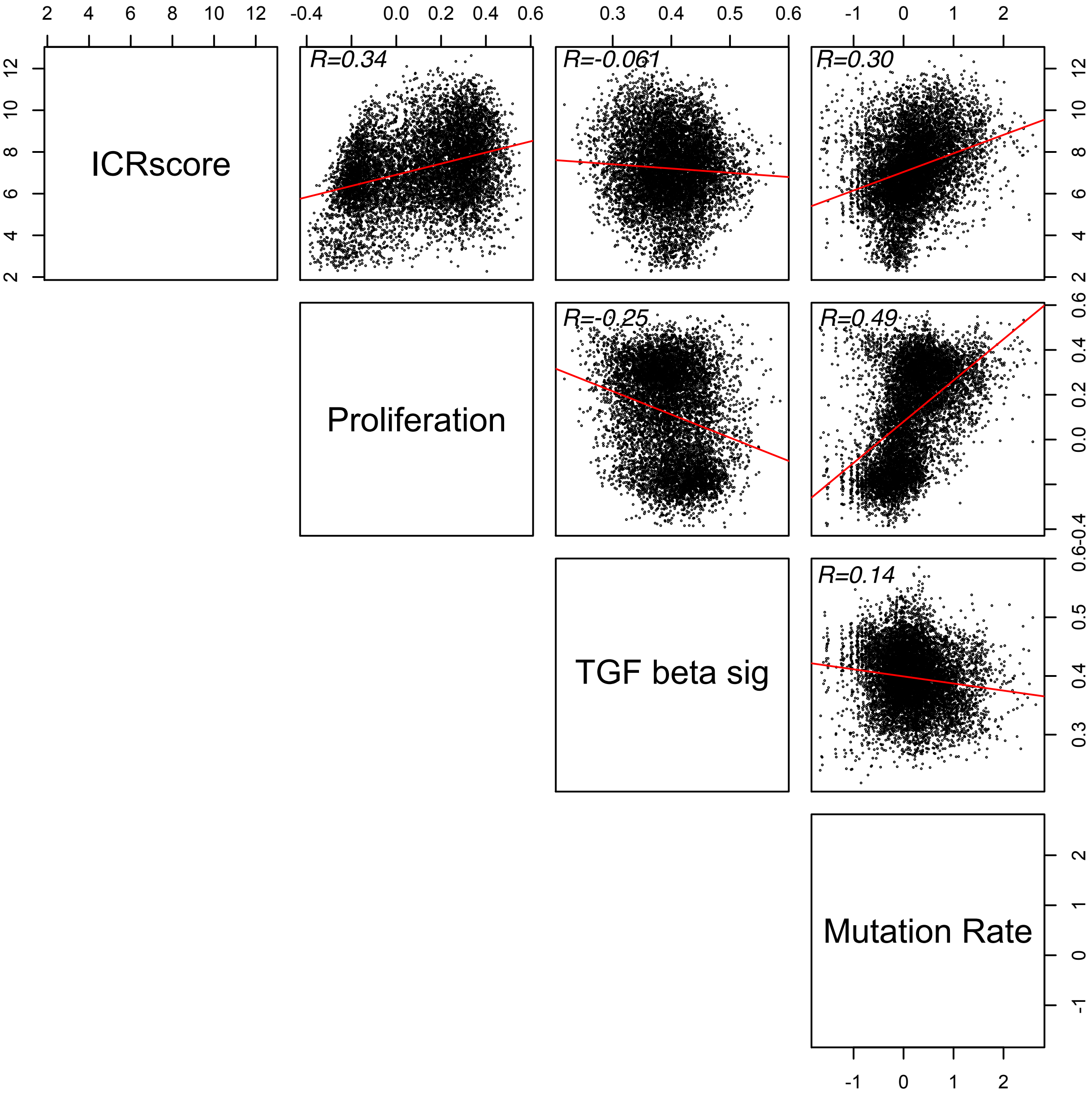

ICRscore
Proliferation
TGF beta sig
Mutation Rate
R=0.34
R=-0.061
R=0.30
R=-0.25
R=0.49
R=0.14

FIG S10

**Proliferation High** | **Proliferation Low**

**Mutation rate High**

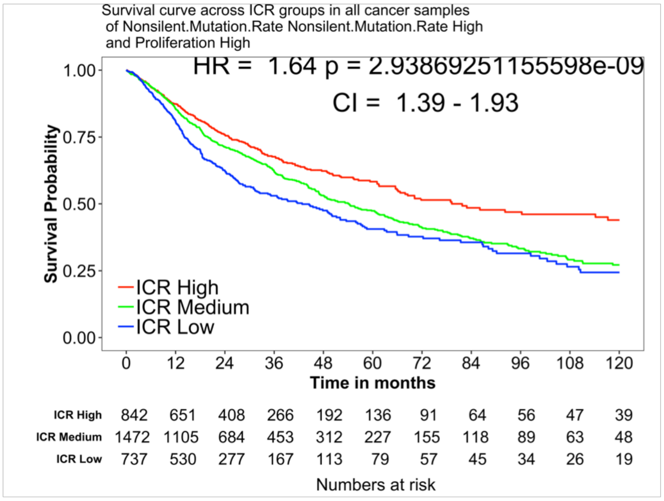


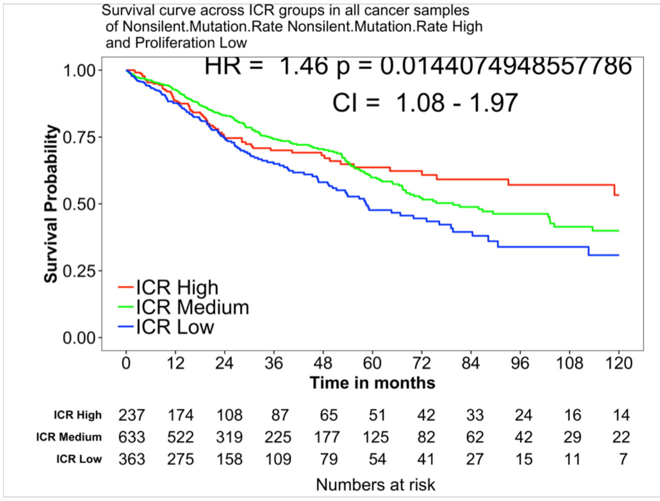


**N = 3051** | **N = 1233**

**Mutation rate Low**

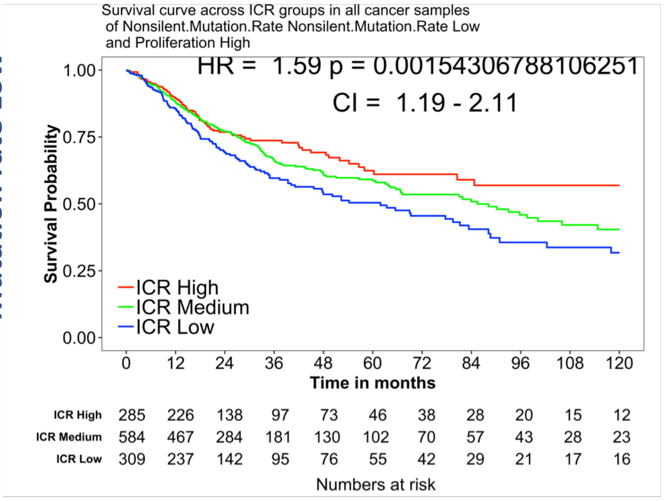


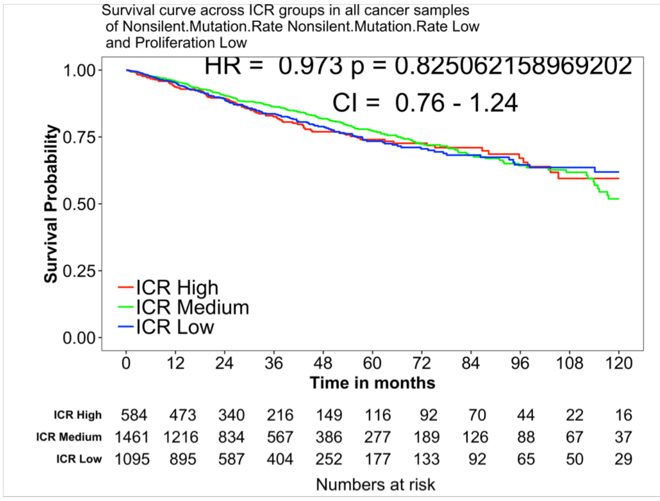


**N = 1178** | **N = 3140**

FIG S11

| Variables | Univariable HR (95% CI) | p | Multivariable HR (95% CI) | p |
|---|---|---|---|---|
| ICR High and Low samples (n = 4452) | | | | |
| ~ ICR cluster (ICR Low vs. High) | 1.244 (1.111-1.392) | 0.000145 *** | 1.430 (1.276- 1.603) | 7.51e-10 *** |
| ~ Proliferation ES (continuous) | 5.818 (4.482-7.554) | <2e-16 *** | 6.643 (5.101-8.651) | < 2e-16 *** |
| ICR High and Low samples (n = 4452) | | | | |
| ~ ICR cluster (ICR Low vs. High) | 1.244 (1.111-1.392) | 0.000145 *** | 1.457 (1.298-1.635) | 1.62e-10 *** |
| ~ Proliferation ES (continuous) | 5.818 (4.482-7.554) | <2e-16 *** | 5.960 (4.486- 7.918) | < 2e-16 *** |
| ~ Mutation rate (continuous) | 1.556 (1.381-1.753) | 3.54e-13 *** | 1.165 (1.015-1.338) | 0.0301 * |
| ICR High and Low samples (n = 4452) | | | | |
| ~ ICR cluster (ICR Low vs. High) | 1.244 (1.111-1.392) | 0.00073 *** | 1.464 (1.304-1.643) | 1.02e-10 *** |
| ~ Proliferation ES (continuous) | 5.818 (4.482-7.554) | <2e-16 *** | 7.929 (5.950-10.566) | < 2e-16 *** |
| ~ Mutation rate (continuous) | 1.556 (1.381-1.753) | 5.05e-09 *** | 1.178 (1.024-1.355) | 0.0217 * |
| ~ TGF beta signaling (continuous) | 21.24 (7.393-61.03) | 1.39e-08 *** | 196.977 (67.610-573.881) | < 2e-16 *** |

FIG S12.1

#### A HR (ICR Low vs. ICR High) in Pathway High groups

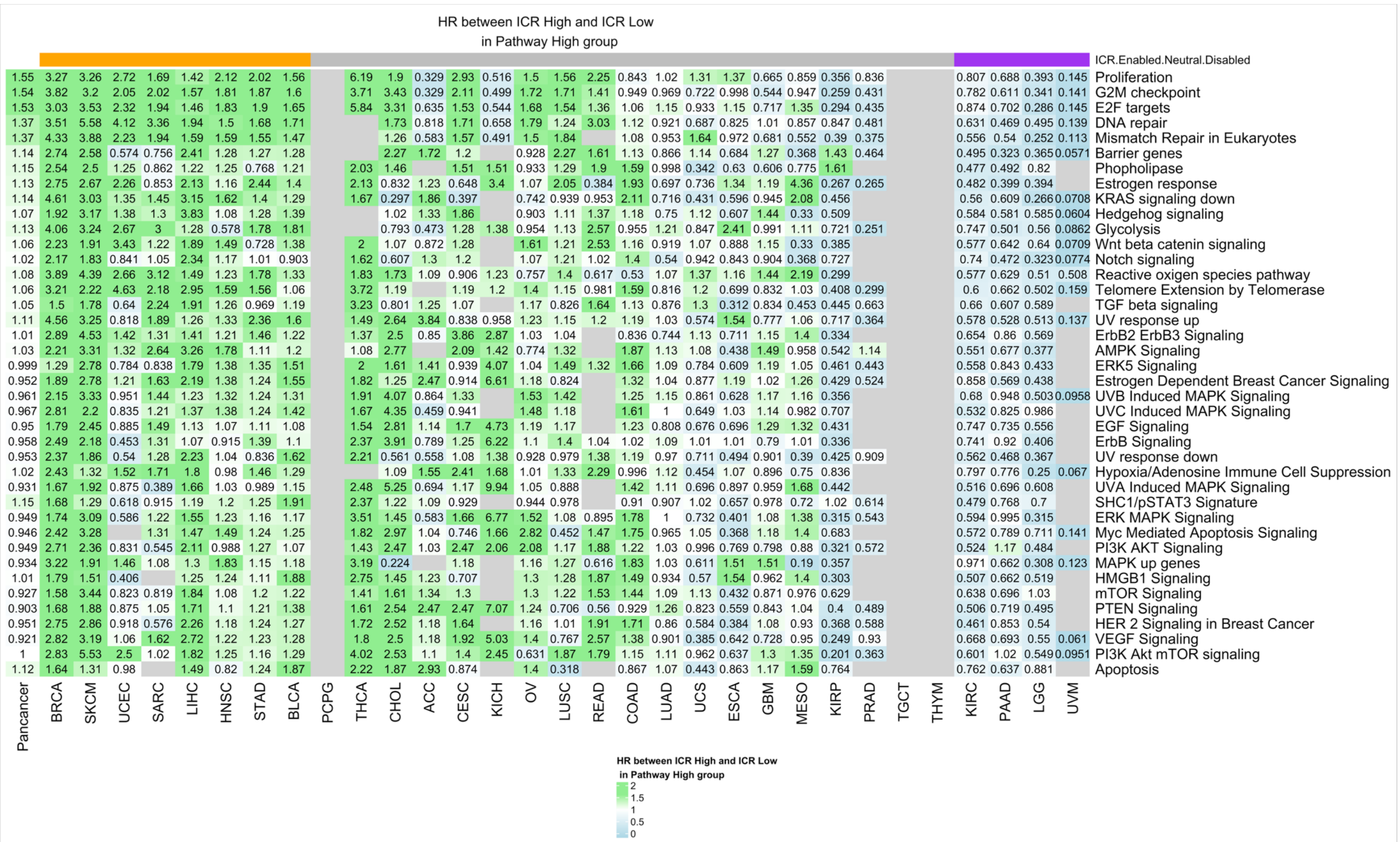


#### B HR (ICR Low vs. ICR High) in Pathway Low groups

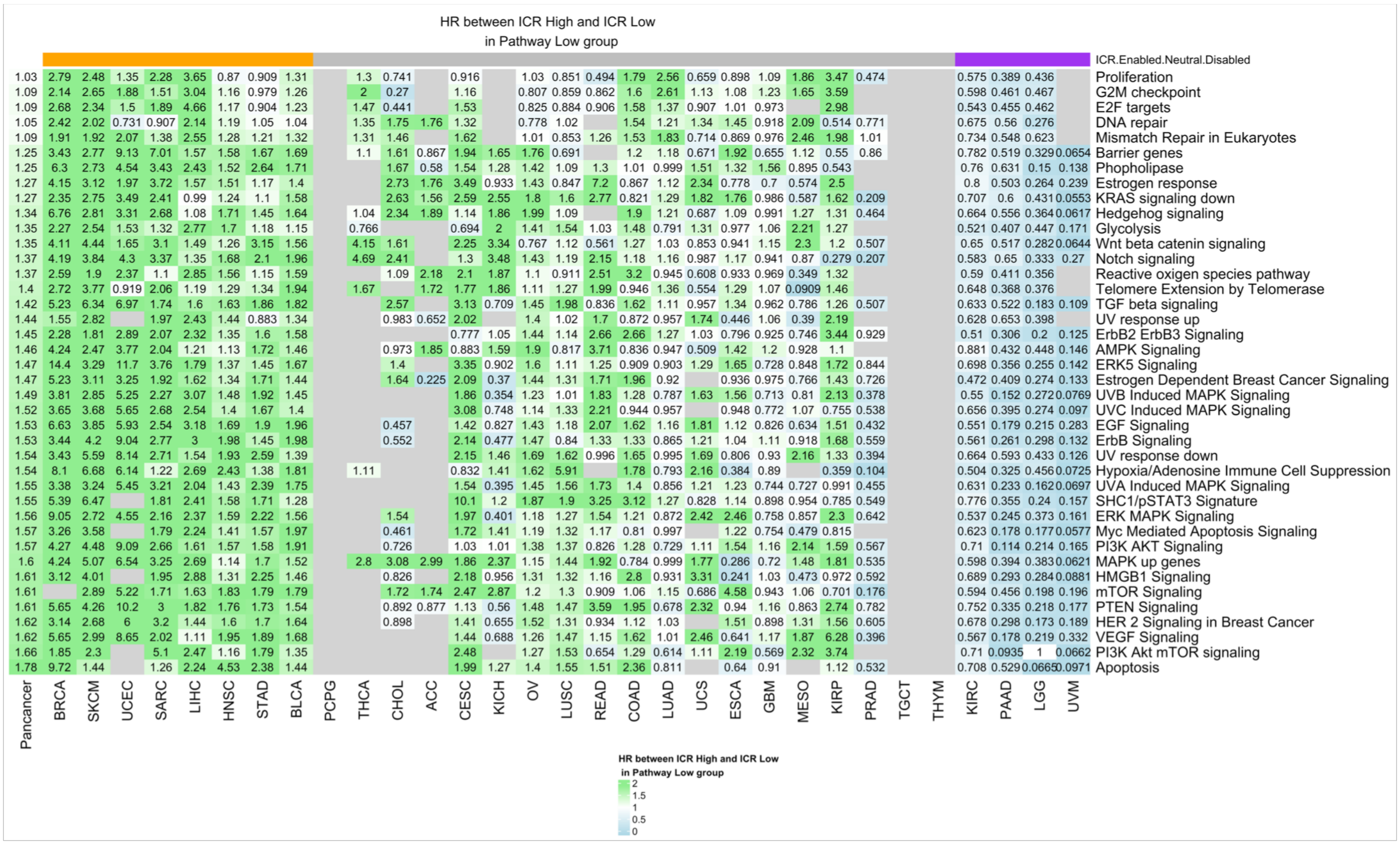

FIG S12.2

#### C Analysis of Delta HR Pathway High – HR Pathway Low

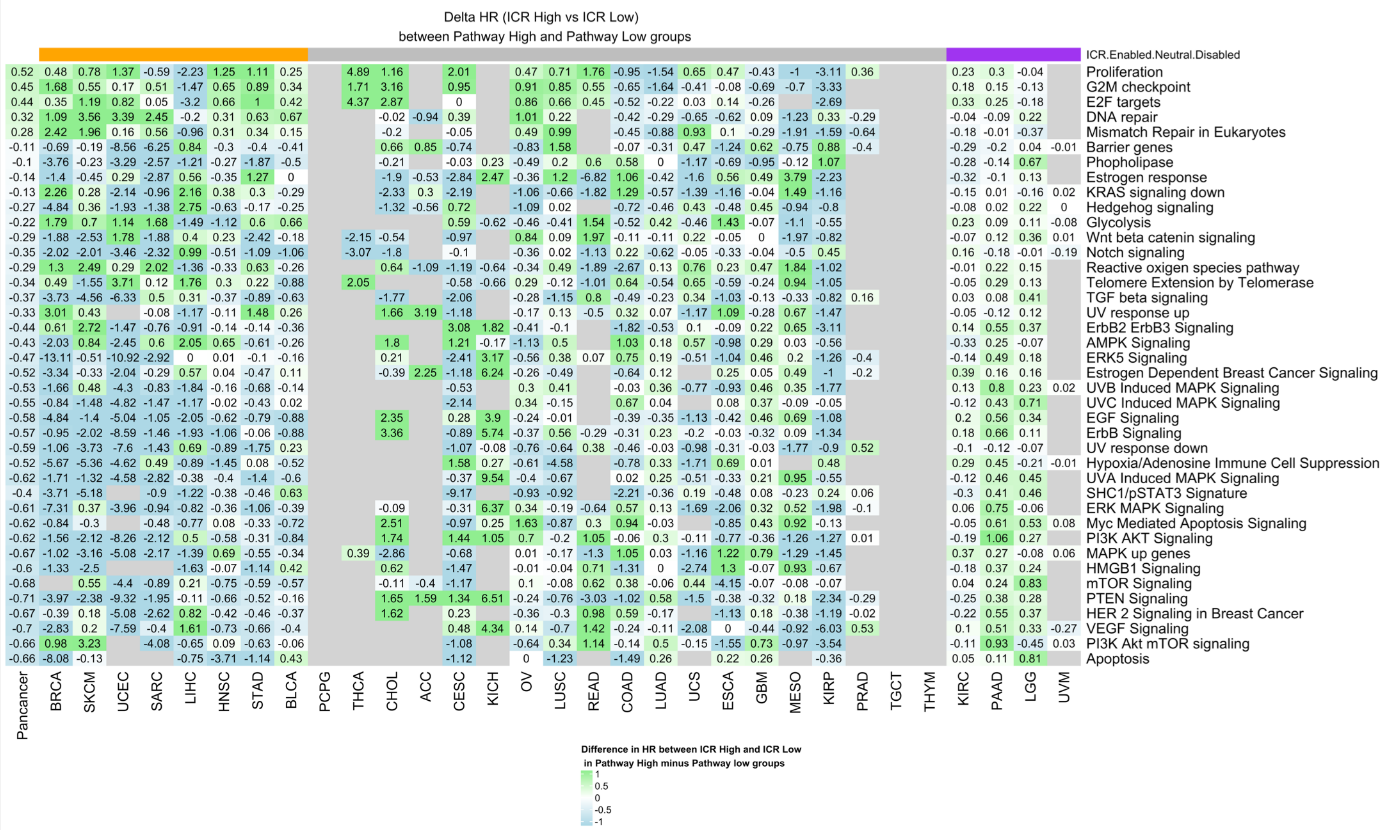

FIG S12
optional

#### D Analysis of Pathway High groups- Only significant HRs

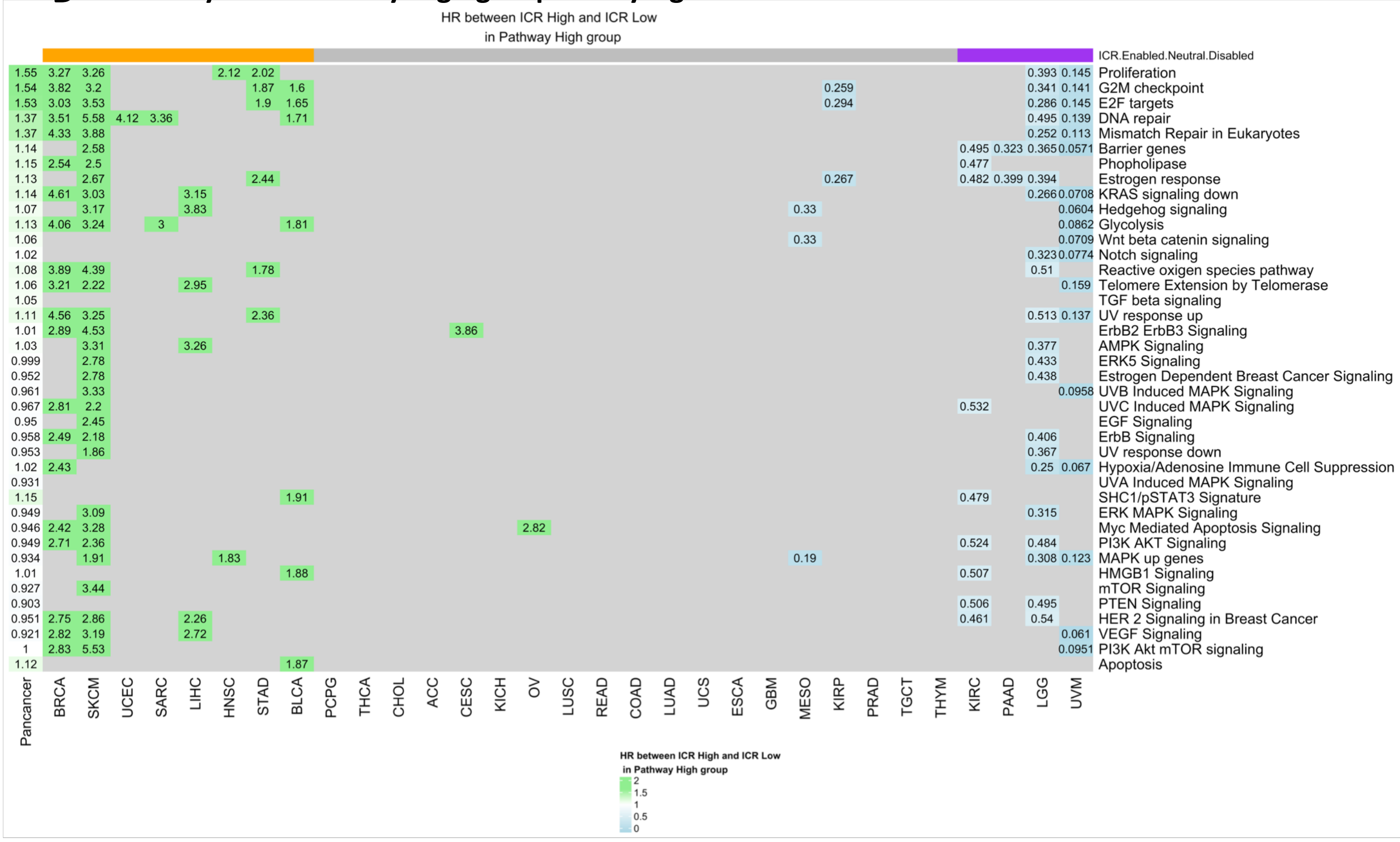


#### E Analysis of Pathway Low groups- Only significant HRs

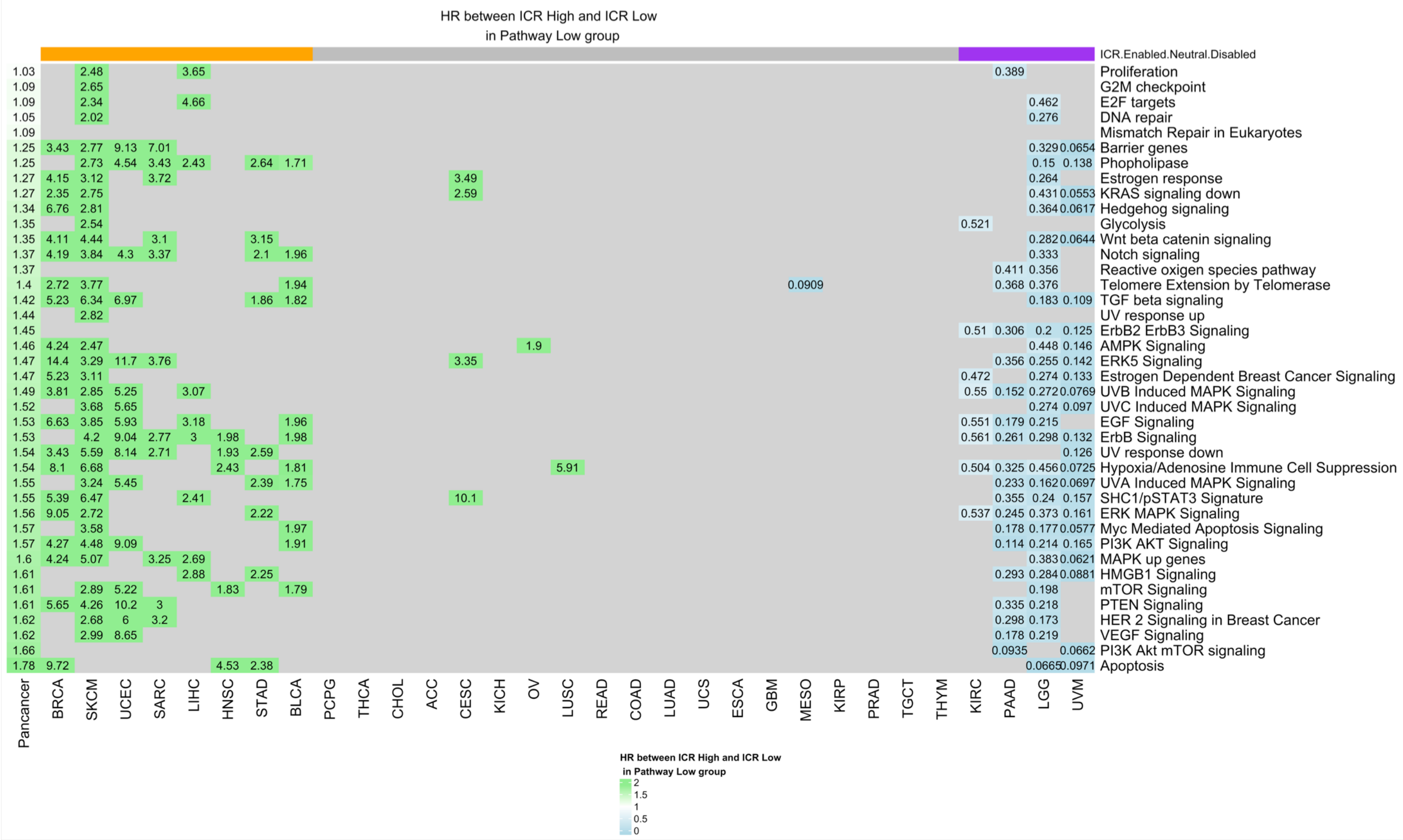

### Supplementary Tables
### (to submit as excel files)

TABLE S1

| Cancer type | N | ICR Low vs. ICR High | | ICR score | |
|---|---|---|---|---|---|
| | | HR | p-value | HR | p-value |
| ACC | 78 | 1.64 | 0.504 | 0.74982546 | 0.11 |
| BLCA | 404 | 1.4 | 0.057 | 0.92726713 | 0.061 |
| BRCA | 1082 | 3.01 | 0.002 | 0.88848161 | 0.033 |
| CESC | 301 | 1.53 | 0.249 | 0.83811938 | 0.022 |
| CHOL | 36 | 1.67 | 0.451 | 0.84823396 | 0.37 |
| COAD | 276 | 1.18 | 0.603 | 0.9231578 | 0.46 |
| ESCA | 182 | 0.929 | 0.873 | 1.075853 | 0.35 |
| GBM | 153 | 0.903 | 0.675 | 1.06974699 | 0.51 |
| HNSC | 514 | 1.43 | 0.095 | 0.89245128 | 0.014 |
| KICH | 65 | 1.51 | 0.575 | 0.71420709 | 0.44 |
| KIRC | 515 | 0.621 | 0.027 | 1.14734061 | 0.051 |
| KIRP | 285 | 0.745 | 0.477 | 1.42999108 | 0.0048 |
| LGG | 513 | 0.353 | 3.29E-05 | 1.55851275 | 3.60E-08 |
| LIHC | 368 | 1.77 | 0.048 | 0.79607172 | 0.01 |
| LUAD | 510 | 0.994 | 0.978 | 0.94336443 | 0.36 |
| LUSC | 486 | 1.22 | 0.503 | 0.96522374 | 0.52 |
| MESO | 87 | 0.856 | 0.700 | 0.91813589 | 0.37 |
| OV | 302 | 1.24 | 0.356 | 0.93242134 | 0.15 |
| PAAD | 156 | 0.542 | 0.038 | 1.07685563 | 0.5 |
| PCPG | 178 | 9.63E+08 | 0.999 | 0.61830021 | 0.42 |
| PRAD | 493 | 0.651 | 0.638 | 0.75338595 | 0.49 |
| READ | 89 | 1.2 | 0.799 | 1.02126437 | 0.94 |
| SARC | 254 | 1.95 | 0.064 | 0.84724121 | 0.0099 |
| SKCM | 469 | 2.89 | 1.20E-05 | 0.81165851 | 1.60E-08 |
| STAD | 401 | 1.41 | 0.099 | 0.95909462 | 0.47 |
| TGCT | 133 | 2.72E-09 | 0.999 | 1.27389557 | 0.44 |
| THCA | 500 | 3.25 | 0.136 | 0.60809675 | 0.041 |
| THYM | 120 | 6.14E-10 | 0.999 | 2.68578687 | 0.0026 |
| UCEC | 173 | 2.25 | 0.095 | 0.79319941 | 0.067 |
| UCS | 57 | 0.969 | 0.946 | 0.97100228 | 0.82 |
| UVM | 80 | 0.0567 | 0.00022 | 1.55280652 | 4.00E-05 |

### TABLE S2

| Pathway | p value | Adj p value (FDR) | Mean ES disabled | Mean ES enabled | Percent increase | Upregulated in | Contribution |
|---|---|---|---|---|---|---|---|
| [LM] Proliferation | 0 | 0 | -0.101502962 | 0.201529573 | -150.3662866 | ICR_enabled | enabling |
| [TBI] Barrier genes | 1.33E-210 | 6.54E-207 | 0.101561488 | 0.326192767 | -68.86458006 | ICR_enabled | enabling |
| [IPA] Mismatch Repair in Eukaryotes | 1.15E-202 | 5.68E-199 | 0.196315537 | 0.26743963 | -26.59444775 | ICR_enabled | enabling |
| [HM] E2F targets | 5.40E-285 | 2.67E-281 | 0.234903815 | 0.314290842 | -25.25909669 | ICR_enabled | enabling |
| [HM] G2M checkpoint | 3.03E-241 | 1.49E-237 | 0.264489805 | 0.331154748 | -20.13105443 | ICR_enabled | enabling |
| [HM] Estrogen response | 1.78E-45 | 8.79E-42 | 0.216199064 | 0.240181921 | -9.985288201 | ICR_enabled | enabling |
| [TPW] NOS1 Signature | 0.035622339 | 1 | -0.189478502 | -0.19687546 | -3.757176088 | ICR_disabled | NA |
| [IPA] p38 MAPK Signaling | 0.001353619 | 1 | 0.136650279 | 0.140091077 | -2.45611538 | ICR_enabled | NA |
| [TPW] PI3Kgamma Signature | 0.46022462 | 1 | 0.186552658 | 0.189398842 | -1.502745999 | ICR_enabled | NA |
| [HM] Epithelial mesenchymal transition | 0.806841286 | 1 | 0.324593367 | 0.325414099 | -0.252211658 | ICR_enabled | NA |
| [IPA] TNFR1 Signaling | 0.817072452 | 1 | 0.273251668 | 0.273577324 | -0.119036457 | ICR_enabled | NA |
| [HM] Myc targets | 7.32E-28 | 3.61E-24 | 0.483914371 | 0.468367558 | 3.319361541 | ICR_disabled | disabling |
| [HM] DNA repair | 1.34E-34 | 6.62E-31 | 0.322702499 | 0.3081982 | 4.706159511 | ICR_disabled | disabling |
| [HM] Angiogenesis | 8.64E-06 | 0.042624944 | 0.30003895 | 0.28492779 | 5.30350529 | ICR_disabled | disabling |
| [HM] mTORC1 signaling | 1.89E-130 | 9.30E-127 | 0.424704896 | 0.402185625 | 5.599223282 | ICR_disabled | disabling |
| [HM] Apoptosis | 1.32E-41 | 6.51E-38 | 0.351122239 | 0.330856488 | 6.125239085 | ICR_disabled | disabling |
| [HM] Glycolysis | 1.37E-40 | 6.75E-37 | 0.303476112 | 0.285473733 | 6.306142056 | ICR_disabled | disabling |
| [HM] Mitotic spindle | 2.18E-110 | 1.08E-106 | 0.381582789 | 0.351624328 | 8.520019358 | ICR_disabled | disabling |
| [HM] p53 pathway | 8.93E-94 | 4.40E-90 | 0.323146371 | 0.297373311 | 8.666904425 | ICR_disabled | disabling |
| [HM] Reactive oxigen species pathway | 1.20E-193 | 5.92E-190 | 0.406450048 | 0.367615832 | 10.56380393 | ICR_disabled | disabling |
| [HM] PI3K Akt mTOR signaling | 0 | 0 | 0.355837625 | 0.316558548 | 12.40815551 | ICR_disabled | disabling |
| [HM] Oxidative phosphorylation | 2.69E-265 | 1.33E-261 | 0.482368517 | 0.42367197 | 13.85424383 | ICR_disabled | disabling |
| [HM] UV response up | 0 | 0 | 0.32317792 | 0.280373634 | 15.26687302 | ICR_disabled | disabling |
| [HM] KRAS signaling down | 1.41E-107 | 6.95E-104 | -0.153148027 | -0.132259584 | 15.79351896 | ICR_enabled | enabling |
| [HM] TGF beta signaling | 5.00E-254 | 2.47E-250 | 0.433467506 | 0.373143115 | 16.16655615 | ICR_disabled | disabling |
| [IPA] ERK MAPK Signaling | 0 | 0 | 0.296571113 | 0.25460962 | 16.48071799 | ICR_disabled | disabling |
| [TPW] SHC1/pSTAT3 Signature | 4.33E-65 | 2.14E-61 | 0.350886019 | 0.300910849 | 16.6079654 | ICR_disabled | disabling |
| [IPA] Myc Mediated Apoptosis Signaling | 0 | 0 | 0.352571235 | 0.298374598 | 18.16395823 | ICR_disabled | disabling |
| [IPA] Estrogen Dependent Breast Cancer Signaling | 0 | 0 | 0.319802592 | 0.27034541 | 18.29407136 | ICR_disabled | disabling |
| [IPA] ErbB2 ErbB3 Signaling | 0 | 0 | 0.369585918 | 0.312122839 | 18.41040524 | ICR_disabled | disabling |
| [IPA] PI3K AKT Signaling | 0 | 0 | 0.399546831 | 0.337241566 | 18.47496607 | ICR_disabled | disabling |
| [IPA] HMGB1 Signaling | 0 | 0 | 0.335852425 | 0.282453833 | 18.90524573 | ICR_disabled | disabling |
| [IPA] VEGF Signaling | 0 | 0 | 0.383494017 | 0.321106768 | 19.42881792 | ICR_disabled | disabling |
| [TPW] Immunogenic Cell Death (ICD) | 1.34E-57 | 6.59E-54 | 0.279761885 | 0.234103093 | 19.50371113 | ICR_disabled | disabling |
| [IPA] EGF Signaling | 0 | 0 | 0.397515236 | 0.33077673 | 20.1763001 | ICR_disabled | disabling |
| [IPA] mTOR Signaling | 0 | 0 | 0.42707648 | 0.353263984 | 20.89443019 | ICR_disabled | disabling |
| [HM] UV response down | 0 | 0 | 0.352923625 | 0.291576254 | 21.03990648 | ICR_disabled | disabling |
| [IPA] ERK5 Signaling | 0 | 0 | 0.356913393 | 0.294692117 | 21.11399429 | ICR_disabled | disabling |
| [IPA] Telomere Extension by Telomerase | 0 | 0 | 0.500118256 | 0.412917561 | 21.11818511 | ICR_disabled | disabling |
| [HM] Hypoxia | 8.11E-190 | 4.00E-186 | 0.332888006 | 0.272460134 | 22.17861078 | ICR_disabled | disabling |
| [IPA] HER 2 Signaling in Breast Cancer | 0 | 0 | 0.324775678 | 0.262364403 | 23.78801156 | ICR_disabled | disabling |
| [IPA] ErbB Signaling | 0 | 0 | 0.296718342 | 0.237525854 | 24.92044016 | ICR_disabled | disabling |
| [IPA] UVA Induced MAPK Signaling | 0 | 0 | 0.30318112 | 0.241680259 | 25.44720077 | ICR_disabled | disabling |
| [IPA] PTEN Signaling | 0 | 0 | 0.377155048 | 0.300084232 | 25.68306094 | ICR_disabled | disabling |
| [HM] Notch signaling | 0 | 0 | 0.340504115 | 0.270225887 | 26.00721512 | ICR_disabled | disabling |
| [IPA] AMPK Signaling | 0 | 0 | 0.253669657 | 0.198475112 | 27.80930253 | ICR_disabled | disabling |
| [IPA] UVC Induced MAPK Signaling | 0 | 0 | 0.327443129 | 0.25438666 | 28.71867148 | ICR_disabled | disabling |
| [HM] Wnt beta catenin signaling | 1.29E-272 | 6.38E-269 | 0.27926302 | 0.216068409 | 29.24750167 | ICR_disabled | disabling |
| [HM] KRAS signaling up | 2.49E-73 | 1.23E-69 | 0.165662676 | 0.12802796 | 29.395701 | ICR_disabled | disabling |
| [IPA] UVB Induced MAPK Signaling | 0 | 0 | 0.29653784 | 0.227544859 | 30.32060646 | ICR_disabled | disabling |
| [TBI] MAPK up genes | 0 | 0 | 0.354955725 | 0.263520522 | 34.69756468 | ICR_disabled | disabling |
| [TPW] Hypoxia/Adenosine Immune Cell Suppression | 1.15E-201 | 5.69E-198 | 0.282834044 | 0.181734296 | 55.63052827 | ICR_disabled | disabling |
| [HM] Hedgehog signaling | 0 | 0 | 0.29185828 | 0.14648611 | 99.23955977 | ICR_disabled | disabling |
| [TBI] Phopholipase | 0 | 0 | 0.157174951 | 0.068744158 | 128.6375398 | ICR_disabled | disabling |

### TABLE S3

| Oncogenic_Pathway | p_value_High | HR_High | CI_lower_High | CI_upper_High | p_value_Low | HR_Low | CI_lower_Low | CI_upper_Low | score_contribution | t.test.fdr.pva |
|---|---|---|---|---|---|---|---|---|---|---|
| [HM] Angiogenesis | 0.015160471 | 1.19 | 1.03 | 1.37 | 0.000583286 | 1.34 | 1.14 | 1.59 | NA | 4.26E-02 |
| [HM] Hypoxia | 0.043490141 | 1.15 | 1 | 1.33 | 4.69E-05 | 1.43 | 1.2 | 1.7 | NA | 4.00E-186 |
| [HM] KRAS signaling up | 5.81E-06 | 1.4 | 1.21 | 1.61 | 0.000432258 | 1.4 | 1.16 | 1.7 | NA | 1.23E-69 |
| [HM] Mitotic spindle | 0.00075894 | 1.26 | 1.1 | 1.44 | 0.037621028 | 1.21 | 1.01 | 1.44 | NA | 1.08E-106 |
| [HM] mTORC1 signaling | 0.000342909 | 1.28 | 1.12 | 1.47 | 0.000587349 | 1.37 | 1.15 | 1.65 | NA | 9.30E-127 |
| [HM] Myc targets | 0.008808615 | 1.2 | 1.05 | 1.38 | 0.004575269 | 1.28 | 1.08 | 1.53 | NA | 3.61E-24 |
| [HM] Oxidative phosphorylation | 0.041989542 | 1.17 | 1.01 | 1.36 | 0.006234439 | 1.24 | 1.06 | 1.44 | NA | 1.33E-261 |
| [HM] p53 pathway | 0.032366297 | 1.17 | 1.01 | 1.35 | 0.000176493 | 1.38 | 1.16 | 1.63 | NA | 4.40E-90 |
| [HM] DNA repair | 2.49E-05 | 1.37 | 1.18 | 1.58 | 0.551731129 | 1.05 | 0.9 | 1.23 | enabling | 6.62E-31 |
| [HM] E2F targets | 3.20E-10 | 1.53 | 1.34 | 1.75 | 0.354086015 | 1.09 | 0.91 | 1.31 | enabling | 2.67E-281 |
| [HM] G2M checkpoint | 1.24E-10 | 1.54 | 1.35 | 1.76 | 0.36823849 | 1.09 | 0.9 | 1.31 | enabling | 1.49E-237 |
| [IPA] Mismatch Repair in Eukaryotes | 6.39E-06 | 1.37 | 1.2 | 1.58 | 0.311865421 | 1.09 | 0.92 | 1.29 | enabling | 5.68E-199 |
| [LM] Proliferation | 1.99E-10 | 1.55 | 1.36 | 1.78 | 0.703483087 | 1.03 | 0.87 | 1.23 | enabling | 0.00E+00 |
| [HM] Apoptosis | 0.115964787 | 1.12 | 0.97 | 1.3 | 2.14E-08 | 1.78 | 1.46 | 2.18 | disabling | 6.51E-38 |
| [HM] Estrogen response | 0.154022515 | 1.13 | 0.96 | 1.32 | 0.000911494 | 1.27 | 1.1 | 1.47 | disabling | 8.79E-42 |
| [HM] Glycolysis | 0.080861609 | 1.13 | 0.99 | 1.3 | 0.000684591 | 1.35 | 1.13 | 1.6 | disabling | 6.75E-37 |
| [HM] Hedgehog signaling | 0.364206051 | 1.07 | 0.93 | 1.23 | 0.000368848 | 1.34 | 1.14 | 1.57 | disabling | 0.00E+00 |
| [HM] KRAS signaling down | 0.123093956 | 1.14 | 0.97 | 1.34 | 0.000812379 | 1.27 | 1.11 | 1.47 | disabling | 6.95E-104 |
| [HM] Notch signaling | 0.785355538 | 1.02 | 0.88 | 1.18 | 0.000116324 | 1.37 | 1.17 | 1.6 | disabling | 0.00E+00 |
| [HM] PI3K Akt mTOR signaling | 0.975613481 | 1 | 0.87 | 1.15 | 2.76E-08 | 1.66 | 1.39 | 1.98 | disabling | 0.00E+00 |
| [HM] Reactive oxigen species pathway | 0.31537027 | 1.08 | 0.93 | 1.25 | 0.000147212 | 1.37 | 1.16 | 1.6 | disabling | 5.92E-190 |
| [HM] TGF beta signaling | 0.500969213 | 1.05 | 0.91 | 1.21 | 2.44E-05 | 1.42 | 1.21 | 1.67 | disabling | 2.47E-250 |
| [HM] UV response down | 0.524574068 | 0.953 | 0.82 | 1.11 | 5.84E-08 | 1.54 | 1.32 | 1.8 | disabling | 0.00E+00 |
| [HM] UV response up | 0.137953972 | 1.11 | 0.97 | 1.27 | 3.30E-05 | 1.44 | 1.21 | 1.72 | disabling | 0.00E+00 |
| [HM] Wnt beta catenin signaling | 0.4632299 | 1.06 | 0.91 | 1.23 | 0.000145971 | 1.35 | 1.16 | 1.58 | disabling | 6.38E-269 |
| [IPA] AMPK Signaling | 0.739147337 | 1.03 | 0.88 | 1.2 | 9.00E-07 | 1.46 | 1.25 | 1.69 | disabling | 0.00E+00 |
| [IPA] EGF Signaling | 0.503347283 | 0.95 | 0.82 | 1.1 | 6.09E-08 | 1.53 | 1.31 | 1.78 | disabling | 0.00E+00 |
| [IPA] ErbB Signaling | 0.570424874 | 0.958 | 0.83 | 1.11 | 1.10E-07 | 1.53 | 1.31 | 1.79 | disabling | 0.00E+00 |
| [IPA] ErbB2 ErbB3 Signaling | 0.858415163 | 1.01 | 0.87 | 1.18 | 7.45E-06 | 1.45 | 1.23 | 1.7 | disabling | 0.00E+00 |
| [IPA] ERK MAPK Signaling | 0.484207728 | 0.949 | 0.82 | 1.1 | 4.37E-08 | 1.56 | 1.33 | 1.83 | disabling | 0.00E+00 |
| [IPA] ERK5 Signaling | 0.984884658 | 0.999 | 0.86 | 1.15 | 2.29E-06 | 1.47 | 1.25 | 1.72 | disabling | 0.00E+00 |
| [IPA] Estrogen Dependent Breast Cancer Signaling | 0.533562949 | 0.952 | 0.81 | 1.11 | 5.63E-07 | 1.47 | 1.26 | 1.71 | disabling | 0.00E+00 |
| [IPA] HER 2 Signaling in Breast Cancer | 0.498376904 | 0.951 | 0.82 | 1.1 | 1.34E-08 | 1.62 | 1.37 | 1.91 | disabling | 0.00E+00 |
| [IPA] HMGB1 Signaling | 0.897438063 | 1.01 | 0.87 | 1.17 | 1.13E-07 | 1.61 | 1.35 | 1.92 | disabling | 0.00E+00 |
| [IPA] mTOR Signaling | 0.314513762 | 0.927 | 0.8 | 1.07 | 2.90E-09 | 1.61 | 1.37 | 1.88 | disabling | 0.00E+00 |
| [IPA] Myc Mediated Apoptosis Signaling | 0.471417808 | 0.946 | 0.81 | 1.1 | 1.01E-07 | 1.57 | 1.33 | 1.85 | disabling | 0.00E+00 |
| [IPA] PI3K AKT Signaling | 0.485708793 | 0.949 | 0.82 | 1.1 | 3.50E-08 | 1.57 | 1.34 | 1.85 | disabling | 0.00E+00 |
| [IPA] PTEN Signaling | 0.186014985 | 0.903 | 0.78 | 1.05 | 1.71E-09 | 1.61 | 1.38 | 1.87 | disabling | 0.00E+00 |
| [IPA] Telomere Extension by Telomerase | 0.471071663 | 1.06 | 0.91 | 1.23 | 1.39E-05 | 1.4 | 1.2 | 1.63 | disabling | 0.00E+00 |
| [IPA] UVA Induced MAPK Signaling | 0.353599437 | 0.931 | 0.8 | 1.08 | 7.26E-08 | 1.55 | 1.32 | 1.81 | disabling | 0.00E+00 |
| [IPA] UVB Induced MAPK Signaling | 0.607306829 | 0.961 | 0.83 | 1.12 | 4.13E-07 | 1.49 | 1.27 | 1.73 | disabling | 0.00E+00 |
| [IPA] UVC Induced MAPK Signaling | 0.657692436 | 0.967 | 0.83 | 1.12 | 1.38E-07 | 1.52 | 1.3 | 1.78 | disabling | 0.00E+00 |
| [IPA] VEGF Signaling | 0.271345717 | 0.921 | 0.79 | 1.07 | 3.25E-09 | 1.62 | 1.38 | 1.9 | disabling | 0.00E+00 |
| [TBI] Barrier genes | 0.090733555 | 1.14 | 0.98 | 1.33 | 0.003090509 | 1.25 | 1.08 | 1.45 | disabling | 6.54E-207 |
| [TBI] MAPK up genes | 0.415776375 | 0.934 | 0.79 | 1.1 | 5.84E-11 | 1.6 | 1.39 | 1.84 | disabling | 0.00E+00 |
| [TBI] Phopholipase | 0.074137822 | 1.15 | 0.99 | 1.34 | 0.003372066 | 1.25 | 1.08 | 1.46 | disabling | 0.00E+00 |
| [TPW] Hypoxia/Adenosine Immune Cell Suppression | 0.754555581 | 1.02 | 0.88 | 1.18 | 3.59E-07 | 1.54 | 1.3 | 1.82 | disabling | 5.69E-198 |
| [TPW] SHC1/pSTAT3 Signature | 0.053781275 | 1.15 | 1 | 1.33 | 1.17E-06 | 1.55 | 1.3 | 1.85 | disabling | 2.14E-61 |
